# A multimodal cell census and atlas of the mammalian primary motor cortex

**DOI:** 10.1101/2020.10.19.343129

**Authors:** BRAIN Initiative Cell Census Network (BICCN), Ricky S. Adkins, Andrew I. Aldridge, Shona Allen, Seth A. Ament, Xu An, Ethan Armand, Giorgio A. Ascoli, Trygve E. Bakken, Anita Bandrowski, Samik Banerjee, Nikolaos Barkas, Anna Bartlett, Helen S. Bateup, M. Margarita Behrens, Philipp Berens, Jim Berg, Matteo Bernabucci, Yves Bernaerts, Darren Bertagnolli, Tommaso Biancalani, Lara Boggeman, A. Sina Booeshaghi, Ian Bowman, Héctor Corrada Bravo, Cathryn René Cadwell, Edward M. Callaway, Benjamin Carlin, Carolyn O'Connor, Robert Carter, Tamara Casper, Rosa G. Castanon, Jesus Ramon Castro, Rebecca K. Chance, Apaala Chatterjee, Huaming Chen, Jerold Chun, Carlo Colantuoni, Jonathan Crabtree, Heather Creasy, Kirsten Crichton, Megan Crow, Florence D. D'Orazi, Tanya L. Daigle, Rachel Dalley, Nick Dee, Kylee Degatano, Benjamin Dichter, Dinh Diep, Liya Ding, Song-Lin Ding, Bertha Dominguez, Hong-Wei Dong, Weixiu Dong, Elizabeth L. Dougherty, Sandrine Dudoit, Joseph R. Ecker, Stephen W. Eichhorn, Rongxin Fang, Victor Felix, Guoping Feng, Zhao Feng, Stephan Fischer, Conor Fitzpatrick, Olivia Fong, Nicholas N. Foster, William Galbavy, James C. Gee, Satrajit S. Ghosh, Michelle Giglio, Thomas H. Gillespie, Jesse Gillis, Melissa Goldman, Jeff Goldy, Hui Gong, Lin Gou, Michael Grauer, Yaroslav O. Halchenko, Julie A. Harris, Leonard Hartmanis, Joshua T. Hatfield, Mike Hawrylycz, Brian Helba, Brian R. Herb, Ronna Hertzano, Houri Hintiryan, Karla E. Hirokawa, Dirk Hockemeyer, Rebecca D. Hodge, Greg Hood, Gregory D. Horwitz, Xiaomeng Hou, Lijuan Hu, Qiwen Hu, Z. Josh Huang, Bingxing Huo, Tony Ito-Cole, Matthew Jacobs, Xueyan Jia, Shengdian Jiang, Tao Jiang, Xiaolong Jiang, Xin Jin, Nikolas L. Jorstad, Brian E. Kalmbach, Jayaram Kancherla, C. Dirk Keene, Kathleen Kelly, Farzaneh Khajouei, Peter V. Kharchenko, Gukhan Kim, Andrew L. Ko, Dmitry Kobak, Kishori Konwar, Daniel J. Kramer, Fenna M. Krienen, Matthew Kroll, Xiuli Kuang, Hsien-Chi Kuo, Blue B. Lake, Rachael Larsen, Kanan Lathia, Sophie Laturnus, Angus Y. Lee, Cheng-Ta Lee, Kuo-Fen Lee, Ed S. Lein, Phil Lesnar, Anan Li, Xiangning Li, Xu Li, Yang Eric Li, Yaoyao Li, Yuanyuan Li, Byungkook Lim, Sten Linnarsson, Christine S. Liu, Hanqing Liu, Lijuan Liu, Jacinta D. Lucero, Chongyuan Luo, Qingming Luo, Evan Z. Macosko, Anup Mahurkar, Maryann E. Martone, Katherine S. Matho, Steven A. McCarroll, Carrie McCracken, Delissa McMillen, Elanine Miranda, Partha P Mitra, Paula Assakura Miyazaki, Judith Mizrachi, Stephanie Mok, Eran A. Mukamel, Shalaka Mulherkar, Naeem M. Nadaf, Maitham Naeemi, Arun Narasimhan, Joseph R. Nery, Lydia Ng, John Ngai, Thuc Nghi Nguyen, Lance Nickel, Philip R. Nicovich, Sheng-Yong Niu, Vasilis Ntranos, Michael Nunn, Dustin Olley, Joshua Orvis, Julia K. Osteen, Pavel Osten, Scott F. Owen, Lior Pachter, Ramesh Palaniswamy, Carter R. Palmer, Yan Pang, Hanchuan Peng, Thanh Pham, Antonio Pinto-Duarte, Nongluk Plongthongkum, Olivier Poirion, Sebastian Preissl, Elizabeth Purdom, Lei Qu, Mohammad Rashid, Nora M. Reed, Aviv Regev, Bing Ren, Miao Ren, Christine Rimorin, Davide Risso, Angeline C. Rivkin, Rodrigo Muñoz-Castañeda, William J. Romanow, Alexander J. Ropelewski, Hector Roux de Bézieux, Zongcai Ruan, Rickard Sandberg, Steven Savoia, Federico Scala, Michael Schor, Elise Shen, Kimberly Siletti, Jared B. Smith, Kimberly Smith, Saroja Somasundaram, Yuanyuan Song, Staci A. Sorensen, David A. Stafford, Kelly Street, Josef Sulc, Susan Sunkin, Valentine Svensson, Pengcheng Tan, Zheng Huan Tan, Bosiljka Tasic, Carol Thompson, Wei Tian, Timothy L. Tickle, Michael Tieu, Jonathan T. Ting, Andreas Savas Tolias, Amy Torkelson, Herman Tung, Eeshit Dhaval Vaishnav, Koen Van den Berge, Cindy T.J. van Velthoven, Charles R. Vanderburg, Matthew B. Veldman, Minh Vu, Wayne Wakeman, Peng Wang, Quanxin Wang, Xinxin Wang, Yimin Wang, Yun Wang, Joshua D. Welch, Owen White, Elora Williams, Fangming Xie, Peng Xie, Feng Xiong, X. William Yang, Anna Marie Yanny, Zizhen Yao, Lulu Yin, Yang Yu, Jing Yuan, Hongkui Zeng, Kun Zhang, Meng Zhang, Zhuzhu Zhang, Sujun Zhao, Xuan Zhao, Jingtian Zhou, Xiaowei Zhuang, Brian Zingg

## Abstract

We report the generation of a multimodal cell census and atlas of the mammalian primary motor cortex (MOp or M1) as the initial product of the BRAIN Initiative Cell Census Network (BICCN). This was achieved by coordinated large-scale analyses of single-cell transcriptomes, chromatin accessibility, DNA methylomes, spatially resolved single-cell transcriptomes, morphological and electrophysiological properties, and cellular resolution input-output mapping, integrated through cross-modal computational analysis. Together, our results advance the collective knowledge and understanding of brain cell type organization: First, our study reveals a unified molecular genetic landscape of cortical cell types that congruently integrates their transcriptome, open chromatin and DNA methylation maps. Second, cross-species analysis achieves a unified taxonomy of transcriptomic types and their hierarchical organization that are conserved from mouse to marmoset and human. Third, cross-modal analysis provides compelling evidence for the epigenomic, transcriptomic, and gene regulatory basis of neuronal phenotypes such as their physiological and anatomical properties, demonstrating the biological validity and genomic underpinning of neuron types and subtypes. Fourth, *in situ* single-cell transcriptomics provides a spatially-resolved cell type atlas of the motor cortex. Fifth, integrated transcriptomic, epigenomic and anatomical analyses reveal the correspondence between neural circuits and transcriptomic cell types. We further present an extensive genetic toolset for targeting and fate mapping glutamatergic projection neuron types toward linking their developmental trajectory to their circuit function. Together, our results establish a unified and mechanistic framework of neuronal cell type organization that integrates multi-layered molecular genetic and spatial information with multi-faceted phenotypic properties.

## INTRODUCTION

Unique among body organs, the human brain is a vast network of information processing units, comprising billions of neurons interconnected through trillions of synapses. Across the brain, diverse neuronal and non-neuronal cells display a wide range of molecular, anatomical, and physiological properties that together shape the network dynamics and computations underlying mental activities and behavior. A remarkable feature of brain networks is their self-assembly through the developmental process, which leverages genomic information shaped by evolution to build a set of stereotyped network scaffolds largely identical among individuals of the same species; life experiences then sculpt neural circuits customized to each individual. An essential step toward understanding the architecture, development, function and neuropsychiatric diseases of the brain is to discover and map its constituent neuronal elements together with the many other cell types that comprise the full organ system.

The notion of “neuron types”, cells with similar properties as the basic units of brain circuits, has been an important concept since the discovery of stereotyped neuronal morphology over a century ago ^1,2^. However, a rigorous and quantitative definition of neuron types has remained surprisingly elusive ^3–7^. Neurons are remarkably complex and heterogeneous, both locally and in their long-range axonal projections that can span the entire brain and connect to many target regions. Many conventional technologies analyze one neuron at a time, and often study only one or two cellular phenotypes in an incomplete way (*e.g.* missing axonal arbors in distant targets). As a result, despite major advances in past decades, until recently phenotypic analyses of neuron types remained severely limited in resolution, robustness, comprehensiveness, and throughput. Besides technical challenges, complexities in the relationship among different cellular phenotypes (multi-modal correspondence) have fueled long-standing debates on how neuron types should be defined ^8^. These debates reflect the lack of a biological framework of cell type organization for understanding brain architecture and function.

In the past decade, single-cell genomics technologies have rapidly swept across many areas of biology including neuroscience, promising to catalyze a transformation from phenotypic description and classification to a mechanistic and explanatory molecular genetic framework for the cellular basis of brain organization ^9–12^. These technologies provide unprecedented resolution and throughput to measure the molecular profiles of individual cells, including the complete sets of actively transcribed genes (the transcriptome) and genome-wide epigenetic landscape (the epigenome). Application of single cell RNA-sequencing (scRNA-seq) to the neocortex, hippocampus, hypothalamus and other brain regions has revealed a complex but tractable hierarchical organization of transcriptomic cell types that are consistent overall with knowledge from decades of anatomical, physiological and developmental studies but with an unmatched level of granularity ^13–19^. Similarly, single-cell DNA methylation and chromatin accessibility studies have begun to reveal cell type-specific genome-wide epigenetic landscapes and gene regulatory networks in the brain ^20–25^. Importantly, the scalability and high information content of these methods allow comprehensive quantitative analysis and classification of cell types, both neuronal and non-neuronal, revealing the molecular basis of cellular phenotypes and properties. Further, these methods are readily applicable to brain tissues across species including humans, providing a quantitative means for comparative analysis that has revealed compelling conservation of cellular architecture as well as specialization of cell types across mammalian species.

Other recent technological advances have crossed key thresholds to provide the resolution and throughput to tackle brain complexity as well, for example for whole-brain neuronal morphology and comprehensive projection mapping ^26,27^. Furthermore, powerful new methods, including imaging-based single-cell transcriptomics, the combination of single-cell transcriptome imaging and functional imaging, and the integration of electrophysiological recording and single-cell sequencing, allow mapping of the spatial organization, function, and electrophysiological, morphological and connectional properties of molecularly defined cell types ^28–32^. Finally, the molecular classification of cell types allows the generation of models for genetic access to specific cell types using transgenic mice and, more recently, short enhancer sequences ^33–39^. All of these methods have been applied to brain tissues in independent studies, but not yet in a coordinated fashion to establish how different modalities correspond with one another, and how explanatory a molecular genetic framework is for other functionally important cellular phenotypes.

Recognizing the unprecedented opportunity to tackle brain complexity brought by these technological advances, the overarching goal of the BRAIN Initiative Cell Census Network (BICCN) is to generate an open-access reference brain cell atlas that integrates molecular, spatial, morphological, connectional, and functional data for describing cell types in mouse, human, and non-human primate brains ^40^. A key concept is the Brain Cell Census, similar conceptually to a population census, which accounts for the population of constituent neuronal and non-neuronal cell types, along with their spatial locations and defining phenotypic characteristics that can be aggregated as cellular populations that make up each brain region. This cell type classification scheme, organized as a taxonomy, should aim for a consensus across modalities and across mammalian species (for conserved types). Beyond the cell census, a Brain Cell Atlas would be embedded in a 3D Common Coordinate Framework (CCF) of the brain ^41^, in which the precise location and distribution of all cell types and their multi-modal features are registered and displayed. Such a cell-type resolution spatial framework will greatly facilitate integration, interpretation and navigation of various types of information for understanding brain network organization and function.

Here we present the cell census and atlas of cell types in one region of the mammalian brain, the primary motor cortex (MOp or M1) of mouse, marmoset and human, through an analysis with unprecedented scope, depth and range of approaches (**Fig. 1**, **Table 1**). MOp is important in the control of complex movement and is well conserved across species. Decades of accumulated anatomical, physiological, and functional studies have provided a rich knowledge base for the integration and interpretation of cell type information in MOp ^42,43^. This manuscript describes a synthesis of results and findings derived from eleven core companion papers through a multi-laboratory coordinated data generation within BICCN. We derive a cross-species consensus transcriptomic taxonomy of cell types and identify conserved and divergent gene expression and epigenomic regulatory signatures from a large and comprehensive set of single-cell/nucleus RNA-sequencing, DNA methylation and chromatin accessibility data. Focusing on mouse MOp, we map the spatial organization of transcriptomic cell types by multiplexed error-robust fluorescence in situ hybridization (MERFISH) and their laminar, morphological and electrophysiological properties by Patch-seq; we report the cell-type resolution input-output wiring diagram of this region by anterograde and retrograde tracing and investigate how axon projection patterns of glutamatergic excitatory neurons correlate with molecularly-defined cell types by Epi-Retro-Seq, Retro-MERFISH (the combination of MERFISH and retrograde labeling), and single-neuron full morphology reconstruction; we describe transgenic driver lines systematically targeting glutamatergic cell types based on marker genes and lineages. Finally, we integrate this vastly diverse array of information into a cohesive depiction of cell types in the MOp region with correlated molecular genetic, spatial, morphological, connectional, and physiological properties and relating them to traditionally described cell types. Such integration is illustrated in detail in example cell types with unique features in MOp: the layer 4 intratelencephalic-projecting (L4 IT) cells and layer 5 extratelencephalic-projecting (L5 ET) cells. This multitude of datasets are organized by the BRAIN Cell Data Center (BCDC) and made public through the BICCN web portal www.biccn.org. Key concepts and terms are described in **Table 2**, including anatomical terms for input and output brain regions for MOp, and hierarchical cell class, subclass and type definitions.

**Figure 1.**
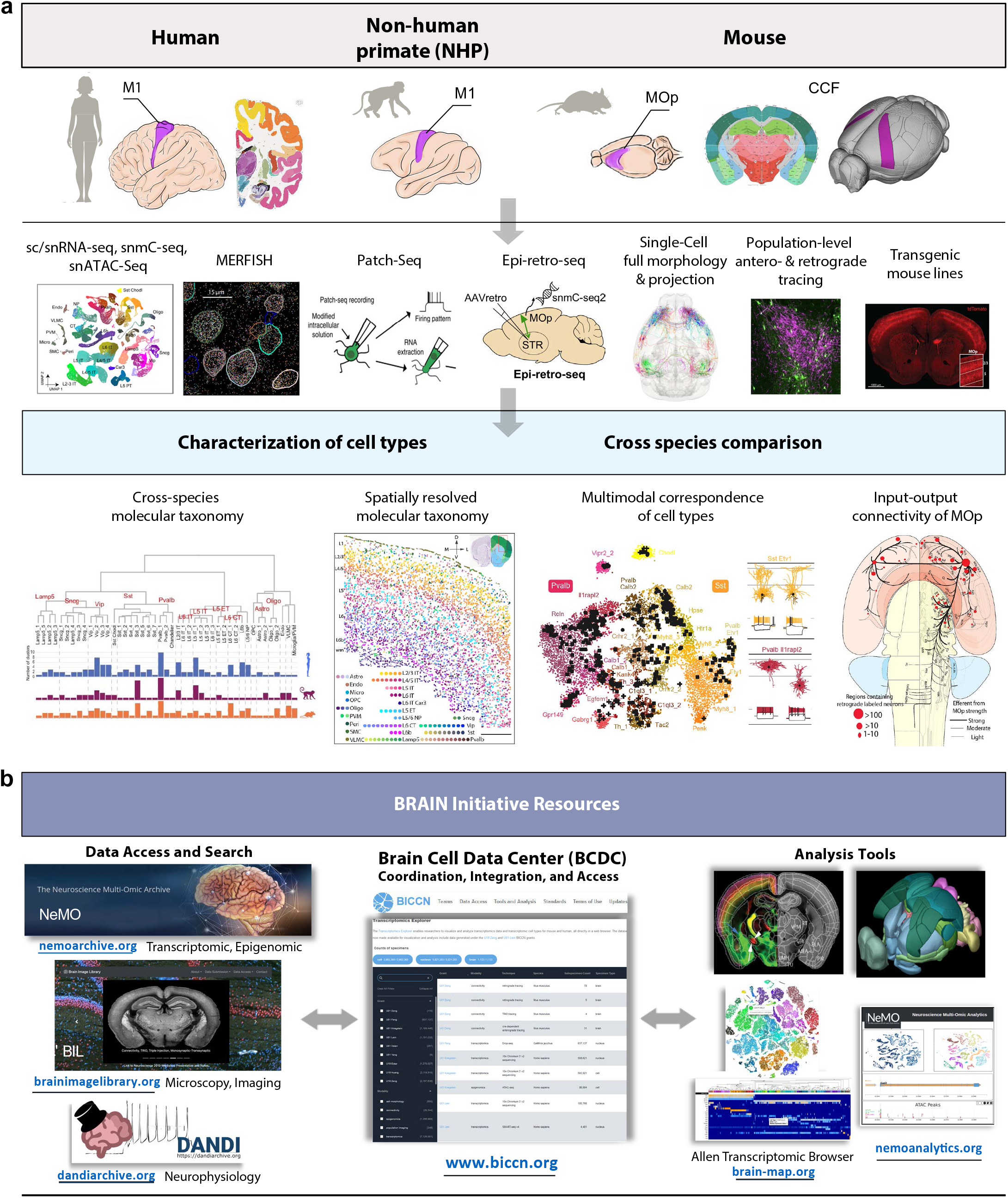
Summary of experimental and computational approaches taken as well as community resources generated by the BICCN. **a**, Comprehensive characterization of cell types in the primary motor cortex (MOp) of three mammalian species using multiple approaches spanning molecular, genetic, physiological and anatomical domains. Integration of these datasets leads to a cohesive multimodal description of cell types in the mouse MOp and a cross-species molecular taxonomy of MOp cell types. **b**, The multimodal datasets are organized by the Brain Cell Data Center (BCDC), archived in the Neuroscience Multi-omic (NeMO) Archive (for molecular datasets), Brain Image Library (BIL, for imaging datasets) and Distributed Archive for Neurophysiology Data Integration (DANDI, for electrophysiology data), and made publicly available through the BICCN web portal www.biccn.org.

**Table 1.**
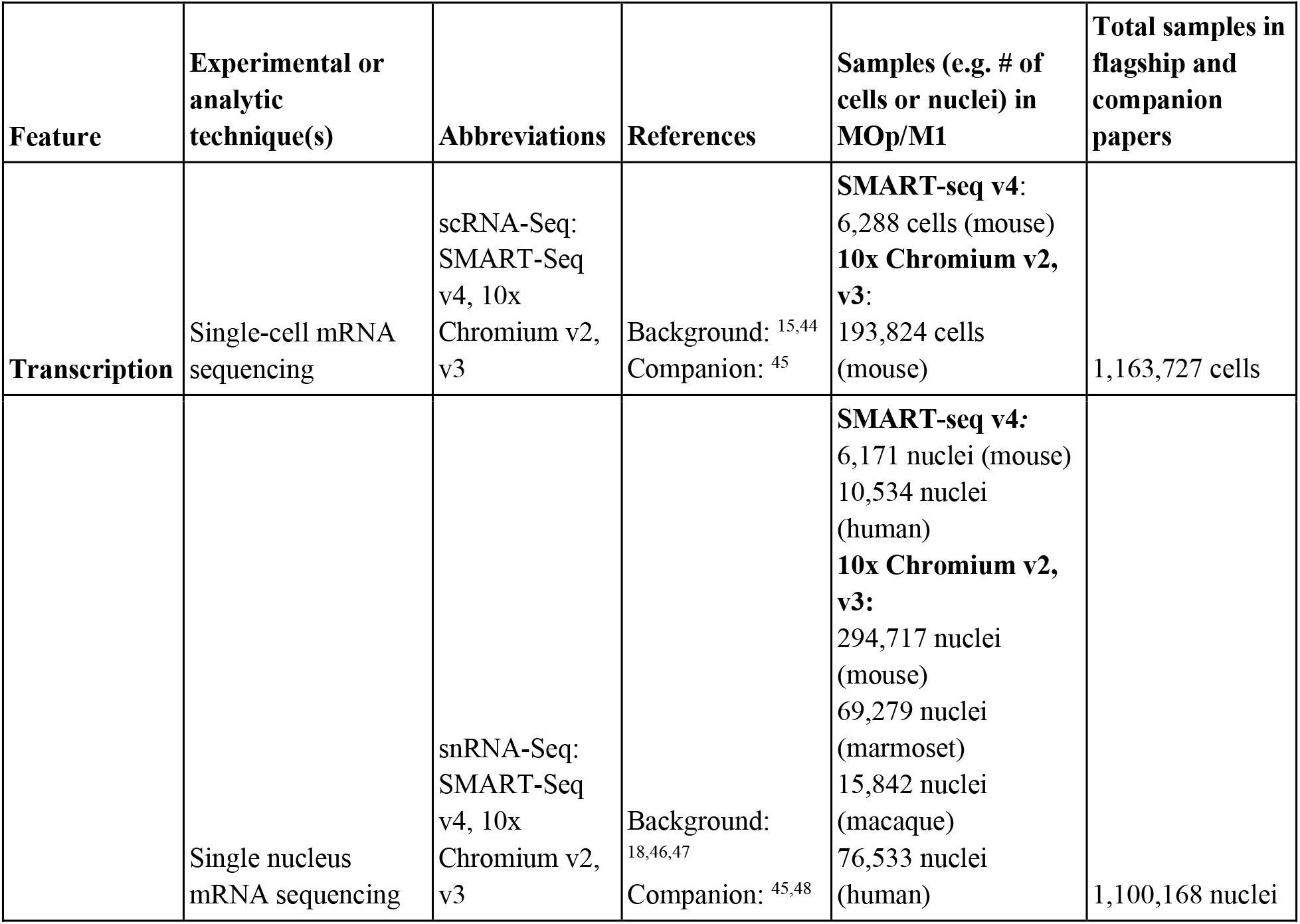
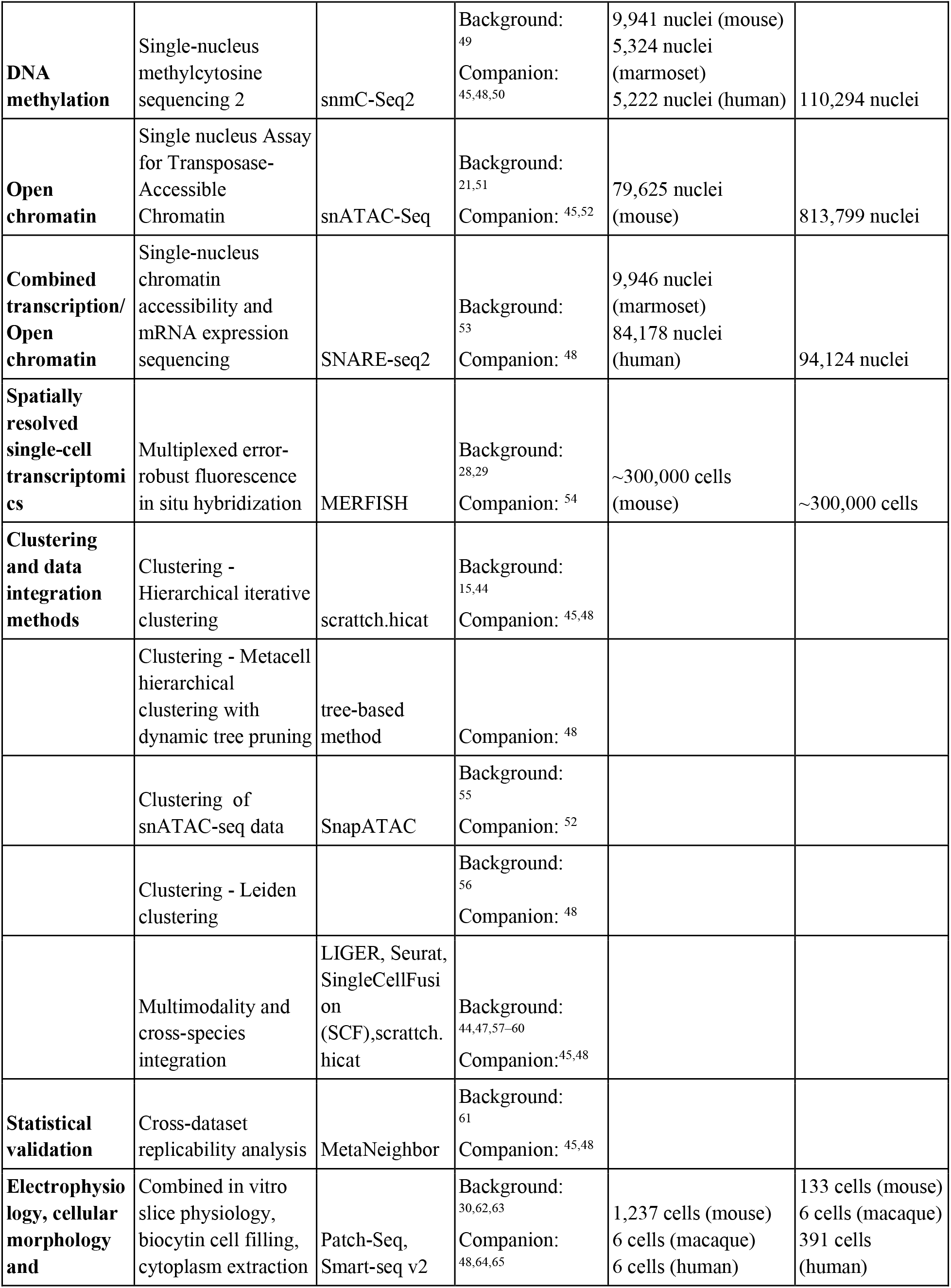
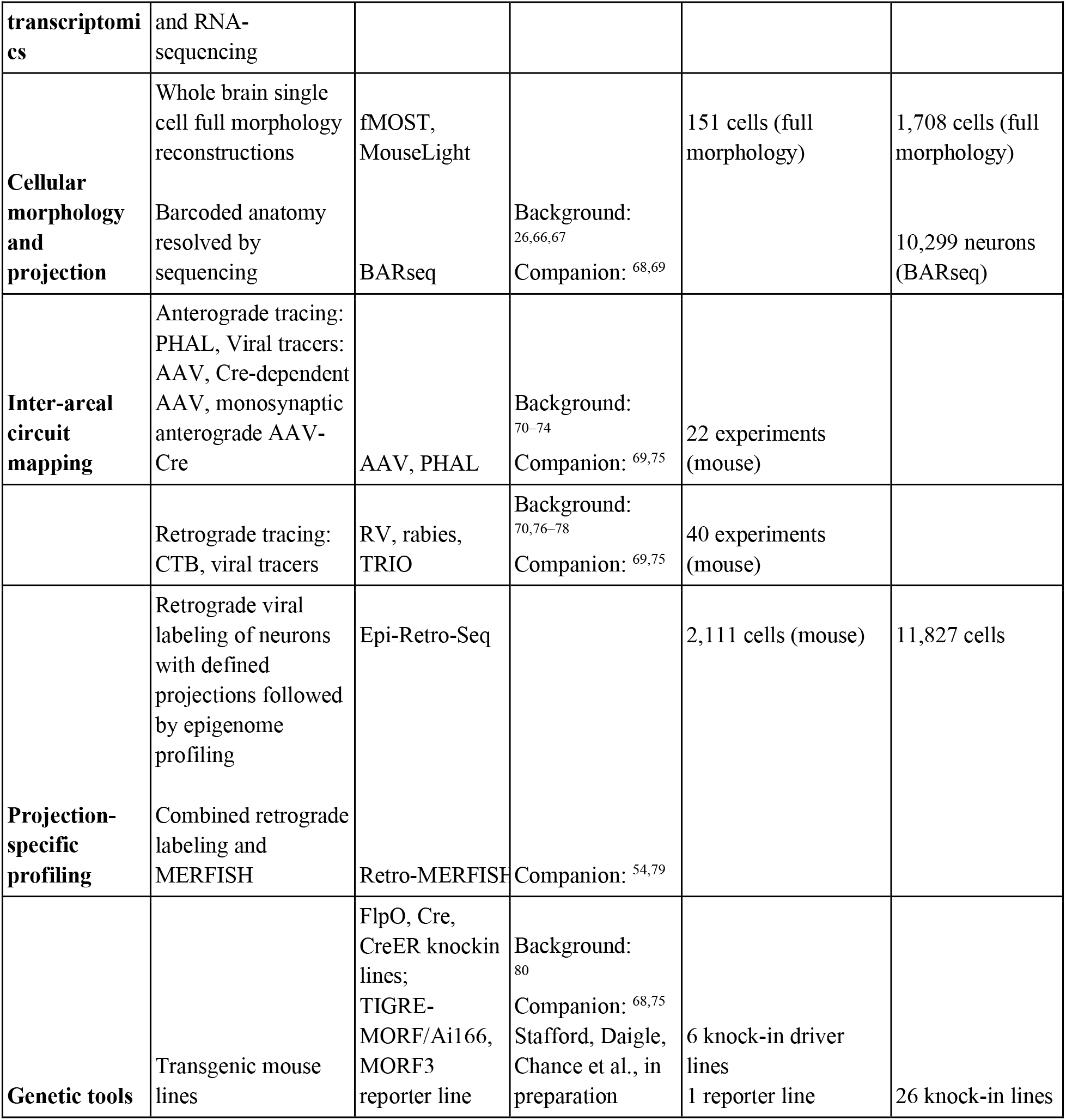
Experimental and computational techniques used in this study and associated datasets

| Feature | Experimental or analytic technique(s) | Abbreviations | References | Samples (e.g. # of cells or nuclei) in MOp/M1 | Total samples in flagship and companion papers |
| --- | --- | --- | --- | --- | --- |
| Transcription | Single-cell mRNA sequencing | scRNA-Seq:<br>SMART-Seq v4, 10x Chromium v2, v3 | Background: <sup>15,44</sup><br>Companion: <sup>45</sup> | **SMART-seq v4:**<br>6,288 cells (mouse)<br>**10x Chromium v2, v3:**<br>193,824 cells (mouse) | 1,163,727 cells |
|  | Single nucleus mRNA sequencing | snRNA-Seq:<br>SMART-Seq v4, 10x Chromium v2, v3 | Background: <sup>18,46,47</sup><br>Companion: <sup>45,48</sup> | **SMART-seq v4:**<br>6,171 nuclei (mouse)<br>10,534 nuclei (human)<br>**10x Chromium v2, v3:**<br>294,717 nuclei (mouse)<br>69,279 nuclei (marmoset)<br>15,842 nuclei (macaque)<br>76,533 nuclei (human) | 1,100,168 nuclei |
| **DNA methylation** | Single-nucleus methylcytosine sequencing 2 | snmC-Seq2 | Background: <sup>49</sup><br>Companion: <sup>45,48,50</sup> | 9,941 nuclei (mouse)<br>5,324 nuclei (marmoset)<br>5,222 nuclei (human) | 110,294 nuclei |
| **Open chromatin** | Single nucleus Assay for Transposase-Accessible Chromatin | snATAC-Seq | Background: <sup>21,51</sup><br>Companion: <sup>45,52</sup> | 79,625 nuclei (mouse) | 813,799 nuclei |
| **Combined transcription/ Open chromatin** | Single-nucleus chromatin accessibility and mRNA expression sequencing | SNARE-seq2 | Background: <sup>53</sup><br>Companion: <sup>48</sup> | 9,946 nuclei (marmoset)<br>84,178 nuclei (human) | 94,124 nuclei |
| **Spatially resolved single-cell transcriptomics** | Multiplexed error-robust fluorescence in situ hybridization | MERFISH | Background: <sup>28,29</sup><br>Companion: <sup>54</sup> | ~300,000 cells (mouse) | ~300,000 cells |
| **Clustering and data integration methods** | Clustering - Hierarchical iterative clustering | scrattch.hicat | Background: <sup>15,44</sup><br>Companion: <sup>45,48</sup> |  |  |
|  | Clustering - Metacell hierarchical clustering with dynamic tree pruning | tree-based method | Companion: <sup>48</sup> |  |  |
|  | Clustering of snATAC-seq data | SnapATAC | Background: <sup>55</sup><br>Companion: <sup>52</sup> |  |  |
|  | Clustering - Leiden clustering |  | Background: <sup>56</sup><br>Companion: <sup>48</sup> |  |  |
|  | Multimodality and cross-species integration | LIGER, Seurat, SingleCellFusion (SCF),scrattch.hicat | Background: <sup>44,47,57-60</sup><br>Companion: <sup>45,48</sup> |  |  |
| **Statistical validation** | Cross-dataset replicability analysis | MetaNeighbor | Background: <sup>61</sup><br>Companion: <sup>45,48</sup> |  |  |
| **Electrophysiology, cellular morphology and** | Combined in vitro slice physiology, biocytin cell filling, cytoplasm extraction | Patch-Seq, Smart-seq v2 | Background: <sup>30,62,63</sup><br>Companion: <sup>48,64,65</sup> | 1,237 cells (mouse)<br>6 cells (macaque)<br>6 cells (human) | 133 cells (mouse)<br>6 cells (macaque)<br>391 cells (human) |
| transcriptomics | and RNA-sequencing |  |  |  |  |
| Cellular morphology and projection | Whole brain single cell full morphology reconstructions | fMOST, MouseLight |  | 151 cells (full morphology) | 1,708 cells (full morphology) |
|  | Barcoded anatomy resolved by sequencing | BARseq | Background: 26,66,67<br>Companion: 68,69 |  | 10,299 neurons (BARseq) |
| Inter-areal circuit mapping | Anterograde tracing: PHAL, Viral tracers: AAV, Cre-dependent AAV, monosynaptic anterograde AAV-Cre | AAV, PHAL | Background: 70–74<br>Companion: 69,75 | 22 experiments (mouse) |  |
|  | Retrograde tracing: CTB, viral tracers | RV, rabies, TRIO | Background: 70,76–78<br>Companion: 69,75 | 40 experiments (mouse) |  |
| Projection-specific profiling | Retrograde viral labeling of neurons with defined projections followed by epigenome profiling | Epi-Retro-Seq |  | 2,111 cells (mouse) | 11,827 cells |
|  | Combined retrograde labeling and MERFISH | Retro-MERFISH | Companion: 54,79 |  |  |
| Genetic tools | Transgenic mouse lines | FlpO, Cre, CreER knockin lines; TIGRE-MORF/Ai166, MORF3 reporter line | Background: 80<br>Companion: 68,75<br>Stafford, Daigle, Chance et al., in preparation | 6 knock-in driver lines<br>1 reporter line | 26 knock-in lines |

**Table 2.**
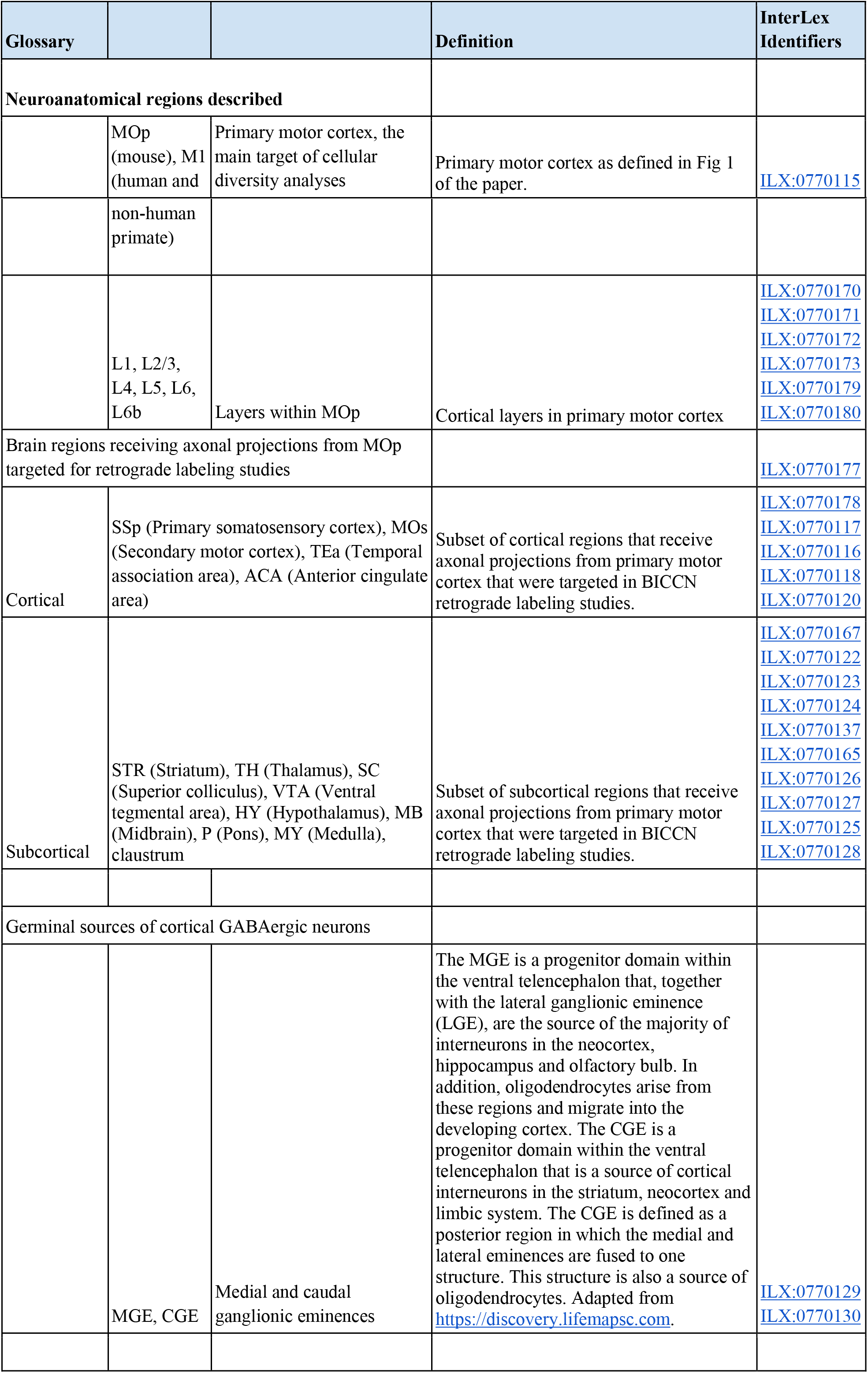
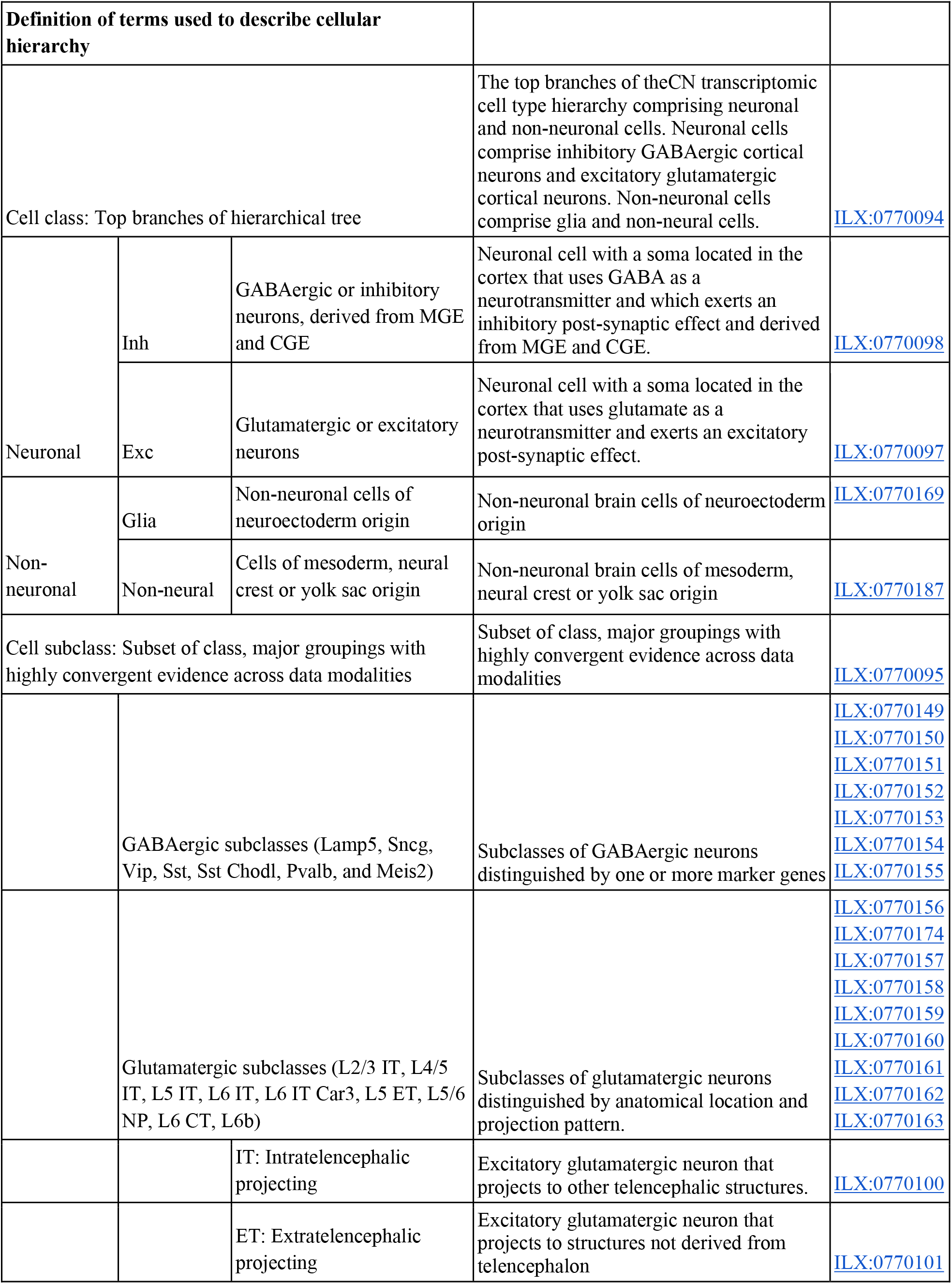
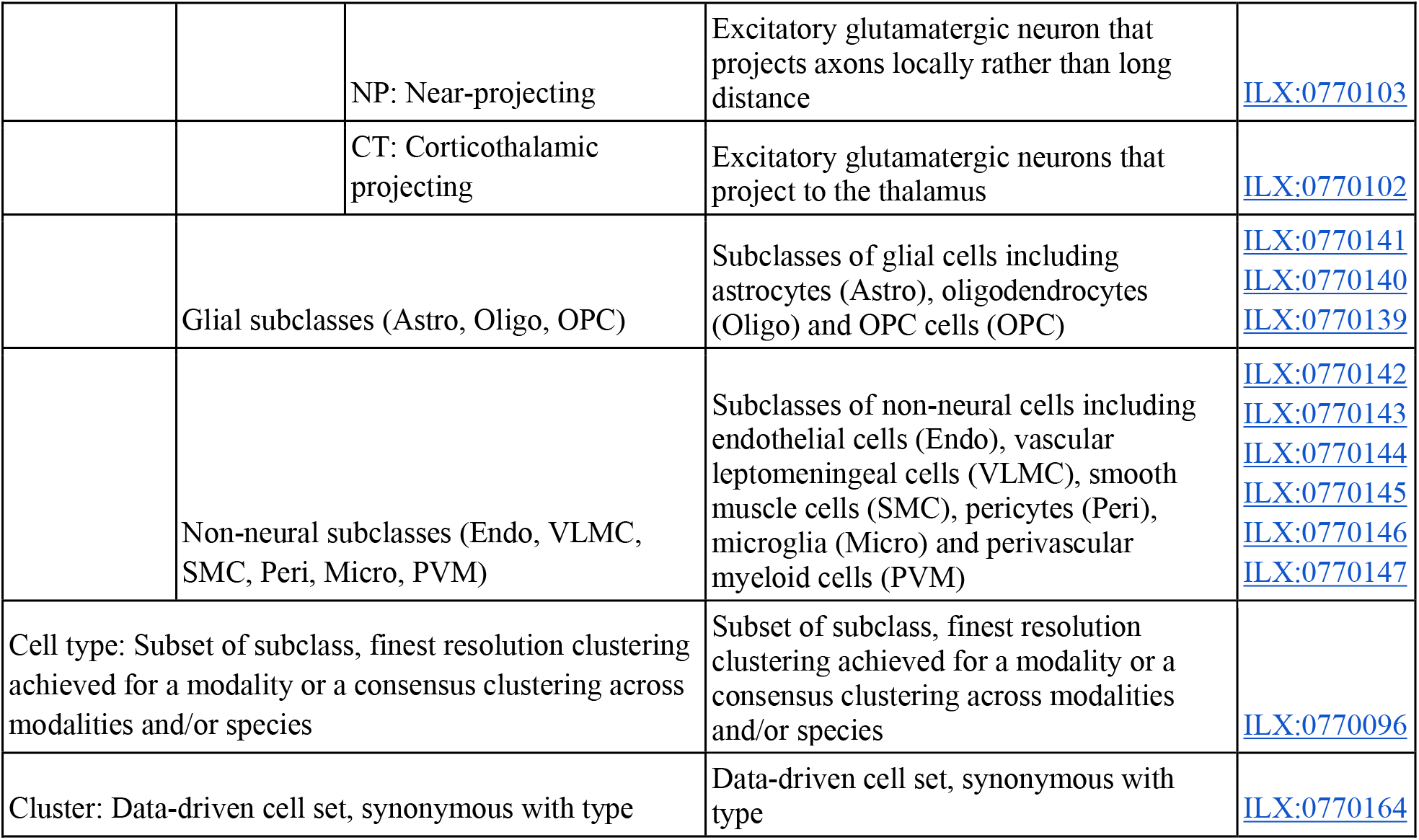
Glossary

Major findings from this coordinated consortium project include:

- Combined single-cell transcriptomic and epigenomic analysis reveals a unified molecular genetic landscape of adult cortical cell types that integrates gene expression, chromatin state and DNA methylation maps.
- Combination of single-cell-omics, MERFISH-based spatially resolved single-cell transcriptomics and Patch-seq generates a census and atlas of cell types, including their population demographics of type, proportion, and spatial distribution across cortical layers and sublayers.
- Comparative analysis of mouse, marmoset and human transcriptomic types achieves a unified cross-species taxonomy of cortical cell types with their hierarchical organization that reflects developmental origins; transcriptional similarity of cell type granularity across species varies as a function of evolutionary distance.
- We observed both highly conserved gene expression and epigenomic signatures of cell identity across species, as well as a large set of species-specific cell type gene expression profiles suggesting a high degree of evolutionary specialization.
- The overall correspondence among transcriptomic, epigenetic, spatial transcriptomic, morphological, and intrinsic physiological datasets reinforces the transcriptomic classification of neuronal subclasses and distinctive types, demonstrating their biological validity and genomic underpinnings, and also reveals continuously varying properties along these axes among some neuronal subclasses and types.
- Multi-faceted anatomic studies yield a cellular resolution wiring diagram of mouse MOp anchored on major transcriptome-defined projection types, including input-output connectivity at subpopulation level and output pathways at genetically-defined single-cell level.
- The long-range axon projection patterns of individual glutamatergic excitatory neurons exhibit a complex and diverse range of relationships (between one-to-one and many-to-many) with transcriptomic and epigenetic types, suggesting another level of regulation in defining single-cell connectional specificity.
- Cell type transcriptional and epigenetic signatures can guide the generation of an extensive genetic toolkit for targeting glutamatergic pyramidal neuron types and fate mapping their progenitor types.
- Multi-site coordination within BICCN and data archives allows a high degree of standardization, computational integration, and creation of open data resources for community dissemination of data, tools and knowledge.

## RESULTS

### Molecular definition of cell types in MOp

The mouse MOp molecular taxonomy is derived from 9 datasets, including seven sc/snRNA-seq datasets and one each of snmC-Seq2 and snATAC-Seq datasets (companion paper ^45^). The combined seven sc/snRNA-seq datasets (>700,000 cells total) had the advantages of large number of cells profiled using the droplet-based 10x Chromium v2 or v3 method and deep full-length sequencing using the plate-based SMART-Seq v4 method, resulting in a consensus transcriptomic taxonomy for the mouse MOp with the greatest resolution compared to other data types, containing 116 clusters or transcriptomic types (t-types), 90 of which were neuronal types^45^. We used this mouse MOp transcriptomic taxonomy as the anchor for comparison and cross-correlation of cell-type classification results across all data types. We further utilized two computational approaches, SingleCellFusion (SCF) and LIGER, to combine the seven transcriptomic with two epigenomic datasets and derive an integrated molecular taxonomy consisting of 56 neuronal cell types (corresponding to the 90 transcriptomic neuronal types) for the mouse MOp, with highly consistent molecular profiles based on transcriptomics, DNA-methylation, and open chromatin ^45^ (**Fig. 2a**). Critically, this integrated taxonomy enabled us to link RNA transcripts with epigenomic marks identifying potential cell-type-specific cis-regulatory elements (CREs) and transcriptional regulatory networks. Similarly, we established M1 cell type taxonomies for human (127 t-types) and marmoset (94 t-types) by unsupervised clustering of snRNA-seq data, followed by integration with epigenomic datasets (companion paper ^48^).

**Figure 2.**
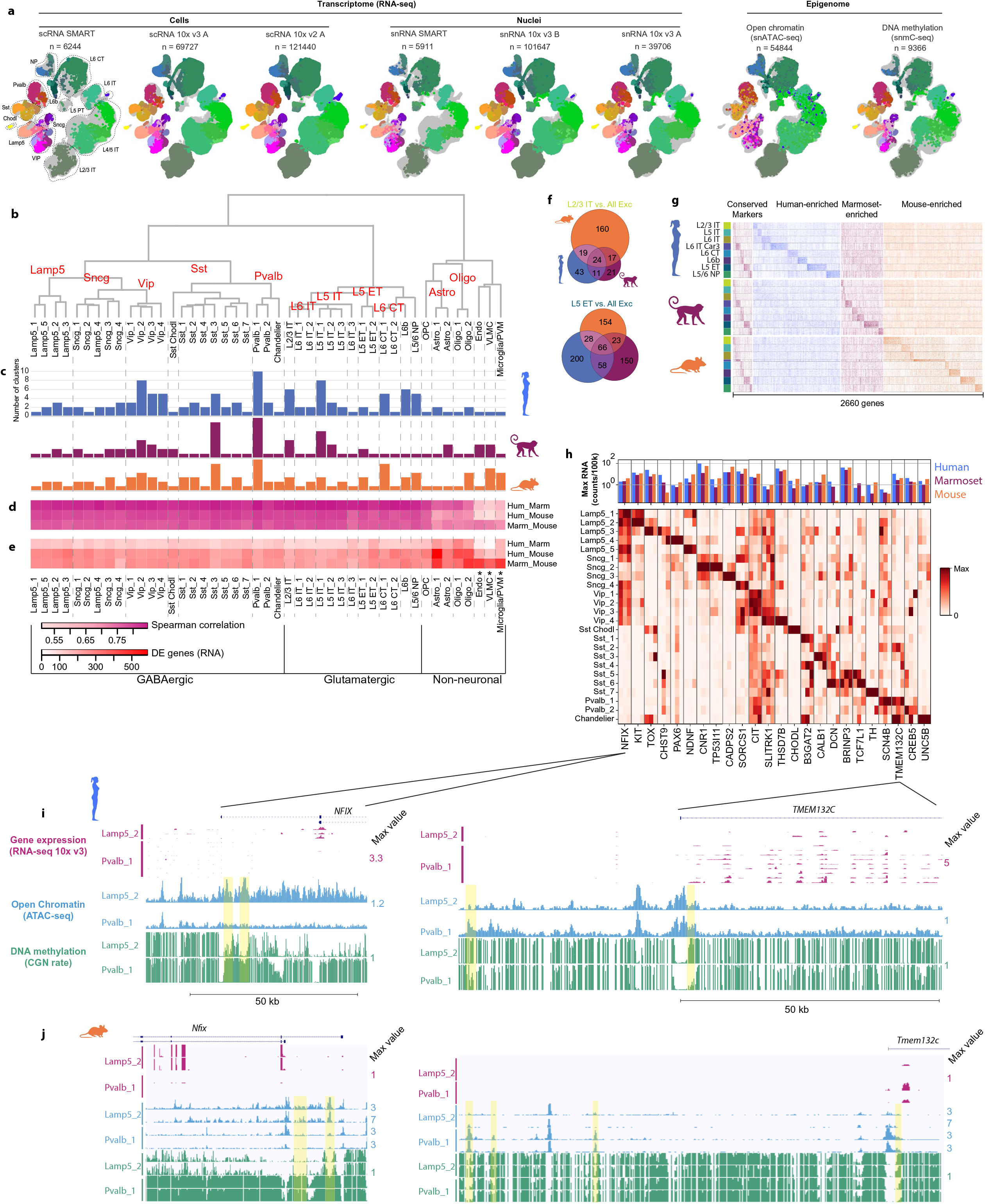
MOp consensus cell type taxonomy. **a**, Integrated transcriptomic and epigenomic datasets using SCF show consistent molecular cell-type signatures as revealed by a low-dimensional embedding in mouse MOp. Each Uniform Manifold Approximation and Projection (UMAP) plot represents one dataset. Colors indicate different subclasses. **b,** Dendrogram of integrated human, marmoset, and mouse cell types based on single nucleus RNA-seq datasets (10x Chromium v3). **c,** Number of within-species clusters that are included in each cross-species cluster. **d-e,** For each consensus cluster, correlations (d) and differentially expressed (DE; Wilcoxon test, FDR < 0.01 and log*_e_* fold-change > 2) genes (e) between pairs of species. Asterisks denote non-neuronal populations that were under-sampled in human. **f,** Venn diagrams of shared DE genes between species for L2/3 IT and L5 ET glutamatergic neuron subclasses. **g**, Conserved and species-specific DE genes for all glutamatergic subclasses. Heatmap shows gene expression normalized by the maximum for each gene for up to 50 randomly sampled nuclei from each subclass and species. **h**, Conserved markers of GABAergic neuron types across three species. **i-j**, Genome browser showing transcriptomic and epigenetic signatures for gene markers of Lamp5_2 (*NFIX*) and Pvalb_1 (*TMEM132C*) GABAergic neurons in human (i) and mouse (j). Yellow bars highlight sites of open chromatin and DNA hypomethylation in the cell type with corresponding marker expression.

To establish a consensus classification of MOp/M1 cell types among mouse, human and marmoset, we integrated snRNA-seq datasets across species and identified 45 conserved transcriptomic types that spanned three major cell classes, including 24 GABAergic, 13 glutamatergic, and 8 non-neuronal types (**Fig. 2b, Extended Data Fig. 1**). These types were grouped into broader subclasses based on shared developmental origin for GABAergic inhibitory neurons [i.e., three caudal ganglionic eminence (CGE)-derived subclasses (Lamp5, Sncg and Vip) and two medial ganglionic eminence (MGE)-derived subclasses (Sst and Pvalb)], layer and projection pattern in mouse for glutamatergic excitatory neurons [i.e., intratelencephalic (IT), extratelencephalic (ET), corticothalamic (CT), near-projecting (NP) and layer 6b (L6b)], and non-neuronal functional subclass (e.g., oligodendrocytes and astrocytes) (**Table 2**). Note that the layer 5 extratelencephalic (L5 ET) neurons had been named as pyramidal tract (PT) neurons or subcerebral projection neurons (SCPN) in the literature ^81,82^; in this study we chose to use the name L5 ET for this subclass of neurons to be more representative across cortical areas and species (**Supplementary Notes**). The resolution of this cross-species conserved taxonomy was lower than that derived from each species alone, due to gene expression variations among species. The degree of species alignments varied across consensus types (**Fig. 2c**); some types could be aligned one-to-one (e.g., Lamp5_1, L6 IT_3), while others aligned several-to-several (e.g., Pvalb_1, L2/3 IT, L5 IT_1). This may reflect over- or under-clustering, limitations in aligning highly similar cell types or species-specific expansion of cell-type diversity (companion paper ^65^).

We hypothesized that cell types would share more similar gene expression profiles between human and marmoset than between either primate and mouse because primates share a more recent common ancestor. Indeed, we found that between primates, transcriptomic profiles of consensus cell types were more correlated and had 25-50% fewer differentially expressed (DE) genes than between primates and mouse (**Fig. 2d,e**). Three non-neuronal types had greater spearman correlations of overall gene expression (**Fig. 2d**, right columns) between marmoset and mouse likely because non-neuronal cells were undersampled in human M1 resulting in fewer rare types ^48^. Robust conservation of cell types across mammals, including types with known specificity in electrical properties and connectivity such as chandelier cells and long-range projecting *Sst*-expressing cells (*Sst Chodl*), is strong evidence for the functional significance of these types.

Glutamatergic subclasses expressed many marker genes (using Seurat’s FindAllMarkers function with test.use set to ‘roc’, >0.7 classification power) compared to other subclasses, and the majority of markers were species-specific (**Fig. 2f,g**). The evolutionary divergence of marker gene expression may reflect species adaptations or relaxed constraints on genes that can be substituted with others for related cellular functions. Subclasses also had a core set of marker genes that were conserved across all three species (**Fig. 2g**); these genes are candidates for consistent labeling of consensus cell types and for determining the conserved features of those cells that are central to their function. GABAergic consensus types also had conserved markers with similar absolute expression levels across species (**Fig. 2h**, bar plots) and relatively specific expressions compared to other cell types (**Fig. 2h**, heatmap). Marker genes of Lamp5_2 (*NFIX*) and Pvalb_1 (*TMEM132C*) GABAergic neurons showed evidence for cell-type-specific enhancers located in regions of open chromatin and DNA hypomethylation in both human (**Fig. 2i**) and mouse (**Fig. 2j**).

In summary, the multi-omic approach reveals a unified molecular genetic landscape of cortical cell types that integrates gene expression, chromatin state and DNA methylation maps and yields a robust molecular classification of cell types that is consistent between transcriptomic and epigenomic analyses. These studies further allow the identification of putative regulatory elements associated with cell type identity. Cell types are generally conserved between primates and rodents, and have a small number of conserved marker genes that are candidates for consistent labeling of conserved cell types.

### Spatially resolved cell atlas of the mouse MOp by MERFISH

Sequencing-based single-cell methods require dissociation of cells from tissues, and hence the spatial organization of neuronal and non-neuronal cells, which is critical for brain function, is lost. To obtain a spatially resolved cell atlas of the mouse MOp region, we used MERFISH, a single-cell transcriptome imaging method ^28,29^, to identify cell types *in situ* and map their spatial organization. We selected a panel of 258 genes (254 of which passed quality control) to image by MERFISH, on the basis of both prior knowledge of marker genes for major subclasses of cells in the cortex and marker genes differentially expressed in the neuronal clusters identified by the sn/scRNA-seq experiments, and we imaged ∼300,000 individual cells across the MOp and its vicinity (companion paper ^54^).

Clustering analysis of the MERFISH-derived single-cell expression profiles resulted in a total of 95 cell clusters in MOp, including 42 GABAergic, 39 glutamatergic, and 14 non-neuronal clusters (**Fig. 3a,b**), as well as four distinct cell clusters observed exclusively outside the MOp (in striatum or lateral ventricle). These 95 clusters showed excellent correspondence with the 116 cell clusters identified by the sn/scRNA-seq datasets ^54^. MERFISH analysis also revealed clusters not identified by scRNA-seq and vice versa, mostly in the form of refined splitting of clusters ^54^.

**Figure 3.**
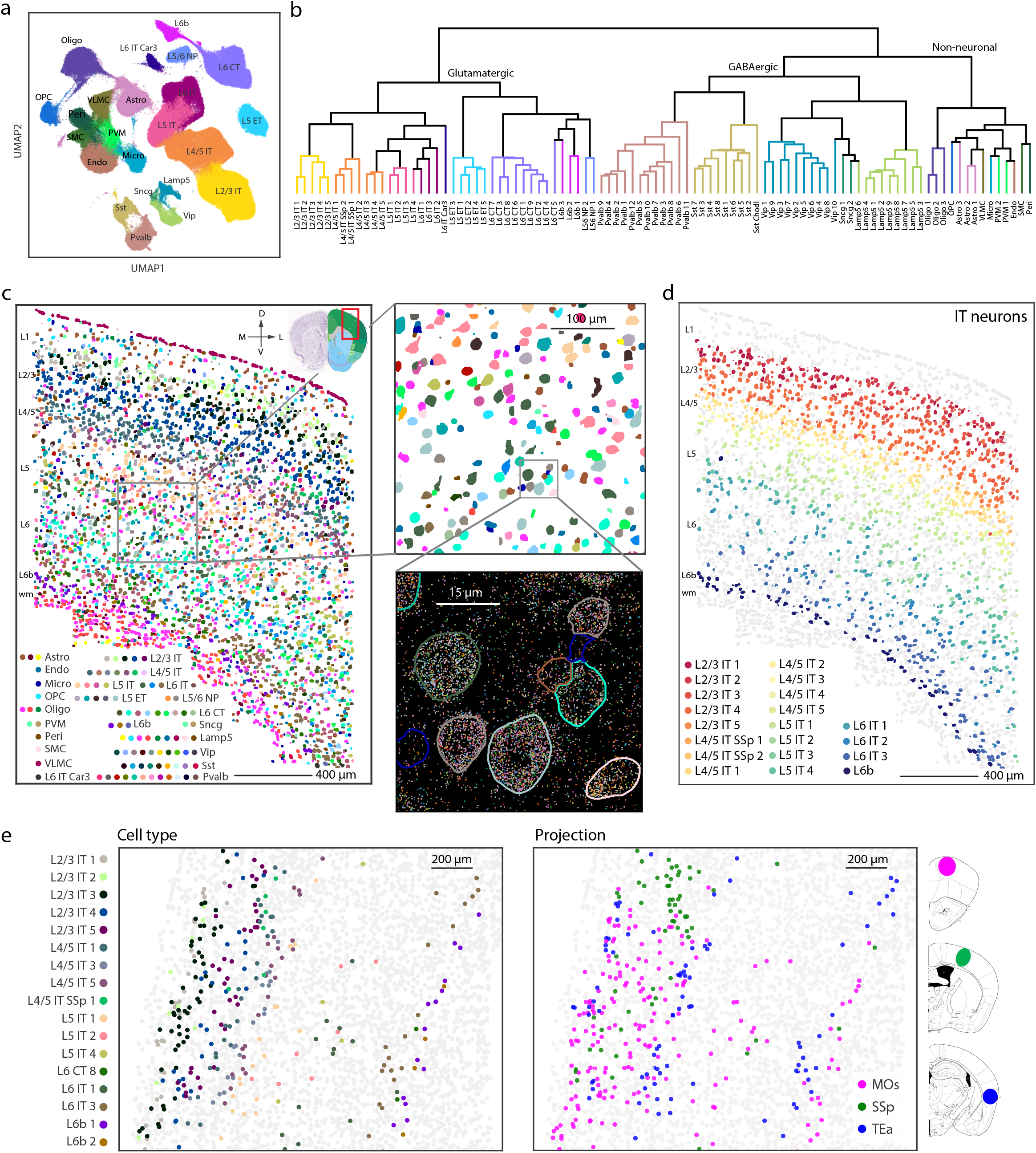
In situ cell-type identification, spatial mapping and projection mapping of individual cells in the MOp by MERFISH. a, UMAP of the ∼300,000 cells in the mouse MOp imaged by MERFISH. Cell clusters are grouped into 23 subclasses, and all cells in the same subclass are plotted in the same color. b, Dendrogram showing the hierarchical relationship among the 39 glutamatergic, 42 GABAergic, and 14 non-neuronal clusters in the mouse MOp identified by MERFISH, colored by the subclass that each cluster belongs to. c, Left: Spatial map of the cell clusters identified in a coronal slice (Bregma +0.90), with cells colored by their cluster identity as shown in the color index. Top right: Zoom-in map of the boxed region of the left panel. Bottom right: Spatial localization of individual RNA molecules in the boxed region of the top right panel, colored by their gene identity. The segmented cell boundaries are colored according to the cell clusters they belong to. d, The IT neurons in the same coronal slice as shown in c. The IT neurons are colored by their cluster identity, as shown in the color index, together with L6b cells in dark blue to mark the bottom border of the cortex. All other cells are shown in grey. e, Neuronal cluster identities of the cells projecting into three other regions of the brain, secondary motor cortex (MOs), primary somatosensory cortex (SSp), and temporal association area (TEa). Dye-labeled cholera toxin b (CTb) are used as retrograde tracers, and the CTb signals and the MERFISH gene panel are imaged in the MOp to determine both the cell cluster identities (left panel) and projection targets (right panel) of individual cells. Only clusters with 3 or more cells labeled by CTb are shown in color and the remaining cells are shown in grey.

The spatial distribution of the clusters derived from MERFISH showed a complex, laminar organization of cells in the MOp (**Fig. 3c**). MERFISH data divided glutamatergic neurons into IT, ET, NP, CT, and L6b subclasses, each of which were further divided into finer clusters. Many of these clusters adopted narrow distributions along the cortical depth direction that subdivided individual cortical layers, though often without discrete boundaries ^54^. Notably, IT cells, the largest branch of neurons in the MOp, formed a largely continuous spectrum of cells with gradual changes both in their expression profiles and in their cortical depth positions, in a highly correlated manner ^54^ (**Fig. 3d**). The five major subclasses of GABAergic neurons (Lamp5, Sncg, Vip, Sst and Pvalb) were also divided into finer clusters. Interestingly, many individual GABAergic clusters showed layered distribution as well, preferentially residing within one or two cortical layers ^54^. Among the non-neuronal cell clusters, VLMCs formed the out-most layer of cells of the cortex, mature oligodendrocytes and some astrocytes were enriched in white matter, whereas other major subclasses of non-neuronal cells were largely dispersed across all layers. In addition to the laminar organization, MERFISH analysis also revealed interesting spatial distributions of cell types along the medial-lateral and anterior-posterior axes ^54^. Overall, the 95 neuronal and non-neuronal cell clusters in the MOp form a complex spatial organization refining traditionally defined cortical layers.

Integration of retrograde tracing with MERFISH (Retro-MERFISH) further allowed us to map the projection targets of different neuronal cell types in the MOp. By injecting retrograde tracers into several different cortical areas (secondary motor cortex, primary somatosensory cortex, and temporal association area) and imaging retrograde labels together with the MERFISH gene panel in the MOp (**Fig. 3e**), we observed that all three examined target regions received inputs from multiple cell clusters in the MOp, primarily from IT cells. In addition, each IT cluster projected to multiple regions, with each region receiving input from a different composition of IT clusters^54^. Overall, the projection of MOp neurons does not follow a simple “one cell type to one target region” pattern, but rather forms a complex many-to-many network.

In summary, these MERFISH measurements revealed the spatial organization of neuronal and non-neuronal cell types in the MOp with an unprecedented resolution and granularity. Integration of MERFISH with retrograde tracing further allowed determination of both gene expression profiles and projection targets with single-cell resolution, revealing the compositions and spatial distributions of MOp neurons that project to several cortical regions.

### Multimodal analysis of cell types with Patch-seq

To characterize the electrophysiological and morphological phenotypes and laminar location of the transcriptomically identified cell types, i.e., the t-types, we used the recently developed Patch-seq technique ^30,62^. We patched >1,300 neurons in MOp of adult mice, recorded their electrophysiological responses to a set of current steps, filled them with biocytin to recover their morphology (∼50% of the cells) and obtained their transcriptomes using Smart-seq2 sequencing (companion paper ^64^). We mapped these cells to the mouse MOp transcriptomic taxonomy ^45^. Our dataset covered all major subclasses of glutamatergic and GABAergic neurons, with cells assigned to 77 t-types (**Fig. 4a**). This allowed us to describe the electrophysiological and morphological phenotypes of most t-types (see examples in **Fig. 4b,c**).

**Figure 4.**
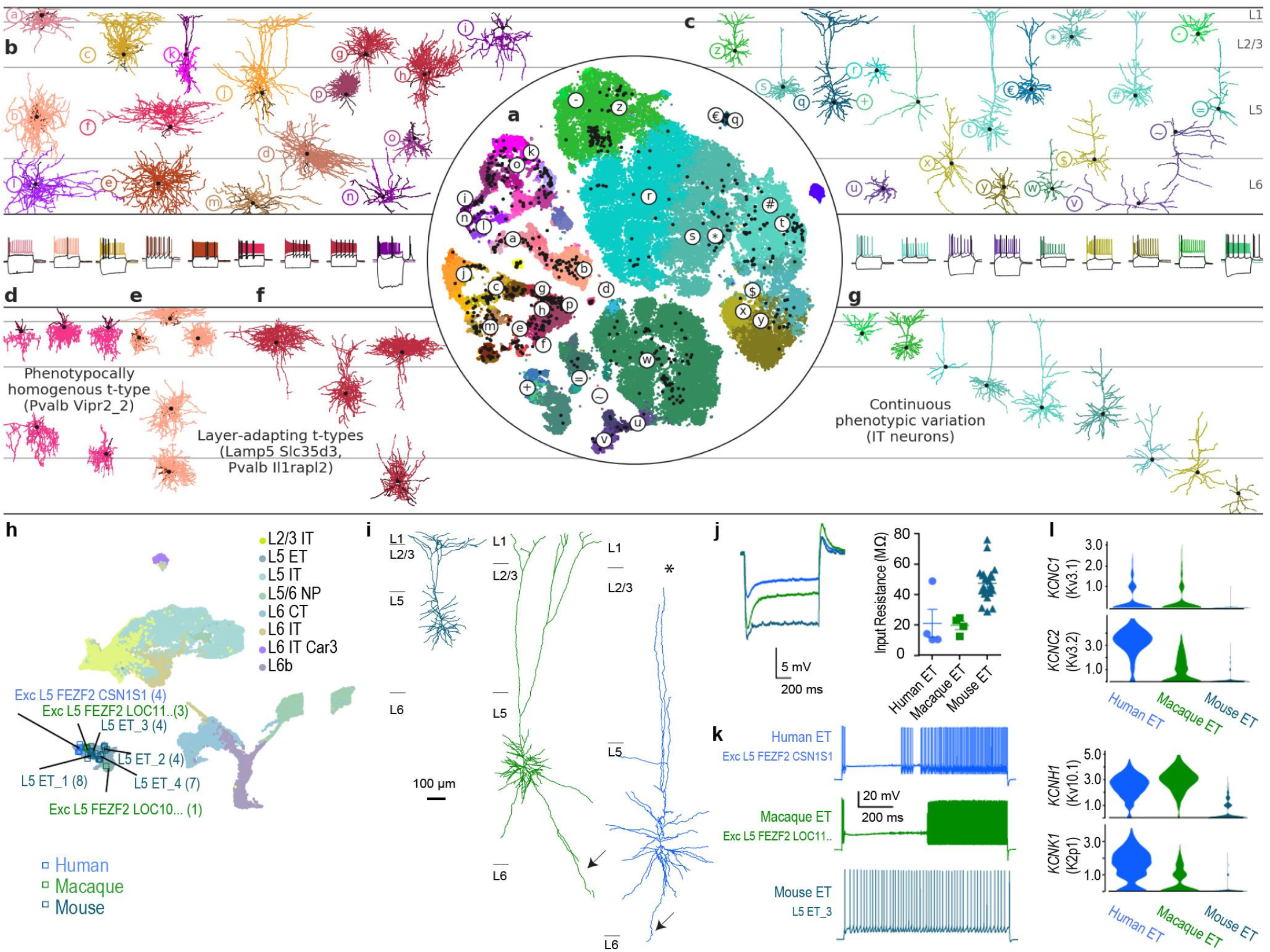
Correspondence between transcriptomic and morpho-electrical properties of mouse MOp neurons by Patch-seq, and cross-species comparison of L5 ET neurons. **a**, t-SNE of the scRNA-seq 10x v2 dataset ^45^ with the Patch-seq neurons (black dots) positioned on top of it ^84^. b, Examples of GABAergic interneuron morphologies and electrophysiological recordings (below). Letters refer to cells marked in a. c, Examples of glutamatergic excitatory neuron morphologies and electrophysiological recordings. d, Example of a phenotypically homogenous t-type (Pvalb Vipr2_2, chandelier neurons). e-f, Two examples of t-types showing layer-adapting morphologies (e, Lamp5 Slc35d3, neurogliaform cells; f, Pvalb Il1rapl2, fast-spiking basket cells). g, Example of a transcriptomic subclass (excitatory IT neurons) that shows continuous within-subclass co-variation between distances in transcriptomic space and morphological space (compare the color ordering in a (right) with the color ordering in g. h, UMAP visualization of cross-species integration of snRNA-seq data for glutamatergic neurons isolated from mouse, macaque and human, with colors corresponding to cell subclass. Patch-seq samples mapping to various ET neuron types are denoted by squares, color-coded by species. i, Dendritic reconstructions of L5 ET neurons. The human (Exc L5 FEZF2 CSN1S1) and macaque (Exc L5 FEZF2 LOC114676463) neurons display classical Betz cell features, including taproot dendrites (arrows). Note, the human neuron is truncated (asterisk) before reaching the pial surface. **j**, Voltage response of mouse, macaque and human ET neurons to a 1 s, -300 pA current injection (left). Input resistance is low in all species, but exceptionally low in human and macaque Betz cells. Error bars represent SEM (right; macaque n=4, human n=4, mouse n=22; FDR corrected two-sided Wilcoxon ranked sum test (human vs mouse W=12, p = 0.31, d=2.09; human vs monkey W = 5, p = .49, d=.08; monkey v mouse W = 0 p = .0004., d =2.5). **k**, Example spike trains in response to a 10s suprathreshold current injection. Macaque and human L5 ET neurons tended to respond with a distinctive, biphasic firing pattern. **l**, Violin plots of enriched potassium channel gene expression in human and macaque compared to mouse L5 ET neurons.

We found that the measured morpho-electrical (me) phenotype of a neuron was largely determined by its transcriptomic subclass, with different subclasses having distinct phenotypes. For example, Sst interneurons were often characterized by large membrane time constants, pronounced hyperpolarization sag, and rebound firing after stimulation offset. However, within each subclass, there was substantial variation in electrophysiological and morphological properties between t-types. This variation was not random but organized such that transcriptomically similar t-types had more similar morpho-electric properties than distant t-types. For example, excitatory t-types from the IT subclasses with more similar transcriptomes were located also at adjacent cortical depths, suggesting that distances in t-space co-varied with distances in the me-space, even within a layer (**Fig. 4g**). Likewise, the electrophysiological properties of Sst interneurons varied continuously across the transcriptomic landscape ^64^.

At the level of single t-types, we found that some t-types showed layer-adapting morphologies across layers (**Fig. 4e,f**) or even considerable within-type morpho-electric variability within a layer. For example, Vip Mybpc1_2 neurons had variable rebound firing strength after stimulation offset. Surprisingly few t-types were entirely homogeneous with regard to the measured morpho-electric properties (**Fig. 4d**).

In summary, we found that the morpho-electric phenotype of a neuron in MOp was primarily determined by the major subclass of neurons it belonged to, with different subclasses being transcriptomically as well as morpho-electrically distinct. Within each subclass, variation in electrophysiological and morphological properties often appeared to be continuous across the transcriptomic landscape, without clear-cut boundaries between neighbouring t-types.

Patch-seq also permits direct comparison of the morpho-electric properties of homologous cell types across species ^48^. Here we focused our analysis on one of the most recognizable mammalian neuron types, the gigantocellular Betz cells found in M1 of primates and large carnivores. These neurons are predicted to be in the layer 5 ET (L5 ET) subclass ^48^, which also contains the homologous corticospinal projecting neurons in the mouse. To allow cross-species analysis of primate Betz cells and mouse ET neurons, we first created a joint embedding of excitatory neurons in mouse, macaque and human, which showed strong homology across all three species for the L5 ET subclass (**Fig. 4h**). Patch-seq recordings were made from L5 neurons in acute and cultured slice preparations of mouse MOp and macaque M1. We also capitalized on a unique opportunity to record from neurosurgical tissue excised from the human premotor cortex, which also contains Betz cells, during an epilepsy treatment surgery. To permit visualization of cells in heavily myelinated macaque M1 and human premotor cortex, AAV viruses were used to drive fluorophore expression in glutamatergic neurons in slice culture.

Patch-seq cells in each species that mapped to the L5 ET subclass (**Fig. 4h**) were all large layer 5 neurons that sent apical dendrites to the pial surface (**Fig. 4i**, note truncation in human Betz cell). However, macaque and human L5 ET neurons were much larger, and had long “tap root” basal dendrites that are a canonical hallmark of Betz cells ^83^. Subthreshold membrane properties were relatively well conserved across species. For example, L5 ET neurons in all three species had a low input resistance, although it was exceptionally low in macaque and human (**Fig. 4j**). Conversely, suprathreshold properties of macaque and human Betz/ET neurons were highly specialized. Most notably, human and macaque neurons responded to prolonged suprathreshold current injections with a biphasic firing pattern in which a pause in firing early in the sweep was followed by a dramatic increase in firing late in the sweep (**Fig. 4k**). Intriguingly, we identified several genes encoding ion channels that were enriched in macaque and human L5 ET neurons compared with mouse (**Fig. 4l**). These primate specific ion channels may contribute to the distinctive suprathreshold properties of primate ET neurons. Together this indicates that primate Betz cells are homologous to mouse thick-tufted L5 ET neurons, but display phenotypic differences in their morphology, physiology and gene expression. Similar to transcriptomics, these results indicate strong conservation of cell subclasses but with significant species specializations in anatomical and functional properties.

### Multimodal correspondence by Epi-Retro-Seq

To obtain a comprehensive view of the molecular diversity among projection neurons in MOp, we developed Epi-Retro-Seq (companion paper ^79^) and applied it to mouse MOp neurons that project to each of the 8 selected brain regions that receive inputs from MOp (**Fig. 5a**). The target regions included two cortical areas, SSp and anterior cingulate area (ACA), and six subcortical areas, striatum (STR), thalamus (TH), superior colliculus (SC), ventral tegmental area and substantia nigra (VTA+SN), pons, and medulla (MY). Specifically, we injected the retrograde tracer rAAV2-retro-Cre ^77^ into the target region in INTACT mice ^85^, which turned on Cre-dependent GFP expression in the nuclei of MOp neurons projecting to the injected target region. Individual GFP-labeled nuclei of MOp projection neurons were then isolated using fluorescence-activated nucleus sorting (FANS). Single-nucleus methylcytosine sequencing (snmC-Seq2) ^49^ was performed to profile the DNA methylation (mC) of each single nucleus.

**Figure 5.**
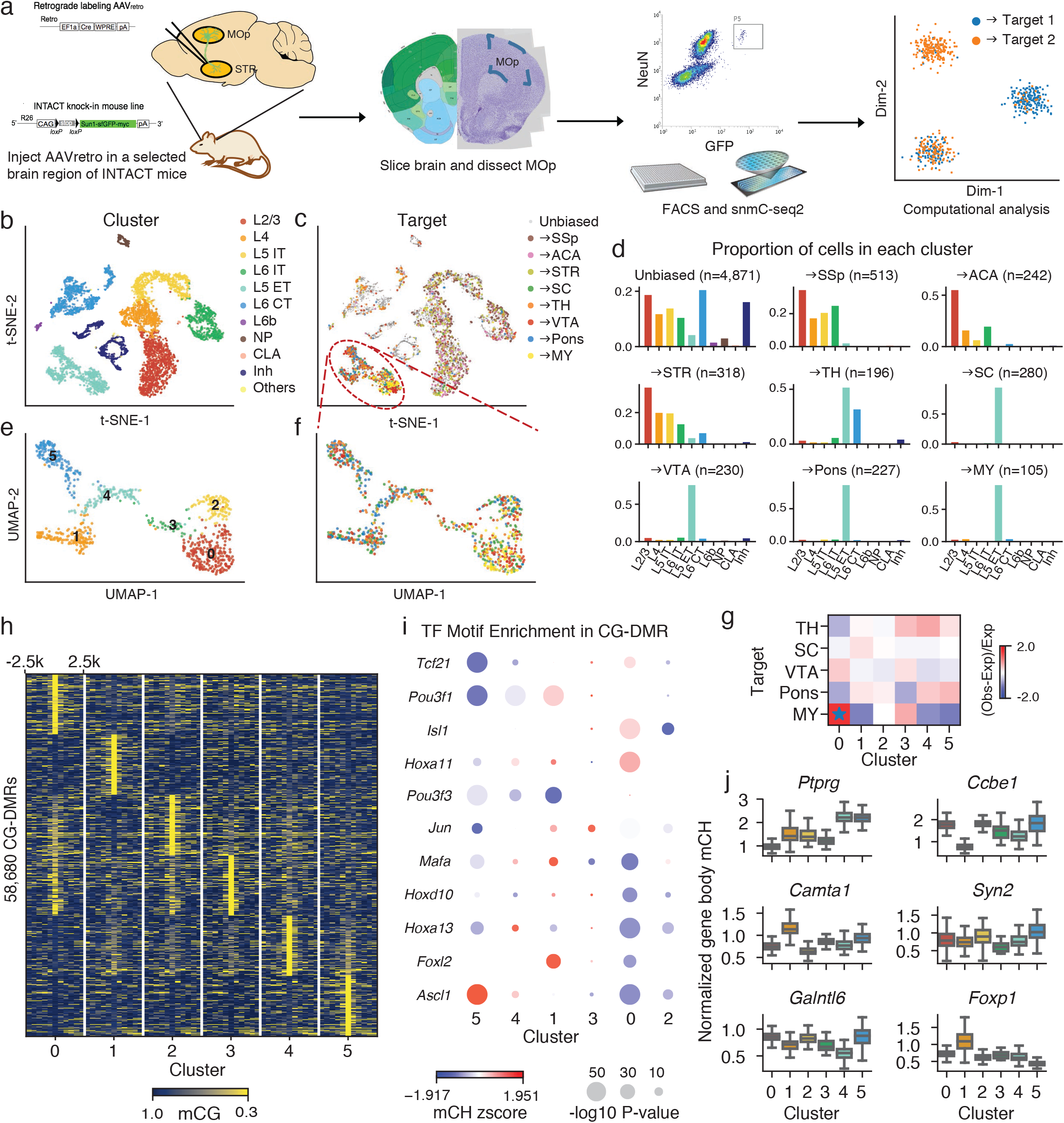
Epi-Retro-Seq links molecular cell type with distal projection targets. **a,** Workflow of Epi-Retro-Seq. **b, c,** UMAP embedding of MOp cells profiled by Epi-Retro-Seq (n=2,115) and unbiased snmC-Seq2 (n=4,871) computed with 100kb-bin-level mCH, colored by subclasses (**b**) or projection targets (**c**). **d,** Distribution across subclasses of neurons from unbiased snmC-Seq2 and neurons projecting to each target. **e, f,** UMAP embedding of L5 ET cells in MOp profiled by Epi-Retro-Seq (n=848) computed with 100kb-bin-level mCH, colored by clusters (**e**) or projection targets (**f**). **g,** Enrichment of L5 ET neurons projecting to each target in each cluster. * represents FDR<0.05. **h,** mCG levels at CG-DMRs identified between the six clusters and their flanking 2.5k regions. Top 100 DMRs in each cluster were shown. **i,** TF motif enrichment in CG-DMRs in each cluster. Color represents z-scored gene-body mCH level of the TFs, and size represents -log10 *P* value of motif enrichment in the CG-DMRs. **j,** Boxplots of normalized mCH levels at gene-bodies of example CH-DMGs in the six clusters. Numbers of cells represented by the boxes are 242, 165, 118, 42, 119, and 162 for the six clusters. The elements of boxplots are defined as: center line, median; box limits, first and third quartiles; whiskers, 1.5× interquartile range.

After removing low-quality cells, potential doublets, and non-neuronal cells, we obtained high-quality methylomes for 2,111 MOp projection neurons. When co-clustering them with MOp neurons collected without enrichment of specific projections, we observed a precise agreement among all of the major cell subclasses (**Fig. 5b,c**), demonstrating the robustness of Epi-Retro-Seq to classify cell types. Although neurons projecting to different target regions were not completely separated on t-SNE, we observed the explicit enrichment of cortico-cortical and cortico-striatal projecting neurons in IT subclasses (L2/3, L4, L5 IT, L6 IT, and L6 IT Car3), and cortico-subcerebral projecting neurons in L5 ET. Many cortico-thalamic projecting neurons were also observed in L6 CT subclass (**Fig. 5d**). These observations are consistent with the known laminar distribution of the cortico-cortical and cortical-subcortical projection neurons ^81^, reflecting the high quality of retrograde-labeling of neuronal nuclei in our Epi-Retro-Seq dataset.

The enrichment of L5 ET neurons in the Epi-Retro-Seq data (40.2% vs. 5.62% in unbiased profiling of MOp using snmC-seq2) allowed a more detailed investigation of the subtypes of L5 ET neurons which are known to project to multiple subcortical targets in TH, VTA+SN, pons and MY ^81^. The 848 L5 ET neurons further segregated into 6 clusters (**Fig. 5e,f**). MY-projecting neurons showed a clear enrichment in L5 ET cluster 0 (**Fig. 5f,g**), in agreement with scRNA-Seq data for anterolateral motor cortex (ALM), part of MOs ^15,86^. We used gene body non-CG methylation (mCH) levels to integrate the L5 ET Epi-Retro-Seq data with the ALM Retro-seq data and also observed the enrichment of MY-projecting cells in the same cluster ^79^.

A major advantage of DNA methylation profiling of neurons is its ability to obtain information for both genes and cis-regulatory elements. Specifically, mCH at gene bodies is strongly anti-correlated with gene expression in neurons, while promoter-distal differentially CG-methylated regions (CG-DMRs) are reliable markers of regulatory elements such as enhancers ^20^. We thus identified 511 differentially CH-methylated genes (CH-DMGs) and 58,680 CG-DMRs across the L5 ET clusters (**Fig. 5h**). We also inferred transcription factors (TFs) that may contribute to defining the cell subclusters by identifying enriched TF-binding DNA sequence motifs within CG-DMRs (**Fig. 5i**). For example, *Ascl1* is a transcription factor whose motif was significantly enriched in the MY-projecting cluster. Previous studies had shown its necessity for neuronal differentiation and specification in multiple regions of the nervous system ^87,88^. In addition, 230 hypo-CH-DMGs were identified between the MY-projecting cluster and other projection neurons. Interestingly, one of the most differentially methylated genes is *Ptprg* (**Fig. 5j**), which interacts with contactin proteins to mediate neural projection development ^89^.

In summary, Epi-Retro-Seq mapping data for MOp revealed specific enrichment of MY-projecting neurons in one of the molecularly-defined subpopulations of MOp L5 ET neurons, allowing identification of regulatory elements for this unique cell type. In addition to MOp, we have performed 63 Epi-Retro-Seq mapping experiments for 7 cortical regions, comprising 26 cortico-cortical projections and 37 cortico-subcortical projections ^79^. Together, these epigenomic mapping data for projection neurons facilitates the understanding of gene regulation in establishing neuronal identity and connectivity, by discovering projection-specific gene regulatory elements which can be used to target specific types of projection neurons.

### MOp projection neuron types and input-output wiring diagram

Building upon the molecularly defined and spatially resolved cell atlas (**Fig. 3**) and the multi-modal correspondence between gene expression and morpho-electric properties of MOp neurons (**Fig. 4**), we next describe a comprehensive cellular resolution input-output MOp wiring diagram. To achieve this, we combined classic tracers, genetic viral labeling in Cre driver lines and single neuron reconstructions with high-resolution, brain-wide imaging, precise 3D registration to CCF, and computational analyses (companion paper ^69^).

First, we systematically characterized the global inputs and outputs of MOp upper limb (MOp-ul) region using classic anterograde (PHAL) and retrograde (CTb) tract tracing ^69^ (**Fig. 6a**). At the macro-scale, MOp-ul projects to more than 110 gray matter regions and cervical spinal cord, and ∼60 structures in the cerebral cortex and thalamus project back to MOp-ul.

**Figure 6.**
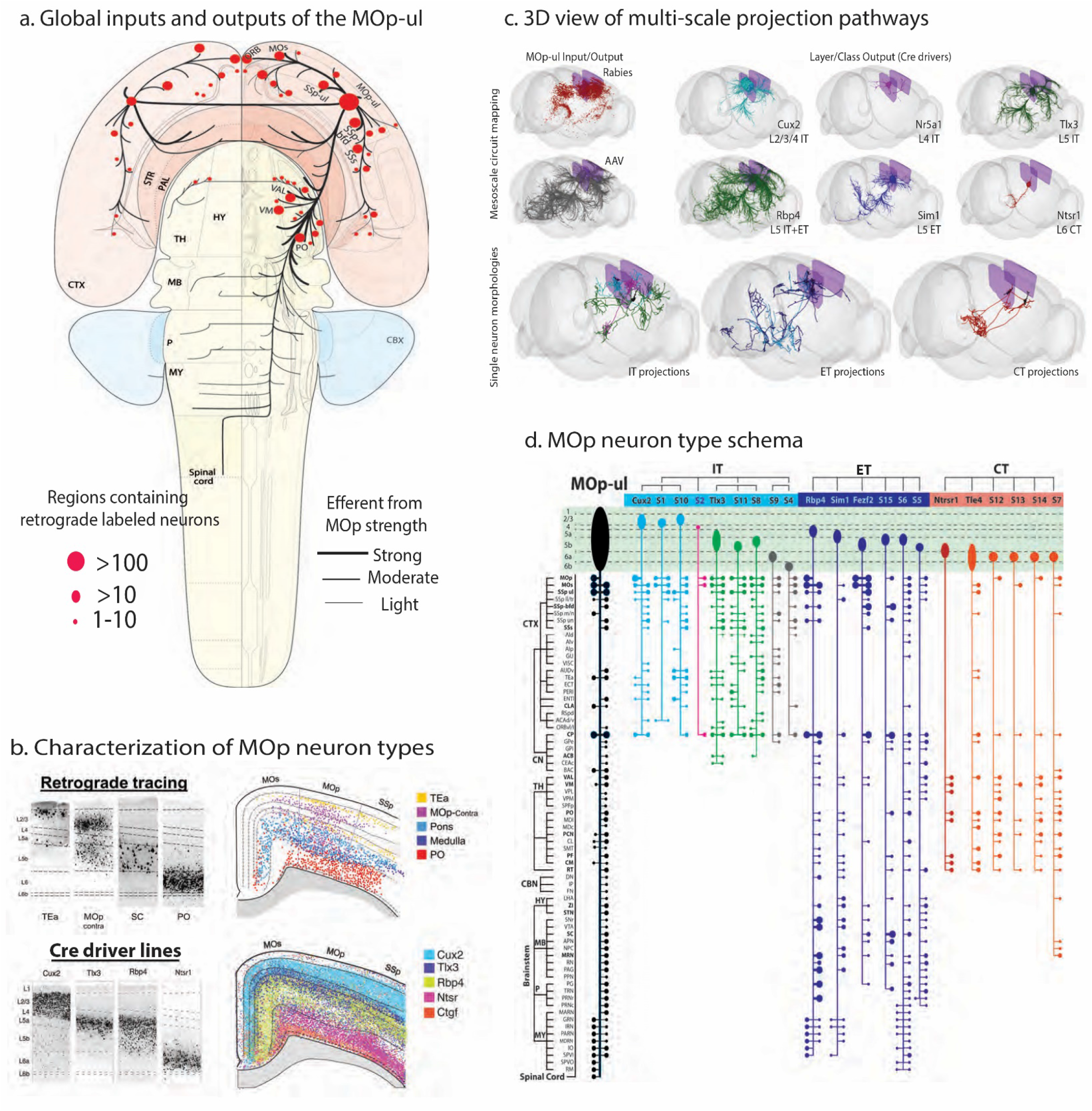
Global wiring diagram and anatomical characterization of MOp-ul neuron types. **a,** Flatmap representation of the MOp-ul input/output wiring diagram. Black lines and red dots indicate axonal projections (outputs) and retrograde labeling sources (inputs), respectively, with line thickness and dot sizes representing relative connection strengths. Most MOp-ul projection targets in the cortex and thalamus also contain input sources, suggesting bi-directional connections. The flatmap is adapted from the Swanson Brainmap 4.0 and the mouse brain flatmap ^92^. **b,** MOp-ul neurons classified by projection targets or transgenic *Cre* expression. (Top) Retrograde tracing using CTB revealed layer-specific distributions of MOp-ul neurons with respect to their major projection targets. Representative images (left) show neurons labeled by CTB injections into cortical areas (TEa, contralateral MOp), superior colliculus (SC) in the midbrain, and posterior complex (PO) of the thalamus. Detected cells were pseudo-colored and overlaid onto a schematic coronal section near the center of MOp-ul (right). MOp neurons that project to TEa are distributed in L2 and L5 (yellow), to the contralateral MOp in L2-L6b (purple), to targets in the pons and medulla in L5b (blue), and to thalamus in L6a (red). (Bottom) The distribution of neurons labeled in 28 transgenic *Cre* lines was mapped in MOp and across the whole cortex. Images (left) show laminar patterns of Cre+ nuclei in MOp-ul from four driver lines (*Cux2*, *Tlx3*, *Rbp4*, and *Ntsr1*). Detected nuclei from these lines, plus the *Ctgf-Cre* line, were pseudo-colored and overlaid onto a schematic coronal section near the center of MOp-ul (right). Cre+ nuclei are found in L2-4 in Cux2; L5a and superficial L5b in Tlx3; L5a and L5b in Rbp4; L6a in Ntsr1, and L6b in Ctgf. **c,** 3D views show brain-wide MOp input-output patterns at the population and single cell resolution. (Top left) Regional MOp inputs and outputs were mapped using retrograde (in red, example shows rabies tracing from the *Tlx3-Cre* driver line) and anterograde (in black, example shows AAV-EGFP) tracing methods. (Top right) Whole-brain axonal trajectories from 6 Cre line-defined subpopulations labeled with Cre-dependent AAV tracer injections at the same MOp-ul location. (Bottom) Individual projection neurons were fully reconstructed following high-resolution whole-brain imaging of sparsely labeled cells. Representative examples of IT, ET, and CT neurons are shown in each panel. The two ET examples represent distinct projection-types; medulla (dark blue)- and non-medulla-projecting (light blue). 3D renderings were generated following registration of projection and reconstruction data into CCFv3 using BrainRender ^93^. **d,** Projection patterns arising from major cell types, IT, ET and CT, with corresponding Cre-line assignment and somatic laminar location, compared with the overall projection pattern from the MOp-ul region (left, black). Along each vertical output pathway, horizontal bars on the right and left sides represent ipsilateral and contralateral collaterals, respectively, with dot sizes indicating the strength of axonal terminals in different targets. Brain structure nomenclature adopted from ARA ^94^.

Next, we generated a fine-grained areal and laminar distribution map of multiple MOp-ul projection neuron populations using retrograde pathway-tracing. Accordingly, we identified 25 distinct neuron projection types based on their unique combinations of axonal targets and laminar somatic distributions (**Fig. 6b**, top; for details see ^69^). For example, IT cells (e.g. TEa-targeting or contralateral MOp-targeting) are distributed throughout L2-L6b; ET cells (pons- or medulla-targeting) are distributed primarily in L5b and most CT (posterior thalamic nucleus-targeting) neurons are distributed in L6a.

In parallel with these tracer-labeled, projection- and layer-defined cell types, we quantitatively characterized the distribution patterns of neuronal subpopulations in the MOp-ul labeled in 28 Cre-expressing “driver” lines (**Fig. 6b**, bottom). These lines selectively label neurons from different IT (e.g. Cux2, Plxnd1, Tlx3), L5 ET (Rbp4, Sim1, Fezf2), and CT (Ntsr1, Tle4) subpopulations with distinct laminar distributions ^75,90,91^.

Subsequently, we used viral tracers to systematically examine MOp-ul cell-type-specific inputs and outputs (**Fig. 6c**). First, neurons projecting *to* Cre-defined starter cells were labeled using transsynaptic rabies viral tracing methods; an example from the *Tlx3* L5 IT line is shown in **Fig. 6c** (upper left, red). Projections *from* MOp were labeled following AAV-GFP injections into C57BL6/J mice, revealing patterns consistent with PHAL tracing results (**Fig. 6a**). Projections from L2/3 IT, L4 IT, L5 IT, L5 ET, and L6 CT cells were mapped following injections of Cre-dependent viral tracers into Cre lines selective for these laminar- and projection-cell subclasses ^71^. Most Cre line anterograde tracing experiments revealed a component of the overall output pathway (**Fig. 6c**). For example, the L6 *Ntsr1* line revealed a typical CT projection pattern with dense projections specific to thalamic nuclei. This result is consistent with labeling from retrograde injections in various thalamic nuclei (PO, VAL, VM) and cortical areas such as MOs and SSp (**Fig. 6b**, top). Further characterization of the distinctive projection patterns of several IT, L5 ET, and CT driver lines is provided in the anatomy companion paper ^69^.

To further refine the projection neuron characterization, we carried out single cell analysis by combining sparse labeling, high-resolution whole-brain imaging, complete axonal reconstruction and quantitative analysis (companion papers ^68,69^); additional analysis was also conducted using BARseq ^69^, a high-throughput projection mapping technique based on *in situ* sequencing ^67^. We augmented the full morphology reconstruction dataset with publicly available single cell reconstructions in MOp from the Janelia Mouselight project ^26^. We systematically characterized axonal projections of 151 single MOp pyramidal neurons. This analysis revealed a rich diversity of projection patterns within the IT, ET and CT subclasses (**Fig. 6c,d**). For example, individual L6 neurons display several distinct axonal arborization targets that likely contribute to the composite subpopulation output described for the *Ntsr1* and *Tle4* diver lines (**Fig. 6d**). Confirming and extending previous reports ^86^, we characterized detailed axonal trajectories and terminations of two major types of L5b ET cells, namely medulla-projecting and non-medulla projecting neurons; both types may collateralize in the thalamus and terminate in the midbrain (**Fig. 6d**). Individual IT cells across L2-L6 also generate richly diverse axonal trajectories (detailed in ^68,69^. Further analyses of complete single neuron morphologies, precisely registered in the CCF, will provide the ultimate resolution toward defining anatomical cell types and clarify the anatomical heterogeneity described at the subpopulation level.

In summary, combining multiple approaches complementary in their coverage, throughput, and resolution, we provide a comprehensive identification of major projection neuron types with correspondence to molecular markers. We further delineate their input-output patterns at the subpopulation level and describe projection patterns at single-cell resolution, deriving the first multi-scale wiring diagram of MOp. A major future goal is to link these anatomic and especially projection types with transcriptomic types (**Fig. 2b**), with precise registration to a spatial atlas (e.g. **Fig. 3e**).

### Cell Type Targeting Tools

The identification and classification of MOp cell types based on single-cell integration of transcriptomes and epigenomes (**Fig. 2**), spatially resolved single-cell transcriptomics (**Fig. 3**) and anatomical and physiological analysis (**Fig. 4–6**) provides deep insights into the molecular basis of cellular diversity. In addition to establishing a principled basis for a taxonomy of brain cell types, knowledge of cellular gene expression also provides information to create mouse models in which genetically encoded reporters and actuators are targeted to these molecularly defined cell types ^33^.

As an embodiment of this approach, we used CRISPR/Cas-9-mediated homologous recombination in ES cells to generate genetically modified mice (Stafford, Daigle, Chance et al., companion manuscript in preparation) in which sequences encoding FlpO and Cre recombinases were targeted respectively to *Npnt* and *Slco2a1*, genes whose differential expression discriminates between two types of L5 ET neurons with distinct subcortical projection target specificities ^15,86^. Confirming the assignment of *Npnt*- and *Slco2a1*-expressing cells to subsets of L2/3 IT and L5 ET neurons in the consensus transcriptomic taxonomy (**Fig. 7a**), FlpO- and Cre-dependent tdTomato reporter expression in *Npnt-P2A-FlpO;Ai65F* and *Slco2a1-P2A-Cre;Ai14* mice localized to these cortical cell layers in MOp (**Fig. 7b**). In *Npnt* mice, both L2/3 and L5 neurons were labeled. In *Slco2a1* mice, predominantly L5 neurons were labeled. It is noteworthy that *Slco2a1* labeled cells occupying a deeper sub-lamina of L5 than those targeted by *Npnt*, in accord with a previous report describing the two types of L5 ET neurons ^86^ (see also **Fig. 9** below). To test the projection specificity of neurons labeled by these novel genetic tools, we injected a recombinant AAV encoding a Cre-dependent EGFP reporter into deep L5 in MOp of a *Slco2a1-P2A-Cre* mouse (**Fig. 7c**). Consistent with previous studies ^86^ as well as those described in **Figures 5, 6 and 9** (below), GFP-labeled axon terminals were found in pontine gray and medulla, indicating that this mouse line labels the medulla-projecting L5 ET cell type.

**Figure 7.**
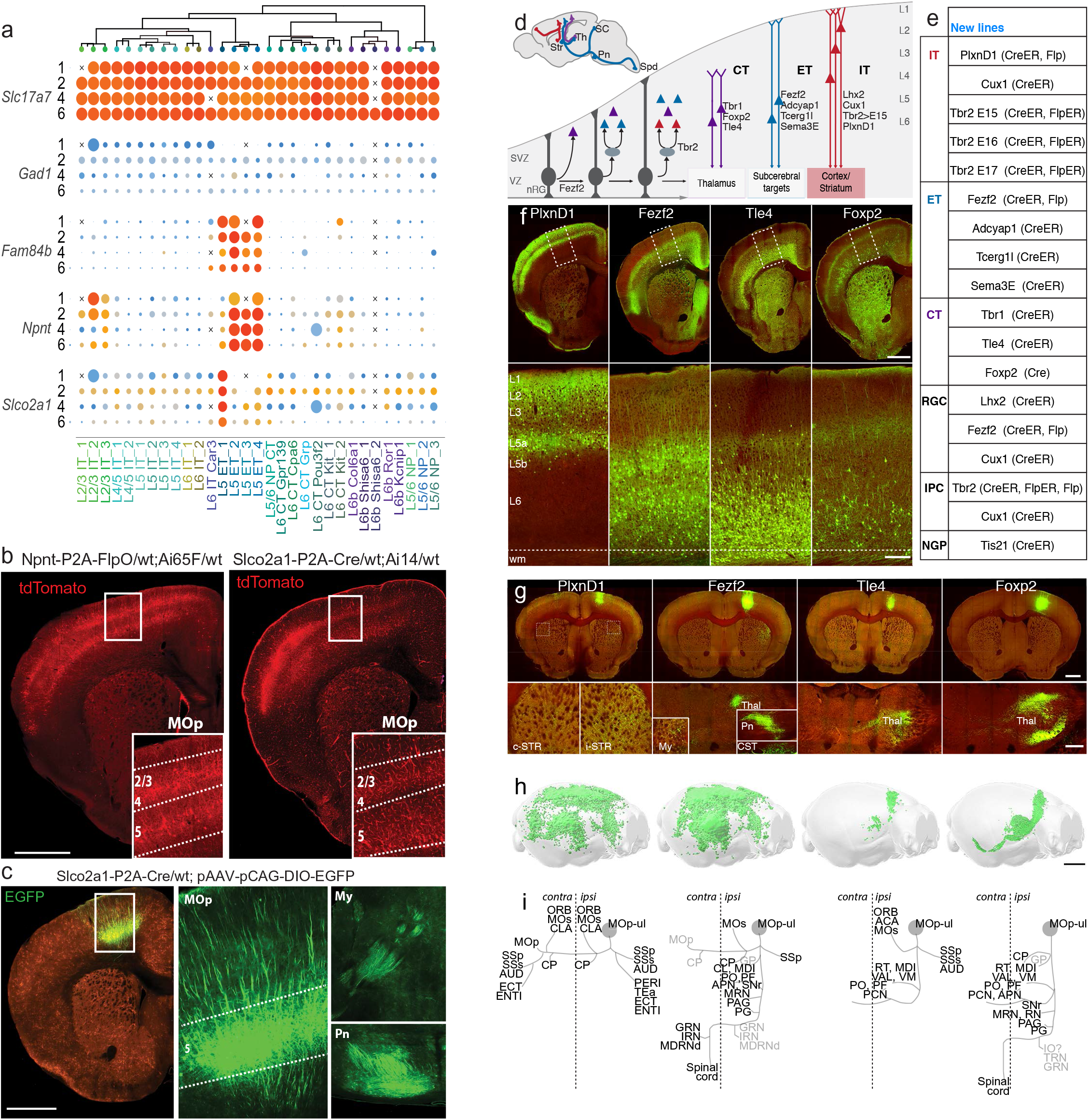
Genetic tools for targeting cortical glutamatergic projection neuron types. **a,** Dendrogram of excitatory cells types within the MOp transcriptomic taxonomy followed by the proportion of cells (dot size) expressing pan-excitatory or -inhibitory marker genes (*Slc17a7* or *Gad1*), or L5 ET marker genes (*Fam84b*, *Npnt*, and *Slco2a1*). Expression results from four different single cell RNA-seq platforms are shown: 1: scRNA-seq SMART-Seq; 2: scRNA-seq 10X v3 A; 4: snRNA-seq SMART-Seq; 6: snRNA-seq 10X v3 B ^45^. **b,** Representative images of native tdTomato fluorescence from MOp of *Npnt-P2A-FlpO;Ai65F* and *Slco2a1-P2A-Cre;Ai14* animals. Reporter expression was observed in L2/3 and L5 neurons with both driver lines and in the vasculature with only the Slco2a1 line. **c,** Representative images of native EGFP fluorescence from MOp, My (medulla), and Pn (pons) in the brain of an *Slco2a1-P2A-Cre* animal injected in MOp with a Cre-dependent reporter AAV (*pCAG-FLEX-EGFP-WPRE*). Robust reporter expression in L5 neurons was observed at the injection site (MOp) and in fibers terminating in My and Pn. **d,** Schematic (upper left panel) depicting several major pyramidal neuron (PyN) projection classes that mediate intra-telencephalic streams (IT-red; cortical and striatal) and cortical output channels (ET-blue, CT-purple). Str, striatum; Th, thalamus; SC, superior colliculus; Spd, spinal cord. Developmental trajectory of PyNs (lower panel) depicting lineage progression from progenitors to mature PyNs across major laminar and projection types. Genes used to target progenitor and PyN subpopulations are listed according to their cellular expression patterns. VZ, ventricular zone; SVZ, subventricular zone. **e,** Table presenting new gene knockin driver mouse lines targeting PyN progenitors and projection types. RGC, radial glia cell; IPC, intermediate progenitor cell; NGP, neurogenic progenitor. **f,** Cre recombination patterns visualized through reporter expression (green) and background autofluorescence (red) from four driver/reporter lines *PlexinD1-2A-CreER* (*PlxnD1*);*Snap25-LSL-EGFP, Fezf2-2A-CreER* (*Fezf2*)*;Ai14, Tle4-2A-CreER* (*Tle4*);*Snap25-LSL-EGFP* and *Foxp2-IRES-Cre* (*Foxp2*);*AAV9-CAG-FLEX-EGFP* (systemic injection). Top row: coronal hemisections containing MOp. Bottom row: a segment of MOp (dashed lines, top row) with laminar delineations. *CreER* Tamoxifen (TM) inductions were at P21 and P28. **g,** Anterograde tracing from PyN subpopulations in MOp. CreER drivers were crossed with a *Rosa26-CAG-LSL-Flp* mouse, and postnatal TM induction to convert to constitutive Flp expression for anterograde tracing with a Flp-dependent AAV vector expressing EGFP (*AAV8-CAG-fDIO-TVA-EGFP*). Representative images of native EGFP fluorescence from the MOp injection site (top row) from cell-type-specific viral vector (green) and background autofluorescence (red) at selected subcortical projection targets for four driver lines: Th; Str; cerebral peduncle (cp), Pn, My and corticospinal tract (CST). **h,** Whole-brain three dimensional renderings of axon projections registered to the CCFv3 for each PyN subpopulation in the MOp cortex (parasagittal view). **i,** Schematics of main projection targets for each PyN subpopulation. Vertical dashed line indicates midline; filled circle indicates MOp injection site. Scale bars: hemisections (f & g) and h, 1mm; bottom row in f, 200μm; bottom row in g, 500μm; h, 2 mm.

**Figure 9.**
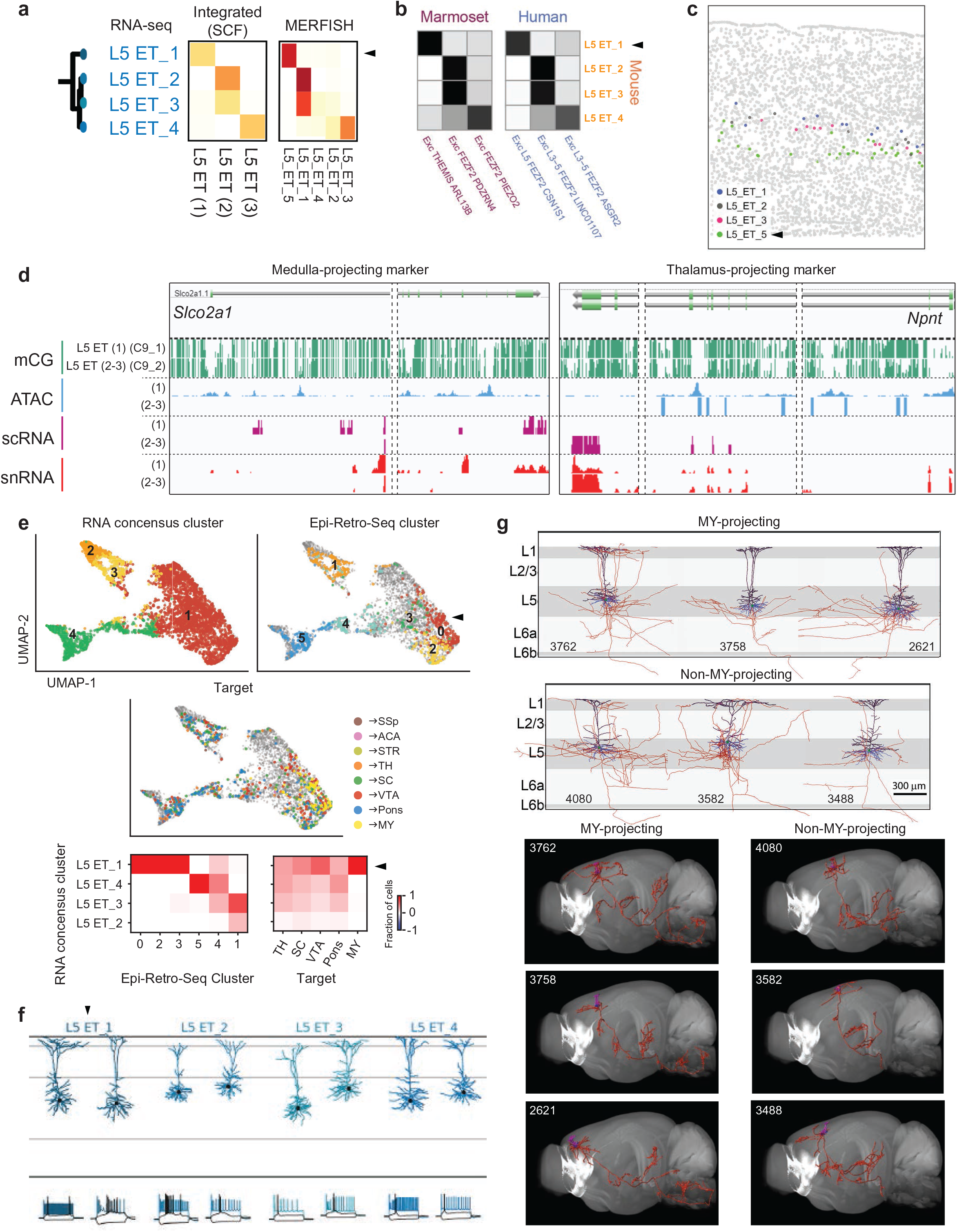
Two distinct L5 ET projection neuron types in MOp. **a**, Within the mouse L5 ET subclass, good correspondence is observed between the 4 transcriptomic clusters and the 3 integrated molecular clusters (SingleCellFusion) or the 5 MERFISH clusters. **b**, Within the L5 ET subclass, the 4 mouse transcriptomic clusters correspond well with the 3 transcriptomic clusters in either human or marmoset. **c**, In MERFISH, cells belonging to the L5_ET_1-4 clusters co-occupy the upper L5, whereas L5_ET_5 cells are distinctly located in lower L5. **d**, Genome browser of gene markers between the MY-projecting (*Slco2a1*) and the non-MY-projecting (*Npnt*) L5 ET neurons. **e**, Integration panels between L5 ET Epi-Retro-Seq clusters and consensus transcriptomic clusters. The transcriptomic dataset used here is snRNA 10x v3 B, which has the largest number of L5 ET cells (>4k). Top panels, UMAP plots colored by consensus transcriptomic clusters, Epi-Retro-Seq clusters and projection targets (retrograde tracer injection sites). Bottom panels, confusion matrices between consensus transcriptomic clusters and Epi-Retro-Seq clusters or major projection targets. The heatmaps are column-wise normalized rather than row-wise to avoid misleading interpretation, since the number of cells sampled from each projection may differ a lot in Epi-Retro-Seq. **f**, Dendritic morphologies and spiking patterns of representative Patch-seq cells corresponding to the 4 mouse transcriptomic L5 ET types. **g**, Local dendritic and axonal morphologies (upper panels) and brain-wide axon projections (lower panels) of representative fully-reconstructed L5 ET neurons, separated into MY-projecting and non-MY-projecting types. Black, apical dendrites. Blue, basal dendrites. Red, axons.

To expand on cell type driver lines, we further built a genetic toolkit for cortical pyramidal neurons (PyNs) with more comprehensive coverage of projection types and with combinatorial strategies for improved specificity (companion paper ^75^). First, we generated and characterized a set of 15 Cre and Flp gene knockin mouse driver lines for targeting major PyN subpopulations and progenitor types, guided by knowledge in their gene expression as well as developmental genetic programs (**Fig. 7d,e**). These include the broad CT (*Tbr1, Tle4, Foxp2*), ET (*Fezf2, Adcyap1, Tcerg1l, Sema3e*) and IT (*Plxnd1, Cux1,* and *Tbr1* late embryonic inductions) subclasses as well as subpopulations within these subclasses. When crossed with reporter alleles, these driver lines activated reporter expression that precisely recapitulated endogenous expression patterns highlighted here with 4 representative lines (**Fig. 7f**): L2/3 and L5a for IT-*Plxnd1* (IT*^Plxnd^*^1^), L5b and L6 for ET-*Fezf2* (ET*^Fezf2^*), L6 for CT-*Tle4* and CT-*Foxp2* (CT*^Tle4^*, CT*^Foxp2^*). To examine the projection pattern of these driver-defined subpopulations, we converted inducible CreER expression to constitutive Flp expression followed by MOp injection of a Flp-dependent AAV reporter vector (**Fig. 7g-i**). Largely as expected, IT*^Plxnd1^* projected to multiple ipsi- and contra-lateral cortical areas and the striatum/caudate putamen; ET*^Fezf2^* projected robustly to several ipsi-lateral cortical sites, striatum, and numerous subcortical targets including thalamus, medulla and the corticospinal tract; CT*^Tle4^* projected to a set of highly specific thalamic nuclei. Surprisingly, CT*^Foxp2^* projected to a set of specific thalamic nuclei as well as to midbrain, brainstem and corticospinal tract. Further characterization of this set of new driver lines (**Fig. 7e**) is presented in ^75^.

Together, these tools and strategies establish an experimental approach for accessing hierarchically organized neuronal cell types at progressively finer resolution. Such genetic access will enable an integrated multi-modal analysis to further validate and characterize these cell populations as well as to explore their multi-faceted function in neural circuit operation and behavior.--

### Integrated multimodal characterization reveals L4 IT neurons in MOp

To investigate if our collective multimodal characterization can lead to an integrated understanding of cell types in MOp, we selected two case studies to demonstrate convergence of multiple corresponding properties onto specific cell types.

Traditionally MOp has been considered an agranular cortical area, defined by the lack of a cytoarchitectonic layer 4 which usually contains spiny stellate or star pyramid excitatory neurons. However, a previous study challenged this view and presented evidence that L4 neurons similar to those typically found in sensory cortical areas also are present in MOp ^95^. Here as the first case study, we used multimodal evidence to confirm the presence of L4-like neurons in mouse MOp and possibly in primate M1 as well (**Fig. 8**).

**Figure 8.**
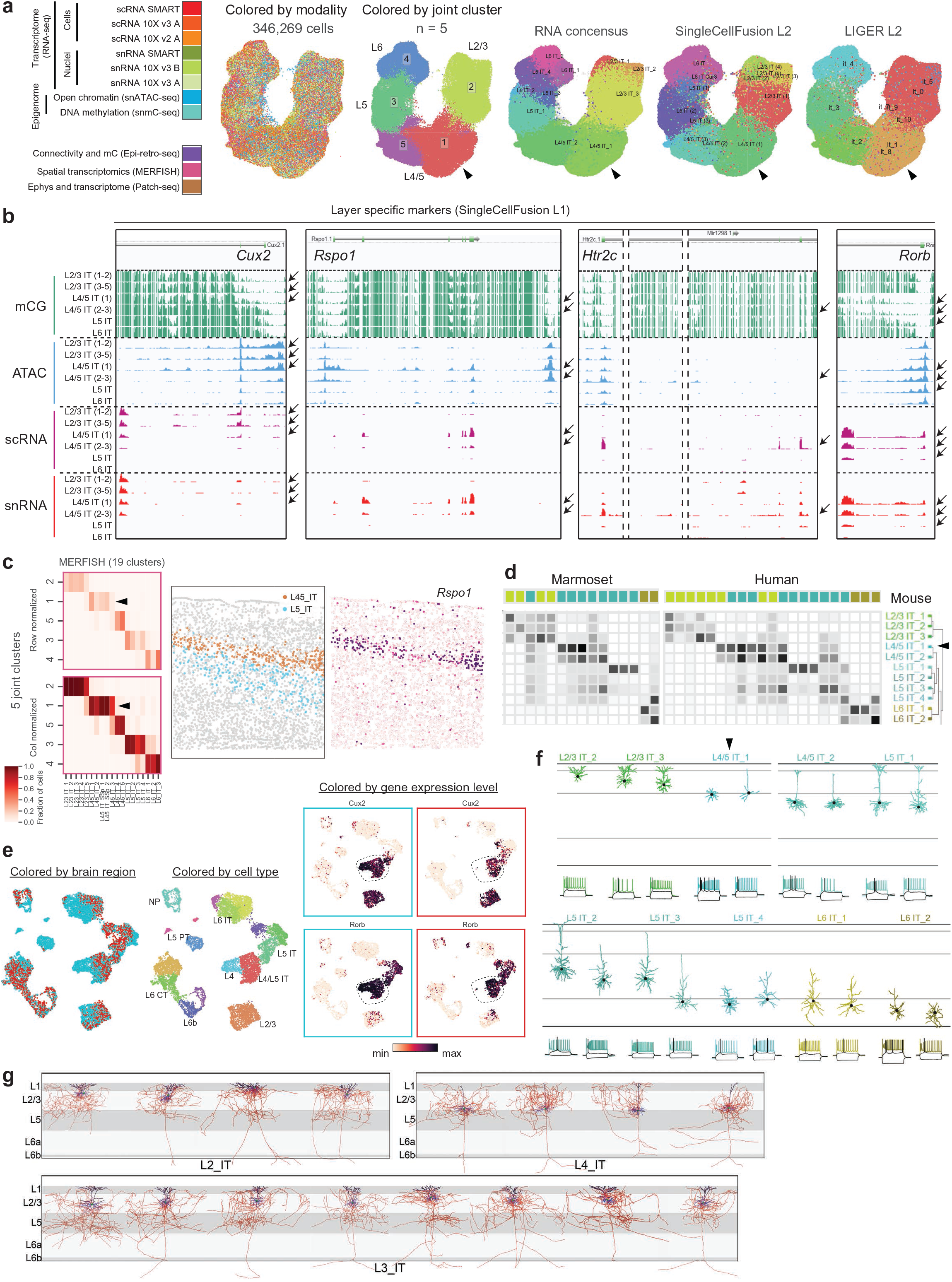
Existence of L4 excitatory neurons in MOp. **a**, UMAP embedding of IT cells from 11 datasets. Cells are colored by modalities, by cluster identities from the 11-dataset joint clustering, and by cluster identities generated from other consensus clustering methods in ^45^. **b**, Genome browser of layer-specific gene markers -from L2/3 to L5 - across IT cell types as defined in ^45^ (SingleCellFusion L1). Arrows indicate cell types with correlated transcription and epigenomic signatures of the specific marker gene. **c**, MERFISH IT clusters correspond well with the above joint clusters from **a** (confusion matrices, left panel) and reveal a L4 specific cluster (L45_IT) distinctly separated from the L5_IT cluster (middle panel) and marked by gene *Rspo1* (right panel). **d**, Correspondence between mouse and human or marmoset transcriptomic IT types. **e**, UMAP embedding of excitatory cells from MOp (scRNA_SMART) ^45^ and VISp ^15^ show that L4 excitatory cells from MOp correspond to a subset of L4 excitatory cells from VISp. Cells are colored by brain regions (MOp, red; VISp, blue), by cell types, and by expression levels (log10(TPM+1)) of marker genes *Cux2* and *Rorb*. **f**, Dendritic morphologies and spiking patterns of representative Patch-seq cells from all IT types (L2/3 to L6). Arrow heads in a, c, d and f indicate the L4/5 IT 1 type. **g**, Local dendritic and axonal morphologies of fully reconstructed IT neurons with somas located in L2, L3 and L4. Black, apical dendrites. Blue, basal dendrites. Red, axons.

We first performed a joint clustering (see Methods) and UMAP embedding of all IT cells (excluding the highly distinct L6 IT Car3 cells) from 11 different mouse molecular datasets, including 6 sc/snRNA-seq datasets, and the snmC-Seq2, snATAC-Seq, Epi-Retro-Seq, MERFISH and Patch-seq data (**Fig. 8a**). This resulted in 5 joint clusters, mostly along a continuous variation axis moving from L2/3 to L4/5 to L5 to L6. The joint clustering enabled linkage of the cells independently profiled by each individual modality into types - transcriptomic, epigenomic, spatially resolved transcriptomic, and morpho-electric-transcriptomic, and cross-correlation of these disparate properties. Consequently, we identified epigenomic peaks linked to cluster-specific marker genes - *Cux2* for L2/3 IT and L4/5 IT (1), *Rspo1* for L4/5 IT (1), *Htr2c* for L4/5 IT (2-3), and *Rorb* for L4/5 IT and L5 IT (**Fig. 8b**, cluster names from SingleCellFusion). MERFISH data also showed that L4/5 IT and L5 IT cells occupied distinct layers, and the L4/5 IT type expressed *Rspo1* (**Fig. 8c**), a L4 cell type marker in sensory cortical areas identified in previous studies ^15^. Transcriptomic IT types from mouse corresponded well with those from human and marmoset, but such correspondence was mostly at main branches or subclass level while significant confusions existed at single cluster level (**Fig. 8d**), likely due to the substantial gene expression variation between rodents and primates (**Fig. 2**). In particular, mouse L4/5 IT 1 and 2 transcriptomic clusters together corresponded to a set of 5-7 L3-5 IT clusters in human and marmoset.

We further compared the L4-like cells in mouse MOp with those from mouse primary visual cortex (VISp) ^15^ after co-clustering all the SMART-Seq glutamatergic neurons from both regions (**Fig. 8e**). In the UMAP representation, L4/5 IT cells in MOp occupied a subspace of the L4 IT co-cluster defined by the intersection of marker genes *Cux2* and *Rorb*, suggesting that L4-like cells in MOp are similar to a subset of L4 cells in VISp while the L4 cells in VISp have additional diversity and specificity.

L4-like IT cells in MOp also exhibited morphological features characteristic of traditionally defined L4 excitatory neurons. From the Patch-seq study ^64^, cells from the L4/5 IT_1 type had no or minimal apical dendrites without tufts in layer 1, in contrast to cells from the neighboring L2/3 IT, L4/5 IT_2 and L5 IT types which had tufted apical dendrites (**Fig. 8f**). We also obtained full morphological reconstructions of excitatory neurons with their somas located in L2, L3 or L4 in MOp or the neighboring secondary motor area (MOs) from fMOST imaging of *Cux2-CreERT2;Ai166* mice ^68,80^. As shown in **Fig. 8b**, *Cux2* is a specific marker gene for L2/3 IT and L4/5 IT_1 types. These full reconstructions allowed us to examine, in addition to dendritic morphologies, the full extent of both local and long-range axon projections. The MOp/MOs neurons with somas in putative L4 (between L2/3 and L5) exhibited two local morphological features characteristic of L4 neurons found in sensory cortical areas (**Fig. 8g**). First, the dendrites of the L4 neurons were simple and untufted whereas those of the L2 and L3 neurons all had extensive tufts. Second, the local axons of L4 neurons mostly projected upward into L2/3 in addition to collateral projections; on the contrary, the local axons of L2 and L3 neurons mostly projected downward, reaching into L5. These local projection patterns are consistent with the canonical feedforward projections within a cortical column observed in somatosensory and visual cortices, with the first feedforward step being from L4 to L2/3 and the second feedforward step from L2/3 to L5 ^96^. We also found that the MOp/MOs L4 neurons had intracortical long-range projections like the L2 and L3 neurons (**Fig. 6d**).

Taken together, our multimodal characterization demonstrates that mouse MOp indeed has excitatory neurons with L4 characteristics, namely, occupying a specific layer between L2/3 and L5, having simple and untufted dendrites and upward-projecting local axons, belonging to a transcriptomic type (L4/5 IT_1) marked by a L4-specific gene *Rspo1* as well as the intersection of a L2/3/4-specific gene *Cux2* and a L4/5-specific gene *Rorb*, and having corresponding epigenomic regulatory elements. L4-like neurons may also exist in human and marmoset M1.

### Integrated multimodal characterization of two L5 ET projection neuron types in Mop

Previous studies had shown that in the mouse anterolateral motor (ALM) cortex, part of MOs, L5 ET neurons have two transcriptomically distinct projection types that may carry out different motor-control functions; the thalamus projecting type may be involved in movement planning whereas the medulla (MY) projecting type may be involved in initiation of the movement ^15,86^. Here as the second case study, through integrated multimodal characterization we demonstrate that L5 ET neurons in MOp can also be divided into MY-projecting and non-MY-projecting types.

As shown in the companion paper ^45^, we mapped the mouse MOp L5 ET transcriptomic types to the previous VISp-ALM transcriptomic taxonomy ^15^. From this mapping we found that the MOp L5 ET 1 type corresponded to the ALM MY-projecting type marked by *Slco2a1*, whereas MOp L5 ET 2-4 types corresponded to the ALM thalamus-projecting types with L5 ET 2/3 marked by *Hpgd* and L5 ET 4 by *Npsr1*. Here we show such distinction is consistent across all molecular datasets (**Fig. 9a-b**). Mouse transcriptomic type L5 ET 1 corresponded well with both integrated molecular type SCF L5 ET (1) and MERFISH clusters L5_ET_5, as well as with a L5 ET transcriptomic type from human and marmoset. Mouse transcriptomic types L5 ET 2-4 corresponded with integrated molecular types SCF L5 ET (2-3), MERFISH clusters L5_ET_1-4, and two L5 ET transcriptomic types from human and marmoset. The laminar distribution of these two groups was revealed by MERFISH, with cells in L5_ET_1-4 clusters intermingled in the upper part of L5 and cells in L5_ET_5 located distinctly in lower L5 (**Fig. 9c**). The two groups were further distinguished by epigenomic peaks associated with specific marker genes, *Slco2a1* for SCF L5 ET (1) type and *Npnt* for SCF L5 ET (2-3) types (**Fig. 9d**), providing validity to the two novel transgenic driver lines we generated, *Slco2a1-P2A-Cre* and *Npnt-P2A-FlpO* (**Fig. 7**).

Epi-Retro-Seq study (see above) revealed more complex long-range projection patterns among the 6 epigenetic L5-ET clusters identified, with cluster 0 predominantly projecting to MY while other clusters having variable and less specific projection patterns (clusters 2 and 3 also containing MY-projecting cells) (**Fig. 5g**). We co-clustered the L5 ET cells from the Epi-Retro-Seq data and the snRNA-seq 10x v3 B data ^45^ to investigate the correspondence of Epi-Retro-Seq clusters and projection targets with transcriptomic clusters (**Fig. 9e**). We found that the consensus transcriptomic cluster L5 ET 1 corresponds to Epi-Retro-Seq clusters 0, 2 and 3, all of which contain MY-projecting neurons. On the other hand, transcriptomic clusters L5 ET 2-4 correspond to Epi-Retro-Seq clusters 1, 4 and 5, which do not contain MY-projecting neurons. Thus, all MY-projecting neurons are mapped to transcriptomic type L5 ET 1, while neurons in the L5 ET 2-4 types do not project to MY.

Anterograde tracing in *Slco2a1-P2A-Cre* mice demonstrated predominant projection from *Slco2a1*-labeled neurons in MOp to MY (**Fig. 7**). We identified multiple full morphology reconstructions of MOp L5 ET neurons from fMOST imaging of *Fezf2-CreER;Ai166* and *Pvalb-T2A-CreERT2;Ai166* transgenic mice ^68^. These reconstructions could be clearly separated into a MY-projecting group and a non-MY-projecting group (**Fig. 9g**), though they were not directly linked to transcriptomic types yet. Both groups of cells had thick-tufted dendrites that were similar to each other (**Fig. 9g**). Consistent with this, Patch-seq cells corresponding to transcriptomic types L5 ET 1-4 also were indistinguishable from each other by their dendritic morphologies (**Fig. 9f**).

Altogether, our integrated multimodal characterization identified two major types of mouse L5 ET projection neurons, MY-projecting and non-MY-projecting, with distinct gene markers, epigenomic elements, laminar distribution, and corresponding types in human and marmoset.

### An integrated synthesis of multimodal features of cell types in the primary motor cortex

As the conclusion of this series of studies from BICCN, we present an overview and integrated synthesis of the knowledge gained in constructing a multimodal census and atlas of cell types in the primary motor cortex of mouse, marmoset and human (**Fig. 10**). A critical aspect of our studies is that this synthesis is only made possible by the systematic integrative computational analyses across multiple transcriptomic and epigenomic data types that connect a diverse range of cellular features together at cell subclass or type level to allow mutual correlation.

**Figure 10.**
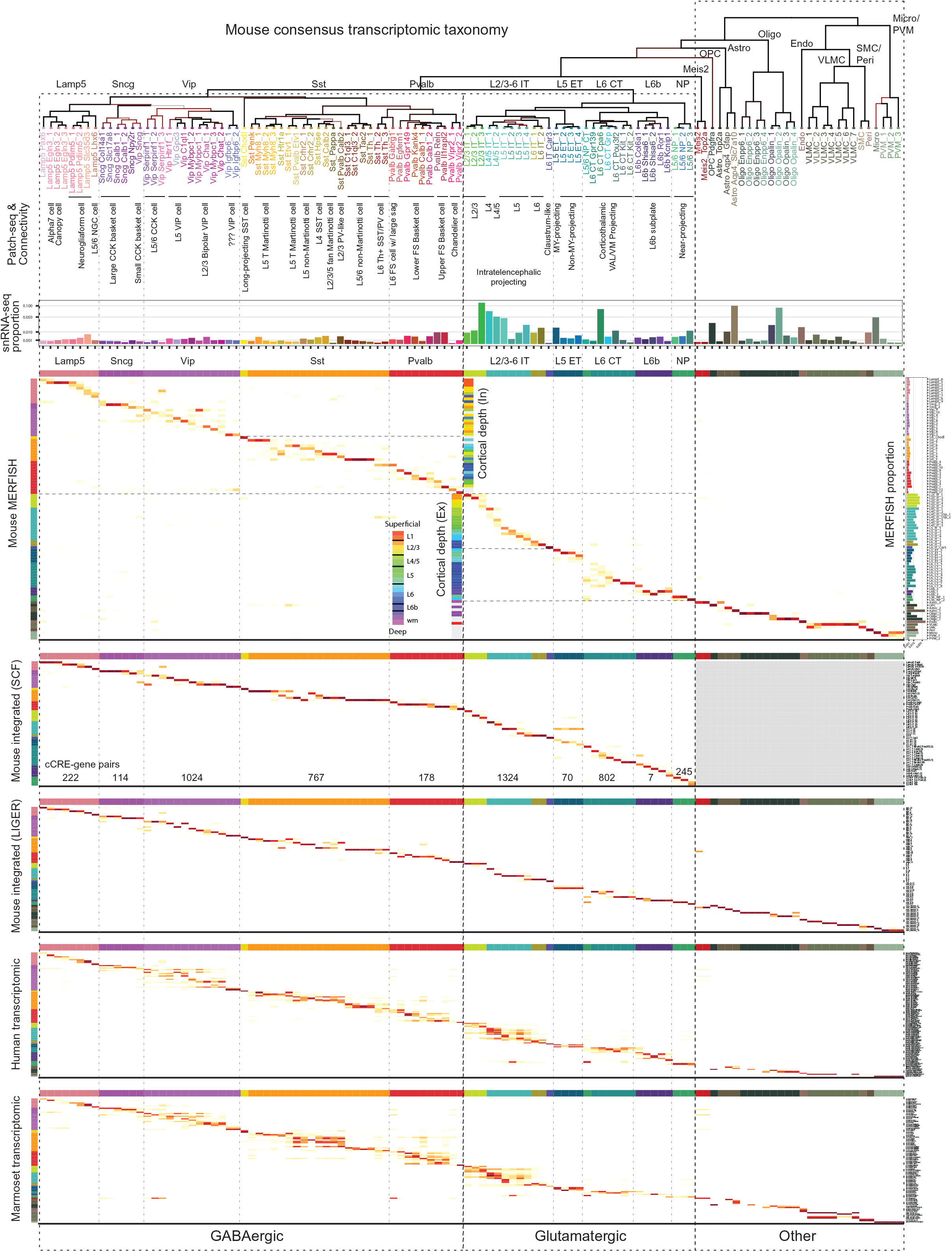
An integrated multimodal census and atlas of cell types in the primary motor cortex of mouse, marmoset and human. The mouse MOp consensus transcriptomic taxonomy at the top is used to anchor cell type features in all the other modalities. Subclass labels are shown above major branches and cluster labels are shown below each leaf node. Confusion matrices show the correspondence between the mouse MOp transcriptomic taxonomy with those derived from other molecular datasets, including mouse MERFISH, the integrated mouse molecular taxonomies by SingleCellFusion (SCF) or LIGER, and the human and marmoset transcriptomic taxonomies. Using Patch-seq and connectivity studies, many transcriptomic neuronal types or subclasses are annotated and correlated with known cortical neuron types traditionally defined by electrophysiological, morphological and connectional properties. (Note: no Patch-seq data were collected for the Vip cells labeled by question marks.) The relative proportions of all cell types within the mouse MOp are calculated from either the snRNA-seq 10x v3 B (horizontal bar graph) or MERFISH (vertical bar graph to the right of the MERFISH matrix) dataset. Median cortical depth position of each cell type derived from MERFISH is shown as color-coded bar graphs at the center of the MERFISH matrix, colored according to the rainbow scheme from superficial (red) to deep (purple) layers as shown on the left. Cell types with dispersed distributions spanning relatively large ranges of cortical depth are colored in grey. The numbers of cCRE-gene pairs in modules corresponding to neuronal subclasses identified by Cicero from the scRNA-seq and snATAC-seq datasets are shown at the bottom of the SCF matrix.

This integrated synthesis uses the mouse MOp consensus transcriptomic taxonomy (containing 18 subclasses and 116 clusters/types) ^45^ as the anchor (**Fig. 10**) because it was derived from the largest and deepest datasets and was the reference taxonomy for nearly all the cross-modality and cross-species comparisons. Correspondence matrices between the different molecular modalities show that the mouse MERFISH-based spatial transcriptomic taxonomy (95 clusters) ^54^, the integrated mouse molecular taxonomies combining transcriptomic and epigenomic data using either SingleCellFusion (SCF, 56 neuronal clusters) or LIGER (71 clusters) approach ^45^, and the human and marmoset transcriptomic taxonomies (127 and 94 clusters, respectively) ^48^ all aligned largely consistently with the mouse consensus transcriptomic taxonomy. Such alignment convincingly demonstrates that cell types in a given brain region can be consistently described by different types of characterization. At the same time, it should also be noted that the alignments are not perfect and disagreements do exist at the individual cluster level (which is most pronounced in cross-species comparisons), suggesting that differential variations exist in different data types and consistency, in particular that across species, may be more appropriately described at an intermediate level of granularity.

In this integrated synthesis, we can further assign additional attributes to the molecularly defined cell types (**Fig. 10**). Based on Patch-seq ^64^, Retro-seq (e.g. Epi-Retro-Seq ^79^), Retro-MERFISH ^54^, and axon projection ^68,69^ studies, we relate many transcriptomic neuronal types or subclasses to cortical neuron types traditionally defined by electrophysiological, morphological and connectional properties, thus bridging our cell type census with historical and community knowledge. We provide the relative proportion of each cell type within the mouse MOp using either snRNA-seq or MERFISH data. The MERFISH data also identify the spatial distribution pattern of each cell type ^54^. For example, we found that excitatory or inhibitory neuron types are distributed along the cortical depth, with many individual types adopting narrow cortical-depth distributions, often occupying predominantly a single layer or a sublayer, and related types (e.g. the L2/3-6 IT excitatory types) can display a gradual transitioning across cortical depths/layers. On the other hand, non-neuronal cell types are either distributed across all layers or specific to layer 1 or the white matter (WM). Patch-seq data also provided the cortical depth positions of a variety of neuronal cell types.

Finally, we demonstrate the possibility to elucidate gene regulatory mechanisms by discovering candidate cis-regulatory elements (cCREs) as well as master transcription factors (TFs) specific to neuronal subclasses by mining the combined mouse MOp transcriptomic and epigenomic datasets to access both RNA expression of genes and accessibility or DNA methylation of cCREs from the same cell clusters.

For example, we found 7,245 distal cCRE (>1 kbp from transcriptional start site)-gene pairs in neuronal cells in MOp that showed positive correlation between accessibility at the 6,280 cCREs and expression levels of 2,490 putative target genes (see Methods, and companion papers ^45,52^). We grouped these putative enhancers into modules based on accessibility across cell clusters (**Extended Data Fig. 2**). 76% of putative enhancers showed remarkable sub-type specific chromatin accessibility and were enriched for lineage-specific transcription factors, while 24% of putative enhancers (1,527) were widely accessible and linked to genes expressed across neuronal cell clusters with highest expression levels in Glutamatergic neurons (module M1, **Extended Data Fig. 2b**). Putative enhancers in this module showed enrichment of sequence motifs recognized by transcription factors CTCF, MEF2 indicating a more general rule of these factors in establishing neuronal gene regulatory programs (**Extended Data Fig. 2c**). Meanwhile, other modules (M2 to M14) of enhancer-gene pairs were active in a subclass-specific manner (**Extended Data Fig. 2b-d**). Thus, using this approach we have identified a large number of enhancer-gene pairs for each subclass of neurons (**Fig. 10**). These enhancers can be potentially used to generate cell type-targeting viral tools.

Similarly, we identified transcription factors showing cell-type specificity supported by both RNA expression and DNA binding motif enrichment in hypo-CG-DMR of MOp subclasses (see Methods, and companion papers ^45,50^) (**Extended Data Fig. 3**). Combining these two orthologous pieces of evidence identified many well-studied TFs in embryonic precursors, such as the Dlx family members for pan-inhibitory neurons, and Lhx6 and Mafb for MGE derived inhibitory neurons. We further identified many additional TFs with more restricted patterns in specific subclasses, such as Rfx3 and Rreb1 (in L2/3 IT), Atoh7 and Rorb (in L4/5 IT), Pou3 family members (in L5 ET), Etv1 (in L5/6 NP), Esrr family members (in Pvalb), and Arid5a (in Lamp5). The agreement of these two modalities suggests a requirement of TF regulatory activity in mature neurons to maintain aspects of cell phenotypes and identity.

In summary, our comprehensive multimodal characterization of cell types from the MOp region demonstrates that transcriptomic, epigenomic, spatial, physiological, morphological and connectional properties can be all correlated and integrated together, to reveal organizational principles of brain cell types and bridge molecular, structural and functional studies in different modalities and across species.

## DISCUSSION

### A cell census and atlas of motor cortex

Understanding the principles of brain circuit organization requires a detailed understanding of its basic components, but the cellular diversity and complexity of the brain, including the neocortex, have defied a comprehensive and quantitative description. Single-cell transcriptomics and epigenomics, as well as spatially resolved single-cell transcriptomics, are accelerating efforts to classify all molecular cell types in many organ systems ^40,99^, including the brain ^5,6,100^. The current effort combines these technologies to derive a robust and comprehensive molecular cell type classification and census of the primary motor cortex of mouse, marmoset and human, coupled with a spatial atlas of cell types and an anatomical input/output wiring diagram in mouse. We demonstrate the robustness and validity of this classification through strong correlations across cellular phenotypes, and strong conservation across species. Together these data comprise a cell atlas of primary motor cortex that encompasses a comprehensive reference catalog of cell types, their proportions, spatial distributions and anatomical and physiological characteristics, and molecular genetic profiles, registered into a Common Coordinate Framework^41^. This cell atlas establishes a foundation for an integrative study of the architecture and function of cortical circuits akin to reference genomes for studying gene function and genome regulatory architecture. Furthermore, it provides a comprehensive map of the genes that contribute to cellular phenotypes and their epigenetic regulation. These data resources and associated tools enabling genetic access for manipulative experimentation are publicly available and provide a roadmap for exploring cellular diversity and organization across brain regions, organ systems, and species.

The molecular classification presented here is overall consistent with prior literature and synthesizes a wide body of existing and new information into a coherent quantitative framework that provides metrics for the robustness of, and the similarities and distinctions between, cell types. For motor cortex, as for other cortical regions ^15,18^, this cellular organization is hierarchical, with different branches comprising major cell classes, subclasses, and types representing the finest resolution clusters afforded by each method. This classification provides strong evidence for the existence of hitherto poorly studied but molecularly distinct subclasses such as the near-projecting (NP) pyramidal neurons, and many more novel cell types. At the level of cell class and subclass (and some highly distinctive types like chandelier cells and long-range projecting Sst Chodl interneurons), we find remarkable concordance across transcriptomics, epigenomics, spatial patterning, physiology and connectivity, as well as strong homology across species. The class and subclass branches clearly represent different developmental programs, such as GABAergic neuron derivatives of different zones of the ganglionic eminences ^101,102^ or the layer-selective glutamatergic neurons derived sequentially from progenitors of the cortical plate, and the hierarchical organization generates new hypotheses about developmental origins of highly distinctive cell types. This quantitative hierarchy also challenges well-established nomenclature systems. For example, the term glia is typically used to encapsulate astrocytes, oligodendrocytes and OPCs, and microglia. However, microglia are not closely related to these neuroectoderm-derived populations based on transcriptomics or developmental origins ^103^ and should be grouped with other more similar non-neuronal cell types such as endothelial cells, VLMCs and pericytes. Substantial challenges remain for redefining data-driven cell ontologies and nomenclature systems ^100,104^.

Comparisons of the MOp results described here to other regions also help to understand what makes the motor cortex functionally distinct. Previous transcriptomic studies suggested that GABAergic interneuron types are shared among cortical regions whereas glutamatergic projection neuron types exhibit gradient-like distribution across the cortical sheet and are more distinct between distant regions but more similar between neighboring regions ^15,44^. Thus the projection neurons in MOp are more similar to those of nearby regions, yet our anatomical tracing study defines a MOp-specific input-output wiring diagram. This result suggests that differential axonal projections of similar molecular types among different cortical areas may be the major feature defining regional functional specificity. We also find substantial variation in the proportion of specific cell types between cortical areas. For example, we identify two glutamatergic neuron types that distinguish MOp from its neighboring primary somatosensory (SSp) region, the L4 IT neurons that are present in MOp at lower abundance level than in SSp and the *Slco2a1*-expressing, medulla-projecting L5 ET neurons that are more abundant in MOp than in SSp ^54,68^. These regional differences in cellular makeup may contribute to the functional specialization of MOp as well.

#### Cell type discreteness, variation and phenotypic concordance

The concordance of transcriptomic and epigenomic results and their overall correlation with other cellular phenotypes, including spatial distributions, morphological properties, electrophysiological properties, and projection/connectivity, strongly argues for a unifying molecular genetic framework for understanding cortical cell types, particularly at the level of subclasses and distinctive cell types. At the same time, substantial multimodal variations at finer granularity appear to preclude a fully discretized representation of cell types with consistency across all cellular phenotypes. One source of variation is differences in granularity with different molecular data modalities, with transcriptomics providing the highest granularity at present. This may reflect true biology or differences in technological information content, for example sparse genome coverage in epigenetic methods. A second source involves continuous rather than discrete variation. For example, while some highly granular cell types are highly distinct from others (e.g. L6 IT Car3, Sst Chodl and Pvalb chandelier cells), many other types exhibit continuous variation in their properties both within types and among closely related types with no clear boundaries between them. However, even at this fine-grained level of continuous variation, spatial, morphological and physiological properties often co-vary with transcriptomic profiles as shown by MERFISH and Patch-seq. Similar findings on continuous as well as unitary variations have been reported for hippocampal interneurons ^16^. These results suggest that continuous phenotypic variation may represent a general organizing principle underlying the diversification of brain cell types.

As shown in our mouse Epi-Retro-Seq, MERFISH, and single-neuron full morphology and projection studies there is a strong correlation between molecular phenotype and axonal target specificity at the subclass level (e.g., IT, L5 ET, L6 CT). This was also the case for medulla-projecting L5 ET type. However, a strict correlation between molecular cell types and specific axonal projection targets was not generally observed. It is possible that axon pathfinding during development involves stochastic decisions and subsequent activity-dependent pruning that mature cell transcriptomes do not represent. Furthermore, individual projection neurons typically have collaterals to many different target regions which complicates understanding these relationships. Comprehensive datasets on the complete axonal projections of individual neurons whose molecular identity is clearly established will be needed to address this issue.

#### Cell type conservation and divergence

Evolutionary conservation is strong evidence of functional significance. The demonstrated conservation of cell types from mouse, marmoset, macaque and human strongly suggests that these conserved types play important roles in cortical circuitry and contribute to a common blueprint essential for cortical function in mammals and even more distantly related species. We also find that similarity of cell types varies as a function of evolutionary distance, with substantial species differences that either represent adaptive specializations or genetic drift. For the most part species specializations tend to appear at the finer branches or leaves of the hierarchical taxonomy. This result is consistent with a recent hypothesis in which cell types are defined by common evolutionary descent and evolve independently, such that new cell types are generally derived from existing genetic programs and appear as specializations at the finer levels of the taxonomic tree ^105^.

A surprising finding across all homologous cell types was the relatively high degree of divergence for genes with highly cell type-specific expression in a given species. This observation provides a clear path to identify the core conserved genes underlying the canonical identity and features of those cell types. Furthermore, it highlights the need to understand species adaptations superimposed on the conserved program, as many specific cellular phenotypes may vary across species including gene expression, epigenetic regulation, morphology and connectivity, and physiological functional properties. As we illustrate in the Betz cells, there is clear homology across species in the layer 5 ET subclass, but variation in many measurable properties across species.

#### A framework for linking model organisms to human biology and disease

The results presented have major utility and implications for the consideration of model organisms to understand human brain function and disease. Despite major investments, animal models of neuropsychiatric disorders have often been characterized by “loss of translation,” fueling heated debates about the utility of model organisms in the search for therapeutic targets for treating human diseases. The molecular genetic framework of cell type organization established by the current study will provide a robust cellular metric system for cross-species translation of knowledge and insight that bridges levels of organization based on their inherent biological and evolutionary relationships. For example, the characterization of cell types and their properties shown in **Figure 10** can be used to infer the main characteristics of homologous cell types in humans and other mammalian species, despite the often extreme difficulty in measuring their specific properties in those species. On the other hand, they also reveal the potential limitations of model organisms and the necessity to study human and closely related primate species to understand the specific features of cell types as they contribute to human brain function and susceptibility to human-specific diseases. Having cell census information aligned across species as illustrated here should be highly valuable for making rational choices about the best models for each disease and therapeutic target. This reductionist dissection of the cellular components provides a foundation for understanding the general principles of neural circuit organization and computation that underlie mental activities and brain disorders.

#### Future directions

The success of the current strategy to systematically and comprehensively dissect cell types and generate a cell census and atlas opens up numerous avenues for future work. This census and atlas form the foundation for the larger community to study specific features of cell types and aggregate information about cell types across species much as genomic databases aggregate information about genes. Classification of cell types and description of their molecular, spatial, and connectional properties in the adult sets the stage for developmental studies to understand the molecular genetic programs underlying cell type specification, maturation and circuit connectivity. The molecular classification and the utility of combined single cell transcriptomics and epigenomics to identify functional enhancers promises to deliver tools for genetic access to the great majority of brain cell types via transgenic and viral strategies. A combination of some of the approaches, such as imaging-based single-cell transcriptomics, with behavior stimulation and functional imaging can further elucidate the functional roles of distinct cell types in circuit computation. This systematic, multi-modal strategy described here is extensible to the whole brain, and major efforts are underway in the BICCN to generate a brain-wide cell census and atlas in the mouse with increasing coverage of human and non-human primates.

## METHODS

### Integrating 10x v3 snRNA-seq datasets across species

To identify homologous cell types across species, human, marmoset, and mouse 10x v3 snRNA-seq datasets were integrated using Seurat’s SCTransform workflow. Each major cell class (glutamatergic, GABAergic, and non-neuronal cells) was integrated separately across species. Expression matrices were reduced to 14,870 one-to-one orthologs across the three species (NCBI Homologene, 11/22/2019). Nuclei were downsampled to have approximately equivalent numbers at the subclass level across species. Marker genes were identified for each species’ cluster and used as input to guide alignment and anchor-finding during integration steps. For full methods see ^48^.

### Estimation of cell type homology

To establish a robust cross-species cell type taxonomy, we applied a tree-based clustering method on integrated class-level datasets (https://github.com/AllenInstitute/BICCN_M1_Evo). The integrated space (from the above mentioned Seurat integration) was over-clustering into small sets of highly similar nuclei for each class (∼500 clusters per class). Clusters were aggregated into metacells, then hierarchical clustering was performed based on the metacell gene expression matrix using Ward’s method. Hierarchical trees were then assessed for cluster size, species mixing, and branch stability by subsampling the dataset 100 times with 95% of nuclei. Finally, we recursively searched every node of the tree, and if certain heuristic criteria were not sufficed for a node below the upper node, all nodes below the upper node were pruned and nuclei belonging to this subtree were merged into one homologous group. We identified 24 GABAergic, 13 glutamatergic, and 8 non-neuronal cross-species consensus clusters that were highly mixed across species and robust. For full methods see ^48^.

### Cross-species differential gene expression and correlations

Expression matrices for each species, for each major cell class (GABAergic, glutamatergic, and non-neuronal cells) were normalized using Seurat’s SCTransform function with default parameters to generate a ‘corrected UMI’ matrix and remove technical variation within each species. SCTransform normalized counts matrices were then counts per 100,000 UMI (CP100K) normalized to account for variable sequencing depths between species. CP100K normalization was performed by multiplying each value in the ‘corrected UMI’ (SCTransform normalized) matrix by 100,000 and dividing by the column sums (total UMIs from each nuclei). SCTransform-CP100K normalized matrices were then used to find DE genes and correlations between species for each cross-species cluster.

DE gene analysis was performed with Seurat’s FindAllMarkers function, using the Wilcoxon rank sum test, between each pair of species for a given cross-species cluster (e.g. human Lamp5_1 vs. marmoset Lamp5_1, human Lamp5_1 vs. mouse Lamp5_1, and marmoset Lamp5_1 vs. mouse Lamp5_1). Marker genes (FDR < 0.01, log fold-change > 2, expressed in at least 10% of nuclei) from each pairwise species comparison were identified for each cross-species cluster. We report the sum of marker genes between each species comparison as a heatmap in Figure 2e and show that human and marmoset have fewer DE genes between each other than with mouse across all cross-species clusters.

To visualize the correspondence of a given cross-species cluster between each pair of species, we first found the average SCTransform-CP100K expression for each cross-species cluster for each species. Average expression was then log-transformed and the spearman correlations between each species pair were identified and reported in the Figure 2d heatmap, which shows human and marmoset have higher correlations than either primate with mouse for all clusters except Endo, VLMC, and Microglia/PVM clusters (likely due to differences in sampling).

### Integrating mouse transcriptomic, spatially resolved transcriptomic, and epigenomic datasets

To integrate IT cell types from different mouse datasets, we first take all cells that are labeled as IT, except for L6_IT_Car3, from the 11 datasets as listed in Figure 8a. These cell labels are either from dataset-specific analyses ^54,64,79^, or from the integrated clustering of multiple datasets^45^. The integrated clustering and embedding of the 11 datasets are then generated by projecting all datasets into the 10x v2 scRNA-seq dataset using SingleCellFusion ^45,59^. Genome browser views of IT and ET cell types (Figure 8b and Figure 9d) are taken from the corresponding cell types of the brainome portal (brainome.ucsd.edu/BICCN_MOp) ^45^.

### Integration of L5 ET cells from Epi-Retro-Seq and 10x snRNA-Seq

For snRNA-Seq, the 4,515 cells from 10x v3 B dataset labeled as L5 ET by SCF were selected^45^. The read counts were normalized by the total read counts per cell and log transformed. Top 5,000 highly variable genes were identified with Scanpy ^106^ and z-score scaled across all the cells. For Epi-Retro-Seq, the posterior methylation levels of 12,261 genes in the 848 L5 ET cells were computed ^79^. Top 5,000 highly variable genes were identified with AllCools ^59^ and z-score scaled across all the cells. The 1,512 genes as the intersection between the two highly variable gene lists were used in Scanorama ^107^ to integrate the z-scored expression matrix and minus z-scored methylation matrix with sigma equal to 100.

### Identification of candidate cis-regulatory elements

For peak calling in the snATAC-seq data, we extracted all the fragments for each cluster, and then performed peak calling on each aggregate profile using MACS2 ^108^ with parameter: “-- nomodel --shift −100 --ext 200 --qval 1e-2 –B --SPMR”. First, we extended peak summits by 250 bp on either side to a final width of 501 bp. Then, to account for differences in performance of MACS2 based on read depth and/or number of nuclei in individual clusters, we converted MACS2 peak scores (-log10(q-value)) to “score per million” ^109^. Next, a union peak set was obtained by applying an iterative overlap peak merging procedure, which avoids daisy-chaining and still allows for use of fixed-width peaks. Finally, we filtered peaks by choosing a “score per million” cut-off of 5 as candidate cis-regulatory elements (cCREs) for downstream analysis.

### Predicting enhancer-promoter interactions

First, co-accessible cCREs are identified for all open regions in all neurons types (cell clusters with less than 100 nuclei from snATAC-seq are excluded), using Cicero ^110^ with following parameters: aggregation k = 50, window size = 500 kb, distance constraint = 250 kb. In order to find an optimal co-accessibility threshold, we generated a random shuffled cCRE-by-cell matrix as background and calculated co-accessible scores from this shuffled matrix. We fitted the distribution of co-accessibility scores from random shuffled background into a normal distribution model by using R package fitdistrplus ^111^. Next, we tested every co-accessible cCRE pair and set the cut-off at co-accessibility score with an empirically defined significance threshold of FDR<0.01. The cCREs outside of ± 1 kb of transcriptional start sites (TSS) in GENCODE mm10 (v16) were considered distal. Next, we assigned co-accessibility pairs to three groups: proximal-to-proximal, distal-to-distal, and distal-to-proximal. In this study, we focus only on distal-to-proximal pairs. We calculated the Pearson’s correlation coefficient (PCC) between gene expression (scRNA SMART-seq) and cCRE accessibility across the joint clusters to examine the relationships between the distal cCREs and target genes as predicted by the co-accessibility pairs. To do so, we first aggregated all nuclei/cells from scRNA-seq and snATAC-seq for every joint cluster to calculate accessibility scores (log2 CPM) and relative expression levels (log2 TPM). Then, PCC was calculated for every gene-cCRE pair within a 1 Mbp window centered on the TSS for every gene. We also generated a set of background pairs by randomly selecting regions from different chromosomes and shuffling of cluster labels. Finally, we fit a normal distribution model and defined a cut-off at PCC score with an empirically defined significance threshold of FDR<0.01, in order to select significant positively correlated cCRE-gene pairs.

### Identification of cis-regulatory modules

We used Nonnegative Matrix Factorization (NMF) to group cCREs into cis-regulatory modules based on their relative accessibility across cell clusters. We adapted NMF (Python package: sklearn) to decompose the cell-by-cCRE matrix V (N×M, N rows: cCRE, M columns: cell clusters) into a coefficient matrix H (R×M, R rows: number of modules) and a basis matrix W (N×R), with a given rank R:

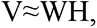

The basis matrix defines module related accessible cCREs, and the coefficient matrix defines the cell cluster components and their weights in each module. The key issue to decompose the occupancy profile matrix was to find a reasonable value for the rank R (i.e., the number of modules). Several criteria have been proposed to decide whether a given rank R decomposes the occupancy profile matrix into meaningful clusters. Here we applied a measurement called “Sparseness” ^112^ to evaluate the clustering result. Median values were calculated from 100 times for NMF runs at each given rank with a random seed, which will ensure the measurements are stable. Next, we used the coefficient matrix to associate modules with distinct cell clusters. In the coefficient matrix, each row represents a module and each column represents a cell cluster. The values in the matrix indicate the weights of clusters in their corresponding module. The coefficient matrix was then scaled by column (cluster) from 0 to 1. Subsequently, we used a coefficient > 0.1 (∼95th percentile of the whole matrix) as a threshold to associate a cluster with a module. Similarly, we associated each module with accessible elements using the basis matrix. For each element and each module, we derived a basis coefficient score, which represents the accessible signal contributed by all clusters in the defined module.

### Identification of subclass-selective TFs by both RNA expression and motif enrichment

All analyses for this section were at the subclass level. For RNA expression, we used the sc SMART-seq dataset and compared each subclass with the rest of the population through a one-tailed Wilcoxon test and FDR correction to select significantly differentially-expressed transcription factors (adjusted P-value < 0.05, cluster average fold change > 2). To perform the motif enrichment analysis, we used known motifs from the JASPAR 2020 database ^113^ and the subclass specific hypo-CG-DMR identified in Yao et al ^45^. The AME software from the MEME suite (v5.1.1) ^114^ was used to identify significant motif enrichment (adjusted P-value < 1e-3, odds ratio > 1.3) using default parameters and the same background region set as described in Yao et al ^45^. All genes in **Extended Data Figure 3** were both significantly expressed and had their motif enriched in at least one of the subclasses.

### Generation and use of transgenic mouse lines

Npnt-P2A-FlpO and Slco2a1-P2A-Cre mouse driver lines were generated by CRISPR/Cas9-mediated homologous recombination (Stafford et al., BICCN companion manuscript in preparation). Details are provided in the Supplementary Methods.

All experimental procedures were approved by the Institutional Animal Care and Use Committees (IACUC) of Cold Spring Harbor Laboratory, University of California, Berkeley and Allen Institute, in accordance with NIH guidelines. Mouse knock-in driver lines are being deposited at the Jackson Laboratory for wide distribution.

### Data and code availability

**Figure 1. Summary of experimental and computational approaches taken as well as community resources generated by the BICCN**

All primary data available through the BICCN portal, data archives, and supporting tools.

- Brain Cell Data Center (BCDC), www.biccn.org
- Neuroscience Multi-Omics Archive (NeMO), www.nemoarchive.org
- Brain Image Library (BIL), www.brainimagelibrary.org
- Neurophysiology (DANDI), <u>dandiarchive.org</u>
- Allen Transcriptomics Explorer, https://portal.brain-map.org/atlases-and-data/rnaseq
- NeMO Analytics, www.nemoanalytics.org
- Morphological reconstructions, NeuroMorpho, www.neuromorpho.org

**Figure 2. MOp consensus cell type taxonomy**

#### Primary Data

|  |  |  |
| --- | --- | --- |
| Panel a |  | http://data.nemoarchive.org/publication_release/MOp_MiniAtlas_2020/ |
| Panels b-g | 10x V3 macaque | http://data.nemoarchive.org/biccn/lab/lein/lein/transcriptomic/sncell/raw/ |
|  | 10x V3 human (10X159-1 through 10x160-8) | http://data.nemoarchive.org/biccn/lab/linnarsson/transcriptome/sncell/10X/raw/10X159-1/ |
|  | 10x V3 marmoset (bi005_m1, bi006_m1) | http://data.nemoarchive.org/biccn/lab/feng/transcriptome/sncell/raw/ |
|  | 10x V3 mouse broad data (files with prefix pBICCNsMMrMOp) | http://data.nemoarchive.org/biccn/grant/huang/macosko/transcriptome/sncell/raw |
| Panel h: |  |  |

#### Intermediate analyses

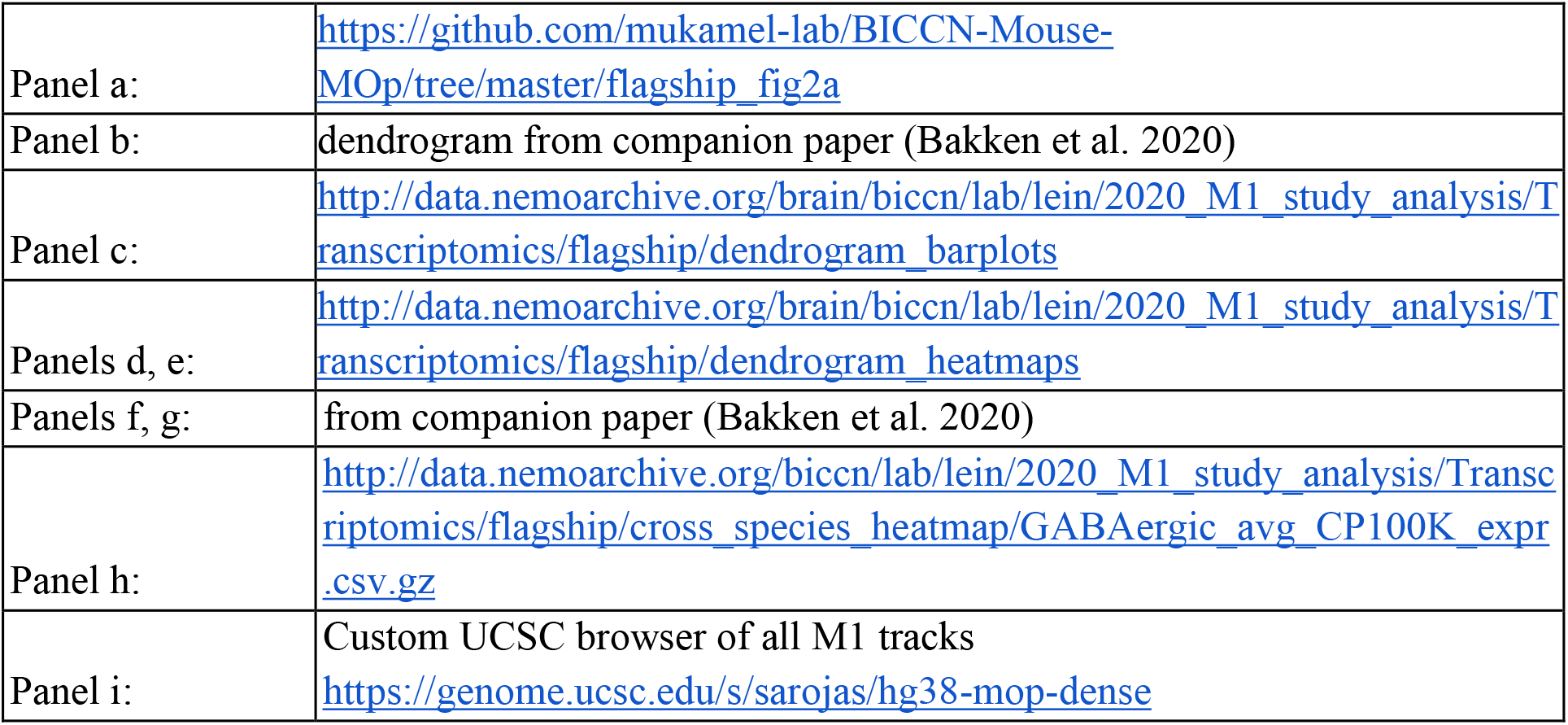

#### Extended Data

|  |  |
| --- | --- |
| Panel j: | Browser https://brainome.ucsd.edu/annoj/BICCN_MOp/ |
| Extended Data Figure 1: | http://data.nemoarchive.org/brain/biccn/lab/lein/2020_M1_study_analysis/Transcriptomics/flagship/cluster_overlap_plots |

**Figure 3: In situ cell-type identification, spatial mapping and projection mapping of individual cells in the MOp by MERFISH**

#### Primary Data

ftp://download.brainimagelibrary.org:8811/02/26/02265ddb0dae51de/

**Figure 4. Correspondence between transcriptomic and morpho-electrical properties of mouse MOp neurons by Patch-seq, and cross-species comparison of L5 ET neurons.**

#### Primary Data

|  |  |  |
| --- | --- | --- |
| Panel b,c,j,k | Electrophysiology data | https://dandiarchive.org/dandiset/000008 |
| Panel h: | 10x V3 macaque | http://data.nemoarchive.org/biccn/lab/lein/lein/transcriptomic/sncell/raw/ |
|  | 10x V3 human (10X159-1 through 10x160-8) | http://data.nemoarchive.org/biccn/lab/linnarsson/transcriptome/sncell/10X/raw/10X159-1/ |
|  | 10x V3 marmoset (bi005_m1, bi006_m1) | http://data.nemoarchive.org/biccn/lab/feng/transcriptome/sncell/raw/ |
|  | 10x V3 mouse broad data (files with prefix pBICCNsMMrMOp) | http://data.nemoarchive.org/biccn/grant/huang/macosko/transcriptome/sncell/raw |

#### Intermediate analyses

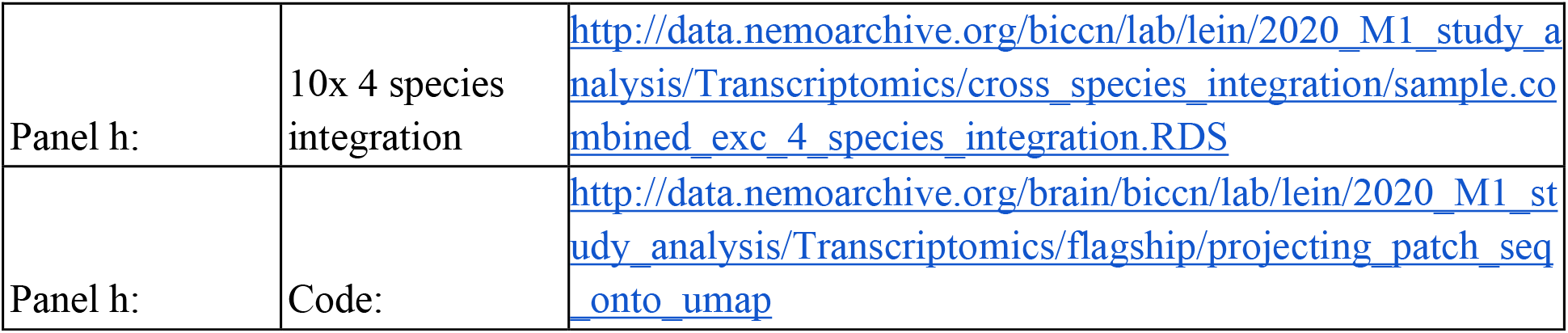

|  |  |  |
| --- | --- | --- |
| Panel h: | 10x 4 species integration | http://data.nemoarchive.org/biccn/lab/lein/2020_M1_study_analysis/Transcriptomics/cross_species_integration/sample.combined_exc_4_species_integration.RDS |
| Panel h: | Code: | http://data.nemoarchive.org/brain/biccn/lab/lein/2020_M1_study_analysis/Transcriptomics/flagship/projecting_patch_seq_onto_umap |

**Figure 5: Epi-Retro-Seq links molecular cell type with distal projection targets**

#### Intermediate analyses

https://github.com/zhoujt1994/BICCN2020Flagship.git

**Figure 6: Global wiring diagram and anatomical characterization of MOp-ul neuron types**

#### Primary Data

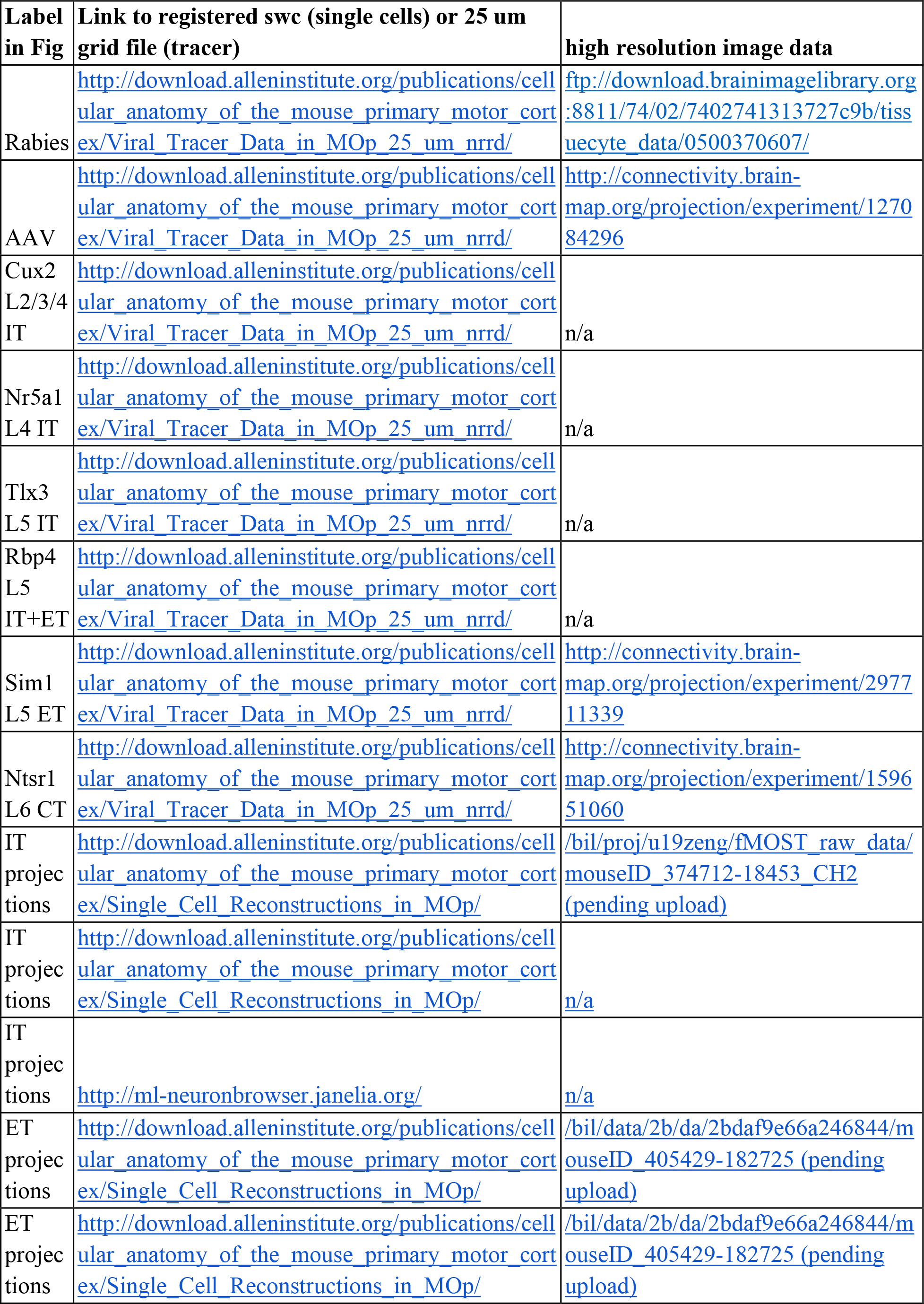
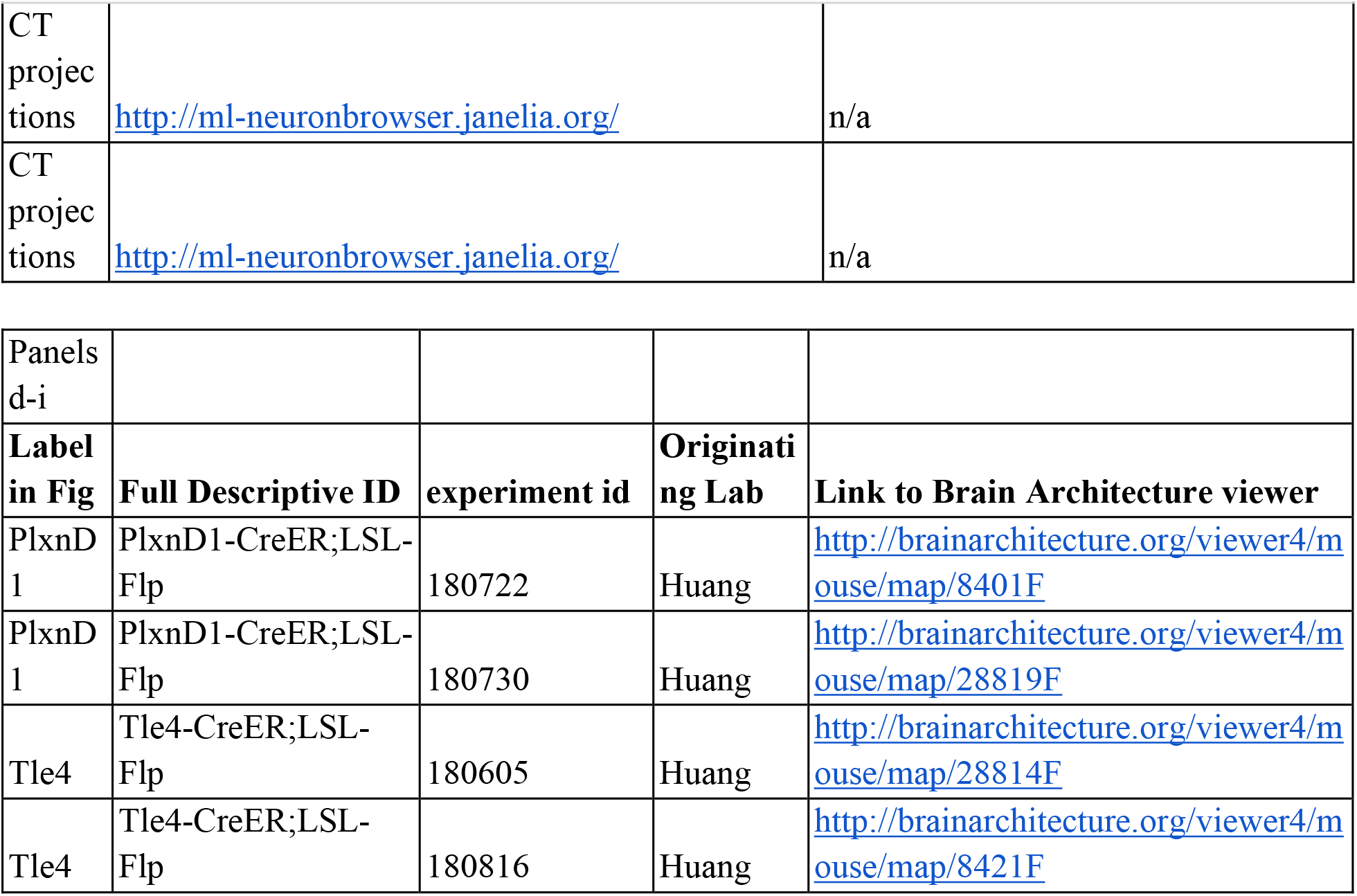

| Label in Fig | Link to registered swc (single cells) or 25 um grid file (tracer) | high resolution image data |
| --- | --- | --- |
| Rabies | http://download.alleninstitute.org/publications/cellular_anatomy_of_the_mouse_primary_motor_cortex/Viral_Tracer_Data_in_MOp_25_um_nrrd/ | ftp://download.brainimagelibrary.org:8811/74/02/7402741313727c9b/tissuecyte_data/0500370607/ |
| AAV | http://download.alleninstitute.org/publications/cellular_anatomy_of_the_mouse_primary_motor_cortex/Viral_Tracer_Data_in_MOp_25_um_nrrd/ | http://connectivity.brain-map.org/projection/experiment/127084296 |
| Cux2<br>L2/3/4<br>IT | http://download.alleninstitute.org/publications/cellular_anatomy_of_the_mouse_primary_motor_cortex/Viral_Tracer_Data_in_MOp_25_um_nrrd/ | n/a |
| Nr5a1<br>L4 IT | http://download.alleninstitute.org/publications/cellular_anatomy_of_the_mouse_primary_motor_cortex/Viral_Tracer_Data_in_MOp_25_um_nrrd/ | n/a |
| Tlx3<br>L5 IT | http://download.alleninstitute.org/publications/cellular_anatomy_of_the_mouse_primary_motor_cortex/Viral_Tracer_Data_in_MOp_25_um_nrrd/ | n/a |
| Rbp4<br>L5<br>IT+ET | http://download.alleninstitute.org/publications/cellular_anatomy_of_the_mouse_primary_motor_cortex/Viral_Tracer_Data_in_MOp_25_um_nrrd/ | n/a |
| Sim1<br>L5 ET | http://download.alleninstitute.org/publications/cellular_anatomy_of_the_mouse_primary_motor_cortex/Viral_Tracer_Data_in_MOp_25_um_nrrd/ | http://connectivity.brain-map.org/projection/experiment/297711339 |
| Ntsr1<br>L6 CT | http://download.alleninstitute.org/publications/cellular_anatomy_of_the_mouse_primary_motor_cortex/Viral_Tracer_Data_in_MOp_25_um_nrrd/ | http://connectivity.brain-map.org/projection/experiment/159651060 |
| IT<br>projections | http://download.alleninstitute.org/publications/cellular_anatomy_of_the_mouse_primary_motor_cortex/Single_Cell_Reconstructions_in_MOp/ | /bil/proj/u19zeng/fMOST_raw_data/mouseID_374712-18453_CH2<br>(pending upload) |
| IT<br>projections | http://download.alleninstitute.org/publications/cellular_anatomy_of_the_mouse_primary_motor_cortex/Single_Cell_Reconstructions_in_MOp/ | n/a |
| IT<br>projections | http://ml-neuronbrowser.janelia.org/ | n/a |
| ET<br>projections | http://download.alleninstitute.org/publications/cellular_anatomy_of_the_mouse_primary_motor_cortex/Single_Cell_Reconstructions_in_MOp/ | /bil/data/2b/da/2bdaf9e66a246844/mouseID_405429-182725 (pending upload) |
| ET<br>projections | http://download.alleninstitute.org/publications/cellular_anatomy_of_the_mouse_primary_motor_cortex/Single_Cell_Reconstructions_in_MOp/ | /bil/data/2b/da/2bdaf9e66a246844/mouseID_405429-182725 (pending upload) |
| CT<br>projec<br>tions | http://ml-neuronbrowser.janelia.org/ | n/a |
| CT<br>projec<br>tions | http://ml-neuronbrowser.janelia.org/ | n/a |

| Panels<br>d-i |  |  |  |  |
| --- | --- | --- | --- | --- |
| Label<br>in Fig | Full Descriptive ID | experiment id | Originati<br>ng Lab | Link to Brain Architecture viewer |
| PlxnD<br>1 | PlxnD1-CreER;LSL-Flp | 180722 | Huang | http://brainarchitecture.org/viewer4/mouse/map/8401F |
| PlxnD<br>1 | PlxnD1-CreER;LSL-Flp | 180730 | Huang | http://brainarchitecture.org/viewer4/mouse/map/28819F |
| Tle4 | Tle4-CreER;LSL-Flp | 180605 | Huang | http://brainarchitecture.org/viewer4/mouse/map/28814F |
| Tle4 | Tle4-CreER;LSL-Flp | 180816 | Huang | http://brainarchitecture.org/viewer4/mouse/map/8421F |

#### Intermediate analyses

|  |  |  |
| --- | --- | --- |
| panel c | https://github.com/AllenInstitute/MOp_anatomy_rendering | code to reproduce rendering of<br>registered data in 3D |

**Figure 7: Genetic tools for targeting cortical glutamatergic projection neuron types**

#### Primary Data

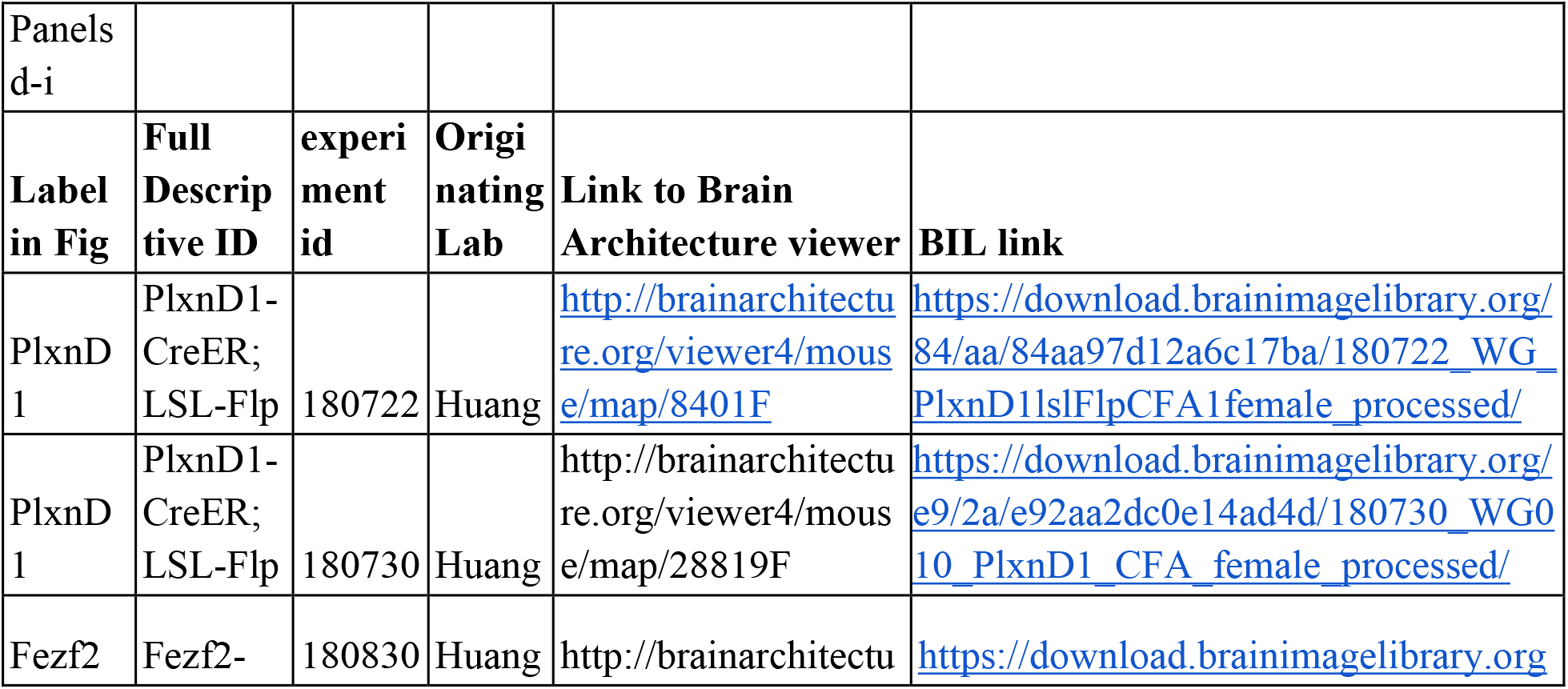
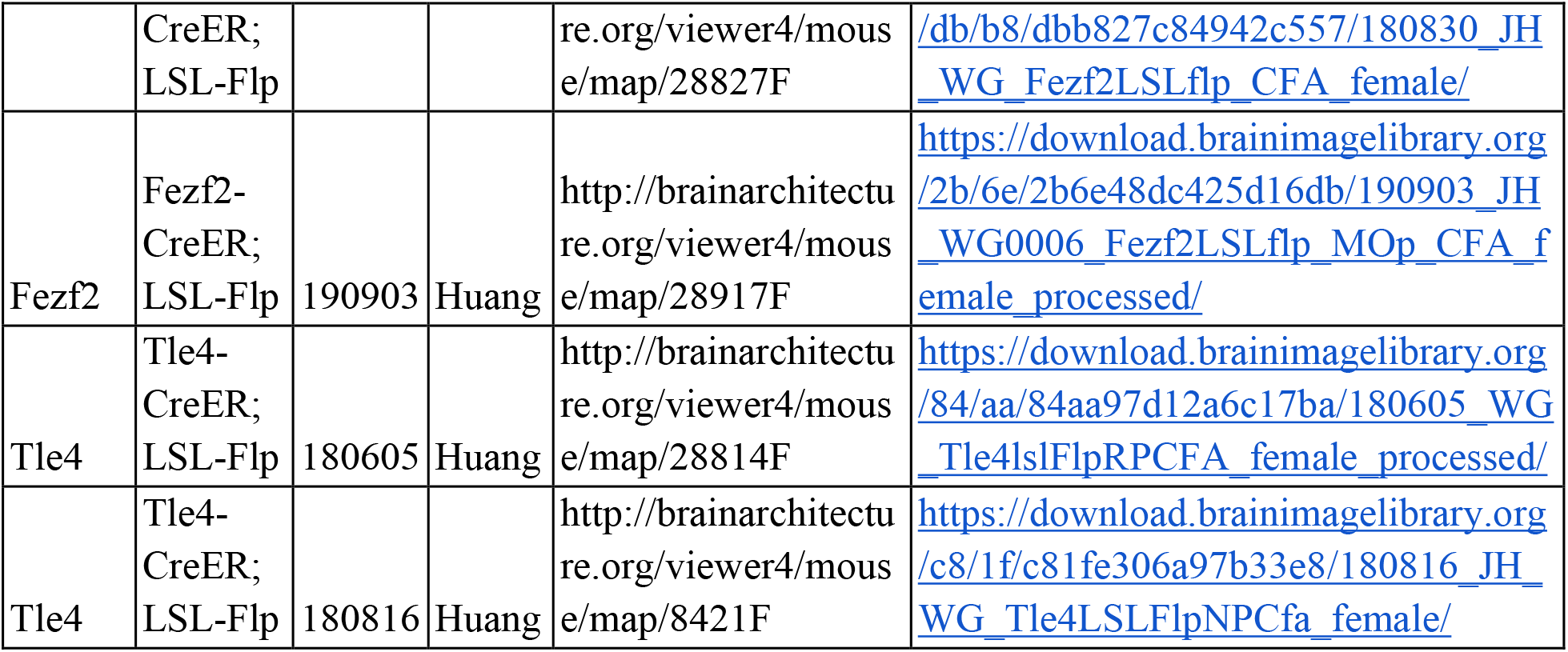

**Figure 8: Existence of L4 excitatory neurons in MOp.**

#### Intermediate analyses

|  |  |
| --- | --- |
| Panels a,c,e | https://github.com/mukamel-lab/BICCN-Mouse-MOp/tree/master/flagship_fig8 |
| Panel b | https://brainome.ucsd.edu/annoj/BICCN_MOp/ |

**Figure 9: Two distinct L5 ET projection neuron types in Mop**

#### Intermediate analyses

|  |  |
| --- | --- |
| Panel d | https://brainome.ucsd.edu/annoj/BICCN_MOp/ |
| Panel e | https://github.com/zhoujt1994/BICCN2020Flagship.git |

**Figure 10: An integrated multimodal census and atlas of cell types in the primary motor cortex of mouse, marmoset and human.**

#### Intermediate analyses

https://github.com/yal054/snATACutils

#### Extended Data

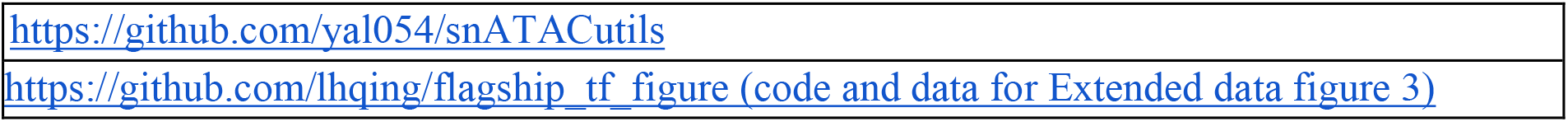

|  |
| --- |
| https://github.com/yal054/snATACutils |
| https://github.com/lhqing/flagship_tf_figure (code and data for Extended data figure 3) |

## SUPPLEMENTARY NOTES

### Nomenclature of the L5 ET subclass of glutamatergic neurons

In this manuscript we have adopted a nomenclature for major subclasses of cortical glutamatergic excitatory neurons, which have long-range projections both within and outside of the cortex, following a long tradition of naming conventions that often classify neurons based on their projection targets. This nomenclature is based on our *de novo* transcriptomic taxonomy (**Fig. 10**) that organizes cell types hierarchically and validates the naming of the primary branches of glutamatergic neurons by their major long-range projection targets. At these levels, glutamatergic neurons are clearly divided into several subclasses, the cortico-cortical and cortico-striatal projecting intratelencephalic (IT) neurons that are distributed across nearly all layers (L2/3 IT, L4/5 IT, L5 IT, L6 IT and L6 IT Car3), the layer 5 neurons projecting to extratelencephalic targets (L5 ET), the cortico-thalamic (CT) projecting neurons in layer 6 (L6 CT), the near-projecting (NP) neurons found in layers 5 and 6, and the L6b neurons whose projection patterns remain largely unknown.

While the IT, CT, NP and L6b neurons have been consistently labeled as such in the field, the L5 ET neurons have not been named consistently in the literature, largely due to their large variety of projection targets and other phenotypic features that vary depending on cortical areas and species. Here we use the term L5 ET (layer 5 extratelencephalic) to refer to this prominent and distinct subclass of neurons as a standard name that can be accurately used across cortical regions and across species, and we provide our rationale below.

It has long been appreciated that cortical layer 5 contains two distinct populations of neurons that can be distinguished, not only based on the presence or absence of projections to ET targets (ET and IT cells), but also based on their predominant soma locations, dendritic morphologies and intrinsic physiology ^81^. Accordingly, various names incorporating these features have been adopted to refer to L5 ET versus L5 IT cells, such as L5b versus L5a, thick-versus thin-tufted and burst-firing versus regular-firing. And the most common term used to refer to L5 ET cells residing in motor cortical areas has been PT, which refers to neurons projecting to the pyramidal tract. As accurately stated in Wikipedia, “The **pyramidal tracts** include both the corticobulbar tract and the corticospinal tract. These are aggregations of efferent nerve fibers from the upper motor neurons that travel from the cerebral cortex and terminate either in the brainstem (*corticobulbar*) or spinal cord (*corticospinal*) and are involved in the control of motor functions of the body.”

Due to the past wide use of the term PT, we do not take the decision to use L5 ET rather than PT lightly. However, in the face of multiple lines of evidence that have accumulated over the last several years ^115,116^ and prominently highlighted in this manuscript, it is now clear that PT represents only a subset of L5 ET cells and is thus unable to accurately encompass the entire L5 ET subclass. This realization is informed by comparisons across species and cortical areas, and by single-cell transcriptomics and descriptions of the projections of single neurons, as well as studies linking transcriptional clusters to projection targets.

As noted above, the overall transcriptomic relationships between cortical neurons are well-described by a hierarchical tree that closely matches developmental lineage relationships as neurons become progressively restricted in their adult fates ^45,48^ (**Fig. 10**). The cortical excitatory neurons are a major branch, distinct from inhibitory, glial, and epithelial cells. Subsequent splitting of the excitatory neurons reveals several major excitatory neuron subclasses – IT, L5 ET, L6 CT, NP and L6b. These major subclasses are conserved across mammalian species ^15,18^, as well as across all cortical areas as shown in mouse ^44^. It is therefore clear that names are needed that both accurately incorporate and accurately distinguish between neurons in these subclasses, and which are applicable across all cortical areas.

Also as noted above, a widely used alternative to L5 ET is PT. Further, this term is traditionally used along with CT to distinguish between cells with these different projections. The two main observations that make these alternative nomenclatures untenable are: 1) PT refers to motor neurons that project into medulla or spinal cord, but in many cortical areas (e.g. visual and auditory areas) none of the L5 ET cells are motor neurons; and 2) even in the motor cortex many cells in the L5 ET subclass do not project to the pyramidal tract and instead project solely to the thalamus (or to thalamus and other non-PT targets). This is revealed by single neuron reconstructions ^26,68,86^ (**Fig. 6 and 9**), BARseq ^67^, projections from neuron populations with known gene expression and anatomical position in mouse lines ^71^, and studies directly linking projections to transcriptomics ^15,54^ and epigenetics ^79^ (**Fig. 5 and 9**). The term PT therefore fails to be inclusive of the entire L5 ET subclass. Furthermore, the L5 CT cells within the L5 ET subclass are largely continuous with PT cells (or “PT-like” cells), not only genetically but also anatomically ^54,64^ (**Fig. 3–4**), as a majority of L5 ET cells project to multiple targets, typically including both the thalamus and the PT structures (e.g., medulla and spinal cord), as well as the midbrain (**Fig. 6 and 9**) ^68^. Thus, the L5 ET subclass should neither be split into PT and CT, nor should the CT-only cells be omitted by use of the term PT. These facts also inform us that it is important to maintain a distinction between L5 CT (a type of L5 ET) and L6 CT (a major subclass of cortical excitatory neurons that is highly distinct from L5 ET, despite the presence of some L6 CT cells at the bottom of layer 5) ^54^. CT can be accurately used as a generic term, but CT neurons do not belong to a single subclass of cortical excitatory neurons.

We recognize that another name that has been used to describe L5 ET cells is SCPN (subcerebral projection neuron) ^82^. Given that the telencephalon is equivalent to the cerebrum, ET and subcerebral have the same meaning and the term L5-SCPN would be an accurate and equivalent alternative. But the “L5” qualifier is crucial in either case in order to distinguish these cells from the L6 CT subclass. We favor the use of ET because SCPN has not been widely adopted and due to symmetry with the widely used “IT” nomenclature. Alternatively, given their evidence that “unlike pyramidal tract neurons in the motor cortex, these neurons in the auditory cortex do not project to the spinal cord”, Chen et al ^67^ used the term “pyramidal tract-like (PT-l).” We also favor L5 ET over L5 PT-l which clings to an inaccurate and now outdated nomenclature.

## Supplementary Methods

### Generation of Npnt-P2A-FlpO and Slco2a1-P2A-Cre mouse lines

To generate lines bearing in-frame genomic insertions of *P2A-FlpO* or *P2A-Cre*, we engineered double-strand breaks at the stop codons of *Npnt* and *Slco2a1*, respectively, using ribonucleoprotein (RNP) complexes composed of SpCas9-NLS protein and in vitro transcribed sgRNA (Npnt: *GATGATGTGAGCTTGAAAAG* and Slco2a1: *CAGTCTGCAGGAGAATGCCT*). These RNP complexes were nucleofected into 10^6^ v6.5 mouse embryonic stem cells (C57/BL6;129/sv; a gift from R. Jaenisch) along with repair constructs in which *P2A-FlpO* or *P2A-Cre* was flanked with the following sequences homologous to the target site, thereby enabling homology-directed repair.

Npnt-P2A-FlpO: TGGCCCTTGAGCTCTAGTGTTCCCACTTGCCATAGAAATCTGATCTTCGGTTTGGGGGAAGGGTTGCCTTACCATGCTCCATGAGTGAGCACTGGGAAAAGGGGCAGAGGAGGCCTGACCAGTGTATACGTTCTCTCCCTAGGTCATCTTCAAAGGTGAAAAAAGGCGTGGTCACACGGGGGAGATTGGATTGGATGATGTGAGCTTGAAGCGCGGAAGATGTGGAAGCGGAGCTACTAACTTCAGCCTGCTGAAGCAGGCTGGAGACGTGGAGGAGAACCCTGGACCTATGGCTCCTAAGAAGAAGAGGAAGGTGATGAGCCAGTTCGACATCCTGTGCAAGACCCCGCCGAAGGTGCTGGTGCGGCAGTTCGTGGAGAGATTCGAGAGGCCCAGCGGCGAAAAGATCGCCAGCTGTGCCGCCGAGCTGACCTACCTGTGCTGGATGATCACCCACAACGGCACCGCGATCAAGAGGGCCACCTTCATGAGTTATAACACCATCATCAGCAACAGCCTGAGTTTTGACATCGTGAACAAGAGCCTGCAGTTCAAGTACAAGACCCAGAAGGCCACCATCCTGGAGGCCAGCCTGAAGAAGCTGATCCCCGCATGGGAGTTCACGATTATCCCTTACAACGGCCAGAAGCACCAGAGCGACATCACCGACATCGTGTCCAGCCTGCAGCTGCAGTTCGAAAGCAGCGAGGAGGCCGACAAGGGGAATAGCCACAGCAAGAAGATGCTGAAGGCCCTGCTGTCCGAAGGCGAGAGCATCTGGGAGATTACCGAGAAGATCCTGAACAGCTTCGAGTACACCAGCAGATTTACCAAAACGAAGACCCTGTACCAGTTCCTGTTCCTGGCCACATTCATCAACTGCGGCAGGTTCAGCGACATCAAGAACGTGGACCCGAAGAGCTTCAAGCTCGTCCAGAACAAGTATCTGGGCGTGATCATTCAGTGCCTGGTCACGGAGACCAAGACAAGCGTGTCCAGGCACATCTACTTTTTCAGCGCCAGAGGCAGGATCGACCCCCTGGTGTACCTGGACGAGTTCCTGAGGAACAGCGAGCCCGTGCTGAAGAGAGTGAACAGGACCGGCAACAGCAGCAGCAACAAGCAGGAGTACCAGCTGCTGAAGGACAACCTGGTGCGCAGCTACAACAAGGCCCTGAAGAAGAACGCCCCCTACCCCATCTTCGCTATTAAAAACGGCCCTAAGAGCCACATCGGC AGGCACCTGATGACCAGCTTTCTGAGCATGAAGGGCCTGACCGAGCTGACAAACGTGGTGGGCAACTGGAGCGACAAGAGGGCCTCCGCCGTGGCCAGGACCACCTACACCCACCAGATCACCGCCATCCCCGACCACTACTTCGCCCTGGTGTCCAGGTACTACGCCTACGACCCCATCAGTAAGGAGATGATCGCCCTGAAGGACGAGACCAACCCCATCGAGGAGTGGCAGCACATCGAGCAGCTGAAGGGCAGCGCCGAGGGCAGCATCAGATACCCCGCCTGGAACGGCATTATAAGCCAGGAGGTGCTGGACTACCTGAGCAGCTACATCAACAGGCGGATCTGAAAGAGGTCGCTGCTGAGAAGACCCCTGGCAGCTCCCGAGCTAGCAGTGAATTTGTCGCTCTCCCTCATTTCCCAATGCTTGCCCTCTTGTCTCCCTCTTATCAGGCCTAGGGCAGGAGTGGGTCAGGAGGAAGGTTGCTTGGTGACTCGGGTCTCGGTGGCCTGTTTTGGTGCAATCCCAGTGAACAGTGACACTCTCGAAGTACAGGAGCATCTGGAGACACCTCCGGGCCCTTCTG

Slco2a1-P2A-Cre: TGCCCCTGGGCCTCACCATACCTGTCTCTTCCTGCCTCATAGGTACCTGGGCCTACAG GTAATCTACAAGGTCTTGGGCACACTGCTGCTCTTCTTCATCAGCTGGAGGGTGAAG AAGAACAGGGAATACAGTCTGCAGGAGAATGCTTCCGGATTGATTGGAAGCGGAGC TACTAACTTCTCCCTGTTGAAACAAGCAGGGGATGTCGAAGAGAATCCTGGACCTAT GGCTCCTAAGAAGAAGAGGAAGGTGATGAGCCAGTTCGACATCCTGTGCAAGACTC CTCCAAAGGTGCTGGTGCGGCAGTTCGTGGAGAGATTCGAGAGGCCCAGCGGCGAG AAGATCGCCAGCTGTGCCGCCGAGCTGACCTACCTGTGCTGGATGATCACCCACAAC GGCACCGCCATCAAGAGGGCCACCTTCATGAGCTACAACACCATCATCAGCAACAG CCTGAGCTTCGACATCGTGAACAAGAGCCTGCAGTTCAAGTACAAGACCCAGAAGG CCACCATCCTGGAGGCCAGCCTGAAGAAGCTGATCCCCGCCTGGGAGTTCACCATC ATCCCTTACAACGGCCAGAAGCACCAGAGCGACATCACCGACATCGTGTCCAGCCT GCAGCTGCAGTTCGAGAGCAGCGAGGAGGCCGACAAGGGCAACAGCCACAGCAAG AAGATGCTGAAGGCCCTGCTGTCCGAGGGCGAGAGCATCTGGGAGATCACCGAGAA GATCCTGAACAGCTTCGAGTACACCAGCAGGTTCACCAAGACCAAGACCCTGTACC AGTTCCTGTTCCTGGCCACATTCATCAACTGCGGCAGGTTCAGCGACATCAAGAACG TGGACCCCAAGAGCTTCAAGCTGGTGCAGAACAAGTACCTGGGCGTGATCATTCAG TGCCTGGTGACCGAGACCAAGACAAGCGTGTCCAGGCACATCTACTTTTTCAGCGCC AGAGGCAGGATCGACCCCCTGGTGTACCTGGACGAGTTCCTGAGGAACAGCGAGCC CGTGCTGAAGAGAGTGAACAGGACCGGCAACAGCAGCAGCAACAAGCAGGAGTAC CAGCTGCTGAAGGACAACCTGGTGCGCAGCTACAACAAGGCCCTGAAGAAGAACGC CCCCTACCCCATCTTCGCTATCAAGAACGGCCCTAAGAGCCACATCGGCAGGCACCT GATGACCAGCTTTCTGAGCATGAAGGGCCTGACCGAGCTGACAAACGTGGTGGGCA ACTGGAGCGACAAGAGGGCCTCCGCCGTGGCCAGGACCACCTACACCCACCAGATC ACCGCCATCCCCGACCACTACTTCGCCCTGGTGTCCAGGTACTACGCCTACGACCCC ATCAGCAAGGAGATGATCGCCCTGAAGGACGAGACCAACCCCATCGAGGAGTGGCA GCACATCGAGCAGCTGAAGGGCAGCGCCGAGGGCAGCATCAGATACCCCGCCTGGA ACGGCATCATCAGCCAGGAGGTGCTGGACTACCTGAGCAGCTACATCAACAGGCGG ATCTGACCTTCAGCTGGGACTACTGCCCTGCCCCAGAGACTGGATATCCTACCCCTC CACACCTACCTATATTAACTAATGTTAGCATGCCTTCCTCCTCCTTCC

Transfected cells were cultured and resulting colonies directly screened by PCR for correct integration using the following genotyping primers:

#### Genotyping primers

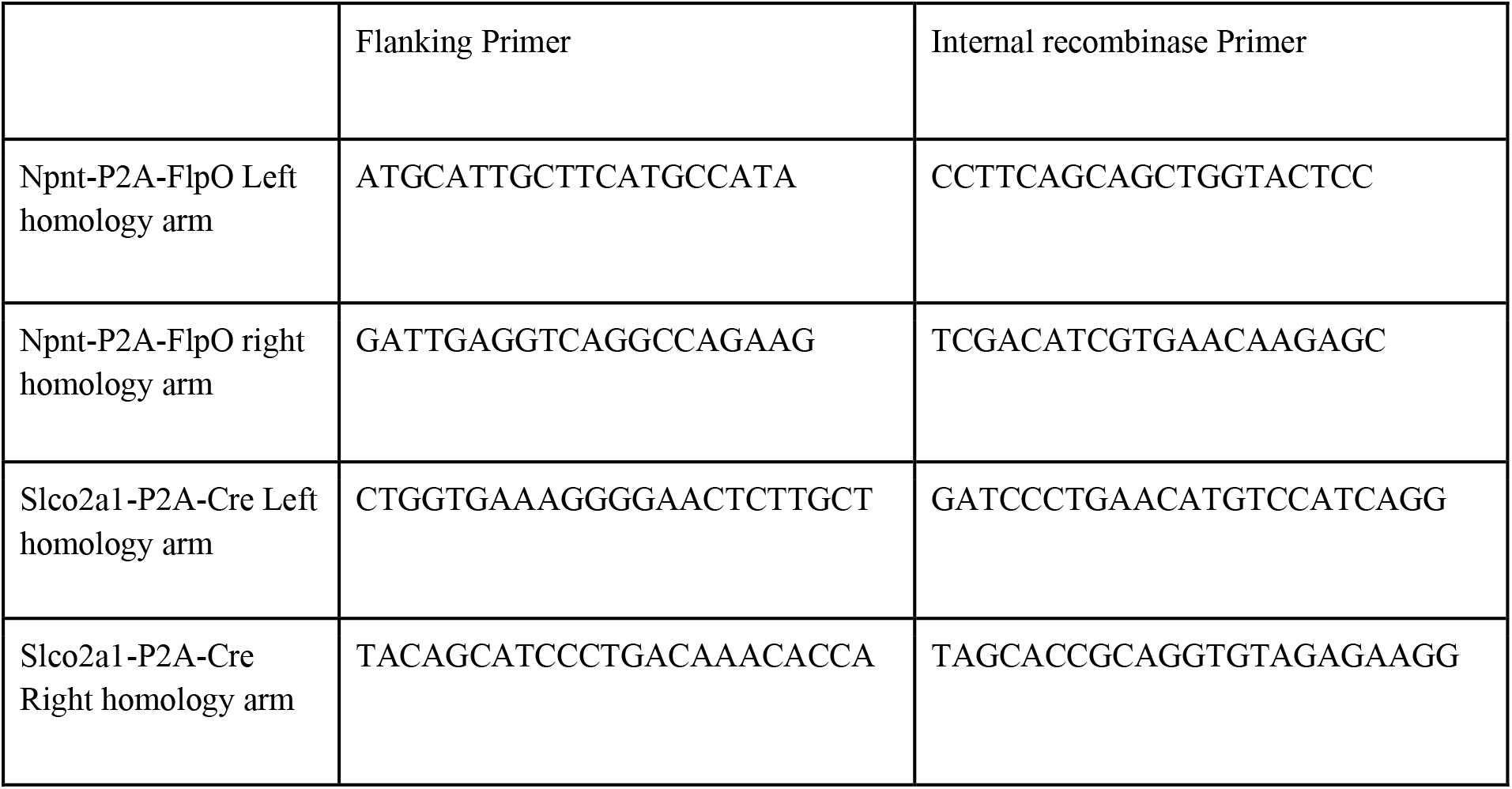

The inserted transgenes were fully sequenced and candidate lines were analyzed for normal karyotype. Lines passing quality control were aggregated with albino morulae and implanted into pseudopregnant females, producing germline-competent chimeric founders which in turn were crossed with the appropriate reporter lines on the C57/BL6 background.

## ACKNOWLEDGEMENTS

We thank additional members of our laboratories and institutions who contributed to the experimental and analytical components of this project. This work was supported by grants from the NIH under: 1U24MH114827, 1U19MH114831, 1U19MH114821, 1U01MH117072, 1U19MH114830, 1U01MH114829, 1U01MH121282, 1U01MH117023, 1U01MH114825, 1U01MH114819, 1U01MH114812, 1U01MH121260, 1U01MH114824, 1U01MH117079, 1U01MH116990, 1U01MH114828, 1R24MH117295, 5R24MH114793, 5R24MH114788, 5R24MH114815. We thank NIH BICCN program officers, in particular Dr. Yong Yao, for their guidance and support throughout this study.

Additional support:

NIH grants R01NS39600 and R01NS86082 to G.A.A. H.S.B. is a Chan Zuckerberg Biohub Investigator. Deutsche Forschungsgemeinschaft through a Heisenberg Professorship (BE5601/4-1), the Cluster of Excellence “Machine Learning --- New Perspectives for Science” (EXC 2064, project number 390727645) and the Collaborative Research Center 1233 “Robust Vision”’ (project number 276693517), the German Federal Ministry of Education and Research (FKZ 01GQ1601 and 01IS18039A) to P.B. This work was supported in part by the Flow Cytometry Core Facility of the Salk Institute with funding from NIH-NCI CCSG: P30 014195 and Shared Instrumentation Grant S10-OD023689. NIH/NIMH R01MH094360 to H.W.D. We thank Marlene Becerra, Tyler Boesen, Chunru Cao, Marina Fayzullina, Kaelan Cotter, Lei Gao, Luis Gacia, Laura Korobkova, Darrick Lo, Christine Mun, Seita Yamashita, Muye Zhu for their technical and informatics support. Howard Hughes Medical Institute for J.E. This work was supported in part by the NNSFC grant 61890953 to H.G. Hearing Health Foundation Hearing Restoration Project grant to R.H. NIH grant OD010425 to G.D.H. This work is supported in part by NIH BRAIN Initiative award RF1MH114126 from the National Institute of Mental Health to E.S.L., J.T.T., and B.P.L. This work was supported in part by the NNSFC grant 81827901 to Q.L. This project was supported in part by NIH grants P51OD010425 from the Office of Research Infrastructure Programs (ORIP) and UL1TR000423 from the National Center for Advancing Translational Sciences (NCATS). Its contents are solely the responsibility of the authors and do not necessarily represent the official view of NIH, ORIP, NCATS, the Institute of Translational Health Sciences at the Washington National Primate Research Center. NSFC grant 61871411 and the University Synergy Innovation Program of Anhui Province GXXT-2019-008 to L.Q. Howard Hughes Medical Institute and the Klarman Cell Observatory for A.R. Howard Hughes Medical Institute for X.Z. V.D.B. is a postdoctoral fellow from the Belgian American Educational Foundation, and is supported by the Research Foundation Flanders (FWO), grant number 1246220N. NSFC Grant No. 32071367 and NSF Shanghai Grant No. 20ZR1420100 to Y.W. NIH grants R01EY023173 and U01MH105982 to H.Z. Researchers from Allen Institute for Brain Science wish to thank the Allen Institute founder, Paul G. Allen, for his vision, encouragement, and support.

## AUTHOR INFORMATION

BRAIN Initiative Cell Census Network (BICCN)

### Corresponding authors

Edward M. Callaway^1^, Hong-Wei Dong^2,†^, Joseph R. Ecker^3,4^, Mike Hawrylycz^5^, Z. Josh Huang^6,†^, Ed S. Lein^5^, John Ngai^7,8,†^, Pavel Osten^6^, Bing Ren^9,10^, Andreas Savas Tolias^11,12^, Owen White^13^, Hongkui Zeng^5^, Xiaowei Zhuang^14^

### BICCN Contributing PI

Giorgio A. Ascoli^15,16^, M. Margarita Behrens^17^, Edward M. Callaway^1^, Jerold Chun^18^, Hong-Wei Dong^2,†^, Joseph R. Ecker^3,4^, Guoping Feng^19–21^, James C. Gee^22^, Satrajit S. Ghosh^23^, Yaroslav O. Halchenko^24^, Mike Hawrylycz^5^, Ronna Hertzano^13,25^, Z. Josh Huang^6,†^, Ed S. Lein^5^, Byungkook Lim^26^, Maryann E. Martone^27,28^, Lydia Ng^5^, John Ngai^7,8,†^, Pavel Osten^6^, Lior Pachter^29^, Alexander J Ropelewski^30^, Timothy L. Tickle^31^, Andreas Savas Tolias^11,12^, Owen White^13^, X. William Yang^32,33^, Hongkui Zeng^5^, Kun Zhang^34^, Xiaowei Zhuang^14^

### Principal manuscript editors

Z. Josh Huang^6,†^, Ed S. Lein^5^, Hongkui Zeng^5^

### Manuscript writing and figure generation

Giorgio A. Ascoli^15,16^, Trygve E. Bakken^5^, Philipp Berens^35,36,37,38^, Edward M. Callaway^1^, Tanya L. Daigle^5^, Hong-Wei Dong^2,†^, Joseph R. Ecker^3,4^, Julie A. Harris^5,†^, Mike Hawrylycz^5^, Z. Josh Huang^6,†^, Nikolas L. Jorstad^5^, Brian E. Kalmbach^5,39^, Dmitry Kobak^35^, Ed S. Lein^5^, Yang Eric Li^10^, Hanqing Liu^3,40^, Katherine S. Matho^6^, Eran A. Mukamel^41^, Maitham Naeemi^5^, John Ngai^7,8,†^, Pavel Osten^6^, Bing Ren^9,10^, Federico Scala^11,12^, Pengcheng Tan^3,42^, Jonathan T. Ting^5,39^, Andreas Savas Tolias^11,12^, Fangming Xie^43^, Hongkui Zeng^5^, Meng Zhang^14^, Zhuzhu Zhang^3^, Jingtian Zhou^3,44^, Xiaowei Zhuang^14^, Brian Zingg^2,†^

### Analysis coordination

Trygve E. Bakken^5^, Edward M. Callaway^1^, Hong-Wei Dong^2,†^, Joseph R. Ecker^3,4^, Julie A. Harris^5,†^, Mike Hawrylycz^5^, Z. Josh Huang^6,†^, Ed S. Lein^5^, Eran A. Mukamel^41^, John Ngai^7,8,†^, Pavel Osten^6^, Bing Ren^9,10^, Andreas Savas Tolias^11,12^, Hongkui Zeng^5^, Xiaowei Zhuang^14^

### Integrated data analysis

Ethan Armand^41^, Trygve E. Bakken^5^, Philipp Berens^35–38^, Hong-Wei Dong^2,†^, Joseph R. Ecker^3,4^, Julie A. Harris^5,†^, Z. Josh Huang^6,†^, Nikolas L. Jorstad^5^, Brian E. Kalmbach^5,39^, Dmitry Kobak^35^, Ed S. Lein^5^, Yang Eric Li^10^, Hanqing Liu^3,40^, Eran A. Mukamel^41^, Pavel Osten^6^, Bing Ren^9,10^, Federico Scala^11,12^, Pengcheng Tan^3,42^, Andreas Savas Tolias^11,12^, Fangming Xie^43^, Zizhen Yao^5^, Hongkui Zeng^5^, Meng Zhang^14^, Zhuzhu Zhang^3^, Jingtian Zhou^3,44^, Xiaowei Zhuang^14^

### Sc/snRNA-seq data generation and processing

Darren Bertagnolli^5^, Tamara Casper^5^, Jerold Chun^18^, Kirsten Crichton^5^, Nick Dee^5^, Dinh Diep^34^, Song-Lin Ding^5^, Weixiu Dong^34^, Elizabeth L. Dougherty^45^, Guoping Feng^19,20,21^, Olivia Fong^5^, Melissa Goldman^46^, Jeff Goldy^5^, Rebecca D. Hodge^5^, Lijuan Hu^47^, C. Dirk Keene^48^, Fenna M. Krienen^46^, Matthew Kroll^5^, Blue B. Lake^34^, Kanan Lathia^5^, Ed S. Lein^5^, Sten Linnarsson^47^, Christine S. Liu^18,49^, Evan Z. Macosko^45^, Steven A. McCarroll^45,46^, Delissa McMillen^5^, Naeem M. Nadaf^45^, Thuc Nghi Nguyen^5,†^, Carter R. Palmer^18,49^, Thanh Pham^5^, Nongluk Plongthongkum^34^, Nora M. Reed^46^, Aviv Regev^45,50,†^, Christine Rimorin^5^, William J. Romanow^18,49^, Steven Savoia^6^, Kimberly Siletti^47^, Kimberly Smith^5^, Josef Sulc^5^, Bosiljka Tasic^5^, Michael Tieu^5^, Amy Torkelson^5^, Herman Tung^5^, Cindy T.J. van Velthoven^5^, Charles R. Vanderburg^45^, Anna Marie Yanny^5^, Hongkui Zeng^5^, Kun Zhang^34^

### ATAC-seq data generation and processing

M. Margarita Behrens^17^, Jerold Chun^18^, Dinh Diep^34^, Weixiu Dong^34^, Rongxin Fang^44^, Xiaomeng Hou^9^, Blue B. Lake^34^, Yang Eric Li^10^, Christine S. Liu^18,49^, Jacinta D. Lucero^17^, Julia K. Osteen^17^, Carter R. Palmer^18,49^, Antonio Pinto-Duarte^17^, Nongluk Plongthongkum^34^, Olivier Poirion^9^, Sebastian Preissl^9^, Bing Ren^9,10^, William J. Romanow^18,49^, Xinxin Wang^9,†^, Kun Zhang^34^

### Methylcytosine data production and analysis

Andrew I. Aldridge^3^, Anna Bartlett^3^, M. Margarita Behrens^17^, Lara Boggeman^51^, Carolyn O’Connor^51^, Rosa G. Castanon^3^, Huaming Chen^3^, Joseph R. Ecker^3,4^, Conor Fitzpatrick^51^, Hanqing Liu^3,40^, Jacinta D. Lucero^17^, Chongyuan Luo^3,4,†^, Joseph R. Nery^3^, Michael Nunn^3^, Julia K. Osteen^17^, Antonio Pinto-Duarte^17^, Angeline C. Rivkin^3^, Wei Tian^3^, Jingtian Zhou^3,44^

### Epi-Retro-Seq data generation and processing

Anna Bartlett^3^, M. Margarita Behrens^17^, Lara Boggeman^51^, Edward M. Callaway^1^, Carolyn O’Connor^51^, Rosa G. Castanon^3^, Bertha Dominguez^52^, Joseph R. Ecker^3,4^, Conor Fitzpatrick^51^, Tony Ito-Cole^1^, Matthew Jacobs^1^, Xin Jin^53^, Cheng-Ta Lee^52^, Kuo-Fen Lee^52^, Paula Assakura Miyazaki^1^, Eran A. Mukamel^41^, Joseph R. Nery^3^, Michael Nunn^3^, Yan Pang^1^, Antonio Pinto-Duarte^17^, Mohammad Rashid^1^, Angeline C. Rivkin^3^, Jared B. Smith^53^, Pengcheng Tan^3,42^, Minh Vu^1^, Elora Williams^53^, Zhuzhu Zhang^3^, Jingtian Zhou^3,44^

### Omics data analysis

Ethan Armand^41^, Trygve E. Bakken^5^, Tommaso Biancalani^45^, A. Sina Booeshaghi^29^, Megan Crow^54^, Dinh Diep^34^, Sandrine Dudoit^55^, Joseph R. Ecker^3,4^, Rongxin Fang^44^, Stephan Fischer^54^, Olivia Fong^5^, Jesse Gillis^54^, Jeff Goldy^5^, Qiwen Hu^56^, Nikolas L. Jorstad^5^, Peter V. Kharchenko^56^, Fenna M. Krienen^46^, Blue B. Lake^34^, Ed S. Lein^5^, Yang Eric Li^10^, Sten Linnarsson^47^, Hanqing Liu^3,40^, Evan Z. Macosko^45^, Eran A. Mukamel^41^, John Ngai^7,8,†^, Sheng-Yong Niu^3^, Vasilis Ntranos^57^, Lior Pachter^29^, Olivier Poirion^9^, Elizabeth Purdom^58^, Aviv Regev^45,50,†^, Davide Risso^59^, Hector Roux de Bézieux^60^, Kimberly Siletti^47^, Kimberly Smith^5^, Saroja Somasundaram^5^, Kelly Street^61^, Valentine Svensson^29^, Bosiljka Tasic^5^, Wei Tian^3^, Eeshit Dhaval Vaishnav^45^, Koen Van den Berge^58,62^, Cindy T.J. van Velthoven^5^, Joshua D. Welch^63^, Fangming Xie^43^, Zizhen Yao^5^, Hongkui Zeng^5^, Jingtian Zhou^3,44^

### Tracing and connectivity data generation

Xu An^6^, Helen S. Bateup^7,8,64^, Ian Bowman^2,†^, Rebecca K. Chance^7^, Hong-Wei Dong^2,†^, Nicholas N. Foster^2,†^, William Galbavy^6^,^66^, Hui Gong^65,67^, Lin Gou^2,†^, Julie A. Harris^5,†^, Joshua T. Hatfield^6^, Houri Hintiryan^2,†^, Karla E. Hirokawa^5,†^, Z. Josh Huang^6,†^, Gukhan Kim^6^, Daniel J. Kramer^7^, Anan Li^65,67^, Xiangning Li^65^, Byungkook Lim^26^, Qingming Luo^67,68^, Katherine S. Matho^6^, Lydia Ng^5^, John Ngai^7,8,†^, Rodrigo Muñoz-Castañeda^6^, David A. Stafford^7^, Hongkui Zeng^5^, Brian Zingg^2,†^

### Morphology data generation and reconstruction

Tanya L. Daigle^5^, Hong-Wei Dong^2,†^, Zhao Feng^65^, Hui Gong^65,67^, Julie A. Harris^5,†^, Karla E. Hirokawa^5,†^, Z. Josh Huang^6,†^, Xueyan Jia^67^, Shengdian Jiang^69^, Tao Jiang^67^, Xiuli Kuang^70^, Rachael Larsen^5^, Phil Lesnar^5^, Anan Li^65,67^, Xiangning Li^65^, Yaoyao Li^70^, Yuanyuan Li^71^, Lijuan Liu^69^, Qingming Luo^67,68^, Hanchuan Peng^69^, Lei Qu^72^, Miao Ren^68^, Zongcai Ruan^69^, Elise Shen^5^, Yuanyuan Song^69^, Wayne Wakeman^5^, Peng Wang^73^, Yimin Wang^74^, Yun Wang^5^, Lulu Yin^69^, Jing Yuan^65,67^, Hongkui Zeng^5^, Sujun Zhao^69^, Xuan Zhao^69^

### OLST/STPT and other data generation

Xu An^6^, William Galbavy^6,66^, Joshua T. Hatfield^6^, Z. Josh Huang^6,†^, Gukhan Kim^6^, Katherine S. Matho^6^, Arun Narasimhan^6^, Pavel Osten^6^, Ramesh Palaniswamy^6^, Rodrigo Muñoz-Castañeda^6^

### Morphology, connectivity and imaging analysis

Xu An^6^, Giorgio A. Ascoli^15,16^, Samik Banerjee^6^, Liya Ding^69^, Hong-Wei Dong^2,†^, Zhao Feng^65^, Nicholas N. Foster^2,†^, William Galbavy^6,66^, Hui Gong^65,67^, Julie A. Harris^5,†^, Joshua T. Hatfield^6^, Z. Josh Huang^6,†^, Bingxing Huo^6^, Xueyan Jia^67^, Gukhan Kim^6^, Hsien-Chi Kuo^5^, Sophie Laturnus^35^, Anan Li^65,67^, Xu Li^6^, Katherine S. Matho^6^, Partha P Mitra^6^, Judith Mizrachi^6^, Maitham Naeemi^5^, Arun Narasimhan^6^, Lydia Ng^5^, Pavel Osten^6^, Ramesh Palaniswamy^6^, Hanchuan Peng^69^, Rodrigo Muñoz-Castañeda^6^, Quanxin Wang^5^, Yimin Wang^74^, Yun Wang^5^, Peng Xie^69^, Feng Xiong^69^, Yang Yu^5^, Hongkui Zeng^5^

### Spatially resolved single-cell transcriptomics (MERFISH)

Hong-Wei Dong^2,†^, Stephen W. Eichhorn^14^, Zizhen Yao^5^, Hongkui Zeng^5^, Meng Zhang^14^, Xiaowei Zhuang^14^, Brian Zingg^2,†^

### Multimodal profiling (Patch-seq)

Philipp Berens^35,36,37,38^, Jim Berg^5^, Matteo Bernabucci^11,12^, Yves Bernaerts^35^, Cathryn René Cadwell^75^, Jesus Ramon Castro^11,12^, Rachel Dalley^5^, Leonard Hartmanis^76^, Gregory D. Horwitz^39,77^, Xiaolong Jiang^11,12,78^, Brian E. Kalmbach^5,39^, C. Dirk Keene^48^, Andrew L. Ko^79,80^, Dmitry Kobak^35^, Sophie Laturnus^35^, Ed S. Lein^5^, Elanine Miranda^11,12^, Shalaka Mulherkar^11,12^, Philip R. Nicovich^5,†^, Scott F. Owen^5,†^, Rickard Sandberg ^76^, Federico Scala^11,12^, Kimberly Smith^5^, Staci A. Sorensen^5^, Zheng Huan Tan^11,12^, Jonathan T. Ting^5,39^, Andreas Savas Tolias^11,12^, Hongkui Zeng^5^

### Transgenic tools

Shona Allen^7^, Xu An^6^, Helen S. Bateup^7,8,64^, Rebecca K. Chance^7^, Tanya L. Daigle^5^, William Galbavy^6,66^, Joshua T. Hatfield^6^, Dirk Hockemeyer^7,64,81^, Z. Josh Huang^6,†^, Gukhan Kim^6^, Rodrigo Muñoz-Castañeda^7^, Angus Y. Lee^82^, Katherine S. Matho^6^, John Ngai^7,8,†^, David A. Stafford^7^, Bosiljka Tasic^5^, Matthew B. Veldman^32,33^, X. William Yang^32,33^, Zizhen Yao^5^, Hongkui Zeng^5^

### NeMO archive and analytics

Ricky S. Adkins^13^, Seth A. Ament^13,83^, Héctor Corrada Bravo^84^, Robert Carter^13^, Apaala Chatterjee^13^, Carlo Colantuoni^13,85^, Jonathan Crabtree^13^, Heather Creasy^13^, Victor Felix^13^, Michelle Giglio^13^, Brian R. Herb^13^, Ronna Hertzano^13,25^, Jayaram Kancherla^84^, Anup Mahurkar^13^, Carrie McCracken^13^, Lance Nickel^13^, Dustin Olley^13^, Joshua Orvis^13^, Michael Schor^13^, Owen White^13^

### Brain Image Library (BIL) archive

Greg Hood^30^, Alexander J Ropelewski^30^

### DANDI archive

Benjamin Dichter^86^, Satrajit S. Ghosh^23^, Michael Grauer^87^, Yaroslav O. Halchenko^24^, Brian Helba^87^

### Brain Cell Data Center (BCDC)

Anita Bandrowski^27^, Nikolaos Barkas^88^, Benjamin Carlin^88^, Florence D. D’Orazi^5^, Kylee Degatano^88^, James C. Gee^22^, Thomas H Gillespie^27^, Mike Hawrylycz^5^, Farzaneh Khajouei^31^, Kishori Konwar^31^, Maryann E. Martone^27,28^, Lydia Ng^5^, Carol Thompson^5^, Timothy L. Tickle^31^

### Project management

Philipp Berens^35,36,37,38^, Florence D. D’Orazi^5^, Hui Gong^65,67^, Houri Hintiryan^2,†^, Kathleen Kelly^6^, Dmitry Kobak^35^, Blue B. Lake^34^, Katherine S. Matho^6^, Stephanie Mok^5^, Michael Nunn^3^, Federico Scala^11,12^, Susan Sunkin^5^, Carol Thompson^5^

### Affiliations

1) Systems Neurobiology Laboratories, The Salk Institute for Biological Studies, La Jolla, CA 92037

2) Center for Integrative Connectomics, USC Mark and Mary Stevens Neuroimaging and Informatics Institute, Department of Neurology, Zilkha Neurogenetic Institute, Keck School of Medicine at USC, University of Southern California, Los Angeles, CA 90033

3) Genomic Analysis Laboratory, The Salk Institute for Biological Studies, La Jolla, CA 92037

4) Howard Hughes Medical Institute, The Salk Institute for Biological Studies, 10010 N. Torrey Pines Road, La Jolla, CA 92037

5) Allen Institute for Brain Science, Seattle, WA, 98109

6) Cold Spring Harbor Laboratory, Cold Spring Harbor, NY 11724

7) Department of Molecular and Cell Biology, University of California, Berkeley, CA 94720

8) Helen Wills Neuroscience Institute, University of California, Berkeley, CA 94720

9) Center for Epigenomics, Department of Cellular and Molecular Medicine, University of California, San Diego School of Medicine, La Jolla, CA, USA

10) Ludwig Institute for Cancer Research, 9500 Gilman Drive, La Jolla, CA 92093, USA

11) Department of Neuroscience, Baylor College of Medicine, Houston, TX, 77030, USA

12) Center for Neuroscience and Artificial Intelligence, Baylor College of Medicine, Houston, TX, 77030, USA

13) Institute for Genome Sciences, University of Maryland School of Medicine, Baltimore, MD

14) Howard Hughes Medical Institute, Department of Chemistry and Chemical Biology, Department of Physics, Harvard University, Cambridge, MA 02138, USA.

15) Center for Neural Informatics, Krasnow Institute for Advanced Study, George Mason University, Fairfax, VA, USA

16) Bioengineering Department, George Mason University, Fairfax, VA, USA.

17) Computational Neurobiology Laboratory, The Salk Institute for Biological Studies, La Jolla, CA 92037

18) Sanford Burnham Prebys Medical Discovery Institute, La Jolla, CA, USA

19) McGovern Institute for Brain Research, Massachusetts Institute of Technology, Cambridge, Massachusetts 02139

20) Department of Brain and Cognitive Sciences, Massachusetts Institute of Technology, Cambridge, Massachusetts 02139

21) Stanley Center for Psychiatric Research, Broad Institute of MIT and Harvard, Cambridge, Massachusetts 02142

22) Department of Radiology, University of Pennsylvania, Philadelphia, PA 19104

23) Massachusetts Institute of Technology, Cambridge, MA

24) Dartmouth College, Hannover, NH

25) Department of Otorhinolaryngology, Anatomy and Neurobiology, University of Maryland School of Medicine, Baltimore, MD

26) Division of Biological Science, Neurobiology Section, University of California, San Diego, La Jolla, CA, USA

27) Department of Neurosciences, University of California, San Diego, La Jolla, CA 92093

28) SciCrunch, Inc.

29) California Institute of Technology, Pasadena, CA 91125

30) Pittsburgh Supercomputing Center, Carnegie Mellon University, 300 South Craig Street, Pittsburgh PA 15213 USA

31) Klarman Cell Observatory and Data Sciences Platform, Broad Institute of MIT and Harvard, Cambridge, MA 02142

32) Semel Institute & Department of Psychiatry and Biobehavioral Science, UCLA, Los Angeles, CA 90095

33) David Geffen School of Medicine, UCLA, Los Angeles, CA 90095

34) Department of Bioengineering, University of California, San Diego, La Jolla, CA 92093

35) Institute for Ophthalmic Research, University of Tübingen, 72076, Germany

36) Center for Integrative Neuroscience, University of Tübingen, 72076, Germany

37) Institute for Bioinformatics and Medical Informatics, University of Tübingen, 72076, Germany

38) Bernstein Center for Computational Neuroscience, University of Tübingen, 72076 Germany

39) Department of Physiology and Biophysics, University of Washington, Seattle, WA 98195

40) Division of Biological Sciences, University of California San Diego, La Jolla, CA 92093

41) Department of Cognitive Science, University of California, San Diego, La Jolla, CA 92037

42) School of Pharmaceutical Sciences, Tsinghua University, Beijing, China, 100084

43) Department of Physics, University of California, San Diego, La Jolla, CA 92037

44) Bioinformatics and Systems Biology Graduate Program, University of California San Diego, La Jolla, CA 92093

45) Broad Institute of MIT and Harvard, Cambridge, MA 02142

46) Department of Genetics, Harvard Medical School, 77 Avenue Louis Pasteur Boston MA 02115

47) Division of Molecular Neurobiology, Department of Medical Biochemistry and Biophysics, Karolinska Institute, S-17177 Stockholm, Sweden

48) Department of Laboratory Medicine and Pathology, University of Washington, Seattle, WA, USA

49) Biomedical Sciences Program, School of Medicine, University of California, San Diego, La Jolla, CA, USA

50) Howard Hughes Medical Institute, Department of Biology, MIT, Cambridge MA 02140

51) Flow Cytometry Core Facility, The Salk Institute for Biological Studies, La Jolla, CA 92037

52) Peptide Biology Laboratories, The Salk Institute for Biological Studies, La Jolla, CA 92037

53) Molecular Neurobiology Laboratory, The Salk Institute for Biological Studies, La Jolla, CA 92037

54) Stanley Institute for Cognitive Genomics, Cold Spring Harbor Laboratory, Cold Spring Harbor, NY 11724

55) Department of Statistics and Division of Biostatistics, University of California, Berkeley, Berkeley, CA 94720-3860

56) Department of Biomedical Informatics, Harvard Medical School, Boston, MA, USA

57) University of California San Francisco, San Francisco, California, 94143,USA

58) Department of Statistics, University of California, Berkeley, Berkeley, CA, USA

59) Department of Statistical Sciences, University of Padova, Italy

60) Division of Biostatistics, School of Public Health, University of California, Berkeley, CA, USA

61) Department of Data Sciences, Dana-Farber Cancer Institute, Boston, MA, 02215

62) Department of Applied Mathematics, Computer Science and Statistics, Ghent University, Ghent, Belgium

63) Department of Computational Medicine and Bioinformatics, University of Michigan, Ann Arbor, MI 48109

64) Chan Zuckerberg Biohub, San Francisco, CA 94158

65) Britton Chance Center for Biomedical Photonics, Wuhan National Laboratory for Optoelectronics, MoE Key Laboratory for Biomedical Photonics, Huazhong University of Science and Technology, Wuhan 430074, China

66) Program in Neuroscience, Department of Neurobiology and Behavior, Stony Brook University, Stony Brook, NY, 11794, USA

67) HUST-Suzhou Institute for Brainsmatics, Suzhou 215123, China

68) School of Biomedical Engineering, Hainan University, Haikou 570228, China

69) SEU-ALLEN Joint Center, Institute for Brain and Intelligence, Southeast University, Nanjing, Jiangsu, China

70) School of Optometry and Ophthalmology, Wenzhou Medical University, Wenzhou, Zhejiang, China

71) Key Laboratory of Intelligent Computation & Signal Processing, Ministry of Education, Anhui University, Hefei, Anhui, China

72) Anhui University, Hefei, Anhui, China

73) Shanghai University, Shanghai, China

74) School of Computer Engineering and Science, Shanghai University, Shanghai, China

75) Department of Pathology, University of California San Francisco, San Francisco, California, 94143,USA

76) Department of Cell and Molecular Biology, Karolinska Institutet, Stockholm, 17177, Sweden

77) Washington National Primate Research Center, University of Washington, Seattle, WA 98195

78) Jan and Dan Ducan Neurological research Institute, Houston,77030, TX, USA

79) Department of Neurological Surgery, University of Washington School of Medicine, Seattle, WA, USA

80) Regional Epilepsy Center at Harborview Medical Center, Seattle, WA, USA

81) Innovative Genomics Institute, University of California, Berkeley, CA, 94720

82) Cancer Research Laboratory, University of California, Berkeley

83) Department of Psychiatry, University of Maryland School of Medicine, Baltimore, MD

84) Center for Bioinformatics and Computational Biology, University of Maryland, College Park, College Park, MD, 20742.

85) Department Neurology & Department Neuroscience, Johns Hopkins School of Medicine, Baltimore, MD

86) CatalystNeuro, LLC

87) Kitware Inc

88) Data Sciences Platform, Broad Institute of MIT and Harvard, Cambridge, MA 02142

^†^Current address for I.B., H.W.D., N.N.F., L.G., H.H, B.Z.: Department of Neurobiology, David Geffen School of Medicine at UCLA, Los Angeles, CA 90095

Current address for J.A.H., K.E.H., P.R.N.: Cajal Neuroscience, 1616 Eastlake Ave E, Suite 208, Seattle, WA 98102

Current address for Z.J.H.: Department of Neurobiology, Duke University School of Medicine, Durham, NC 27705

Current address for C.L.: Department of Human Genetics, University of California, Los Angeles Current address for J.N.: National Institute of Neurological Disorders and Stroke, National Institutes of Health, Bethesda, MD USA

Current address for S.F.O.: Department of Neurosurgery, Stanford University School of Medicine, Stanford, CA 94305

Current address for A.R.: Genentech, 1 DNA Way, South San Francisco, CA

Current address for X.W.: McDonnell Genome Institute, Washington University School of Medicine, St. Louis, MO, USA

## Author contributions

See the consortium author list for full details of author contributions. BICCN Contributing PIs: G.A.A., M.M.B., E.M.C., J.C., H.D., J.R.E., G.F., J.C.G., S.S.G., Y.O.H., M.H., R.H., Z.J.H., E.S.L., B.L., M.E.M., L.N., J.N., P.O., L.P., A.J.R., T.L.T., A.S.T., O.W., X.W.Y., H.Z., K.Z., X.Z. Principal manuscript editors: Z.J.H., E.S.L., H.Z. Manuscript writing and figure generation: G.A.A., T.E.B., P.B., E.M.C., T.L.D., H.D., J.R.E., J.A.H., M.H., Z.J.H., N.L.J., B.E.K., D.K., E.S.L., Y.E.L., H.L., K.S.M., E.A.M., M.N., J.N., P.O., B.R., F.S., P.T., J.T.T., A.S.T., F.X., H.Z., M.Z., Z.Z., J.Z., X.Z., B.Z. Analysis coordination: T.E.B., E.M.C., H.D., J.R.E., J.A.H., M.H., Z.J.H., E.S.L., E.A.M., J.N., P.O., B.R., A.S.T., H.Z., X.Z. Integrated data analysis: E.A., T.E.B., P.B., H.D., J.R.E., J.A.H., Z.J.H., N.L.J., B.E.K., D.K., E.S.L., Y.E.L., H.L., E.A.M., P.O., B.R., F.S., P.T., A.S.T., F.X., Z.Y., H.Z., M.Z., Z.Z., J.Z., X.Z. Sc/snRNA-seq data generation and processing: D.B., T.C., J.C., K.C., N.D., D.D., S.D., W.D., E.L.D., G.F., O.F., M.G., J.G., R.D.H., L.H., C.D.K., F.M.K., M.K., B.B.L., K.L., E.S.L., S.L., C.S.L., E.Z.M., S.A.M., D.M., N.M.N., T.N.N., C.R.P., T.P., N.P., N.M.R., A.R., C.R., W.J.R., S.S., K.S., K.S., J.S., B.T., M.T., A.T., H.T., C.T.V.V., C.R.V., A.M.Y., H.Z., K.Z. ATAC-seq data generation and processing: M.M.B., J.C., D.D., W.D., R.F., X.H., B.B.L., Y.E.L., C.S.L., J.D.L., J.K.O., C.R.P., A.P., N.P., O.P., S.P., B.R., W.J.R., X.W., K.Z. Methylcytosine data production and analysis: A.I.A., A.B., M.M.B., L.B., C.O., R.G.C., H.C., J.R.E., C.F., H.L., J.D.L., C.L., J.R.N., M.N., J.K.O., A.P., A.C.R., W.T., J.Z. Epi-Retro-Seq data generation and processing: A.B., M.M.B., L.B., E.M.C., C.O., R.G.C., B.D., J.R.E., C.F., T.I., M.J., X.J., C.L., K.L., P.A.M., E.A.M., J.R.N., M.N., Y.P., A.P., M.R., A.C.R., J.B.S., P.T., M.V., E.W., Z.Z., J.Z. Omics data analysis: E.A., T.E.B., T.B., A.S.B., M.C., D.D., S.D., J.R.E., R.F., S.F., O.F., J.G., J.G., Q.H., N.L.J., P.V.K., F.M.K., B.B.L., E.S.L., Y.E.L., S.L., H.L., E.Z.M., E.A.M., J.N., S.N., V.N., L.P., O.P., E.P., A.R., D.R., H.R.D.B., K.S., K.S., S.S., K.S., V.S., B.T., W.T., E.D.V., K.V.D.B., C.T.V.V., J.D.W., F.X., Z.Y., H.Z., J.Z. Tracing and connectivity data generation: X.A., H.S.B., I.B., R.K.C., H.D., N.N.F., W.G., H.G., L.G., J.A.H., J.T.H., H.H., K.E.H., Z.J.H., G.K., D.J.K., A.L., X.L., B.L., Q.L., K.S.M., L.N., J.N., R.M., D.A.S., H.Z., B.Z. Morphology data generation and reconstruction: T.L.D., H.D., Z.F., H.G., J.A.H., K.E.H., Z.J.H., X.J., S.J., T.J., X.K., R.L., P.L., X.L., Y.L., Y.L., L.L., Q.L., H.P., L.Q., M.R., Z.R., E.S., Y.S., W.W., P.W., Y.W., Y.W., L.Y., J.Y., H.Z., S.Z., X.Z. OLST/STPT and other data generation: X.A., W.G., J.T.H., Z.J.H., G.K., K.S.M., A.N., P.O., R.P., R.M. Morphology, connectivity and imaging analysis: X.A., G.A.A., S.B., L.D., H.D., Z.F., N.N.F., W.G., H.G., J.A.H., J.T.H., Z.J.H., B.H., X.J., G.K., H.K., S.L., A.L., X.L., K.S.M., P.P.M., J.M., M.N., A.N., L.N., P.O., R.P., H.P., R.M., Q.W., Y.W., Y.W., P.X., F.X., Y.Y., H.Z. Spatially resolved single-cell transcriptomics (MERFISH): H.D., S.W.E., Z.Y., H.Z., M.Z., X.Z., B.Z. Multimodal profiling (PATCH-seq): P.B., J.B., M.B., Y.B., C.R.C., J.R.C., R.D., L.H., G.D.H., X.J., B.E.K., C.D.K., A.L.K., D.K., S.L., E.S.L., E.M., S.M., P.R.N., S.F.O., R.S., F.S., K.S., S.A.S., Z.H.T., J.T.T., A.S.T., H.Z. Transgenic tools: S.A., X.A., H.S.B., R.K.C., T.L.D., W.G., J.T.H., D.H., Z.J.H., G.K., D.J.K., A.Y.L., K.S.M., J.N., D.A.S., B.T., M.B.V., X.W.Y., Z.Y., H.Z. NeMO archive and analytics: R.S.A., S.A.A., H.C.B., R.C., A.C., C.C., J.C., H.C., V.F., M.G., B.R.H., R.H., J.K., A.M., C.M., L.N., D.O., J.O., M.S., O.W. Brain Image Library (BIL) archive: G.H., A.J.R. DANDI archive: B.D., S.S.G., M.G., Y.O.H., B.H. Brain Cell Data Center (BCDC): A.B., N.B., B.C., F.D.D., K.D., J.C.G., T.H.G., M.H., F.K., K.K., M.E.M., L.N., C.T., T.L.T. Project management: P.B., F.D.D., H.G., H.H., K.K., D.K., B.B.L., K.S.M., S.M., M.N., F.S., S.S., C.T.

## Competing interests

A.B. is a cofounder of SciCrunch, a company devoted to improving scientific communication. J.R.E. is a member of Zymo Research SAB. J.A.H. is currently employed by Cajal Neuroscience. K.E.H. is currently employed by Cajal Neuroscience. P.V.K. serves on the Scientific Advisory Board to Celsius Therapeutics Inc. M.E.M. is a founder and CSO of SciCrunch Inc., a UCSD tech start up that produces tools in support of reproducibility including RRIDs. P.R.N. is currently employed by Cajal Neuroscience. A.R. is a founder and equity holder of Celsius Therapeutics, an equity holder in Immunitas Therapeutics and until August 31, 2020 was an SAB member of Syros Pharmaceuticals, Neogene Therapeutics, Asimov and ThermoFisher Scientific. From August 1, 2020, A.R. is an employee of Genentech. B.R. is shareholder of Arima Genomics, Inc. and Epigenome Technologies, Inc. K.Z is a co-founder, equity holder and serves on the Scientific Advisor Board of Singlera Genomics. X.Z. is a co-founder and consultant of Vizgen.

---

**Extended Data Figure 1.**
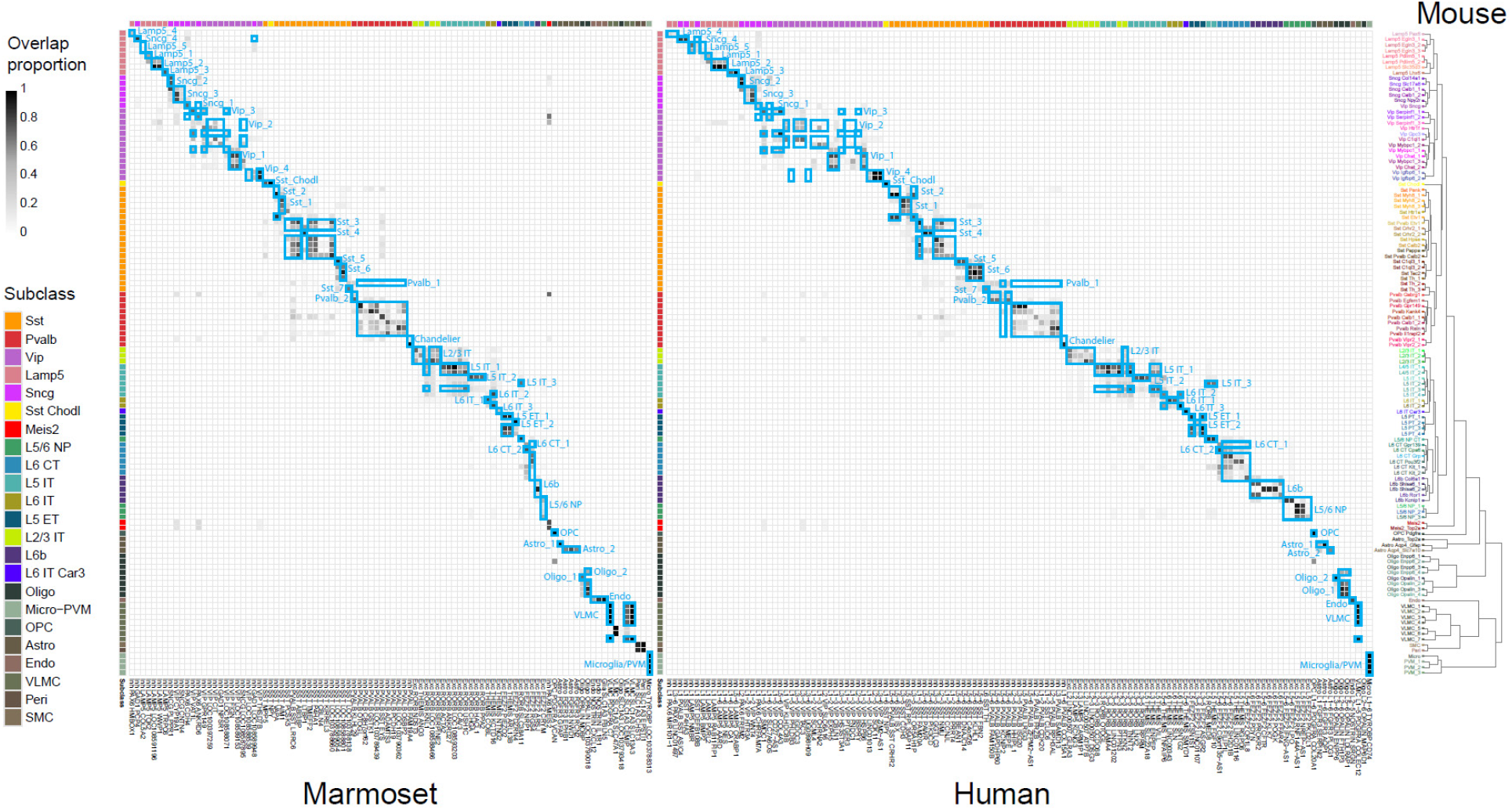
Cluster overlap heatmap showing the proportion of nuclei in each pair of species clusters that are mixed in the cross-species integrated space. Cross-species consensus clusters are indicated by labeled blue boxes. Mouse clusters (rows) are ordered by the mouse MOp transcriptomic taxonomy dendrogram reproduced from ^45^. Marmoset (left columns) and human (right columns) transcriptomic clusters (reproduced from ^48^ are ordered to align with mouse clusters. Color bars at top and left indicate subclasses of within-species clusters.

**Extended Data Figure 2.**
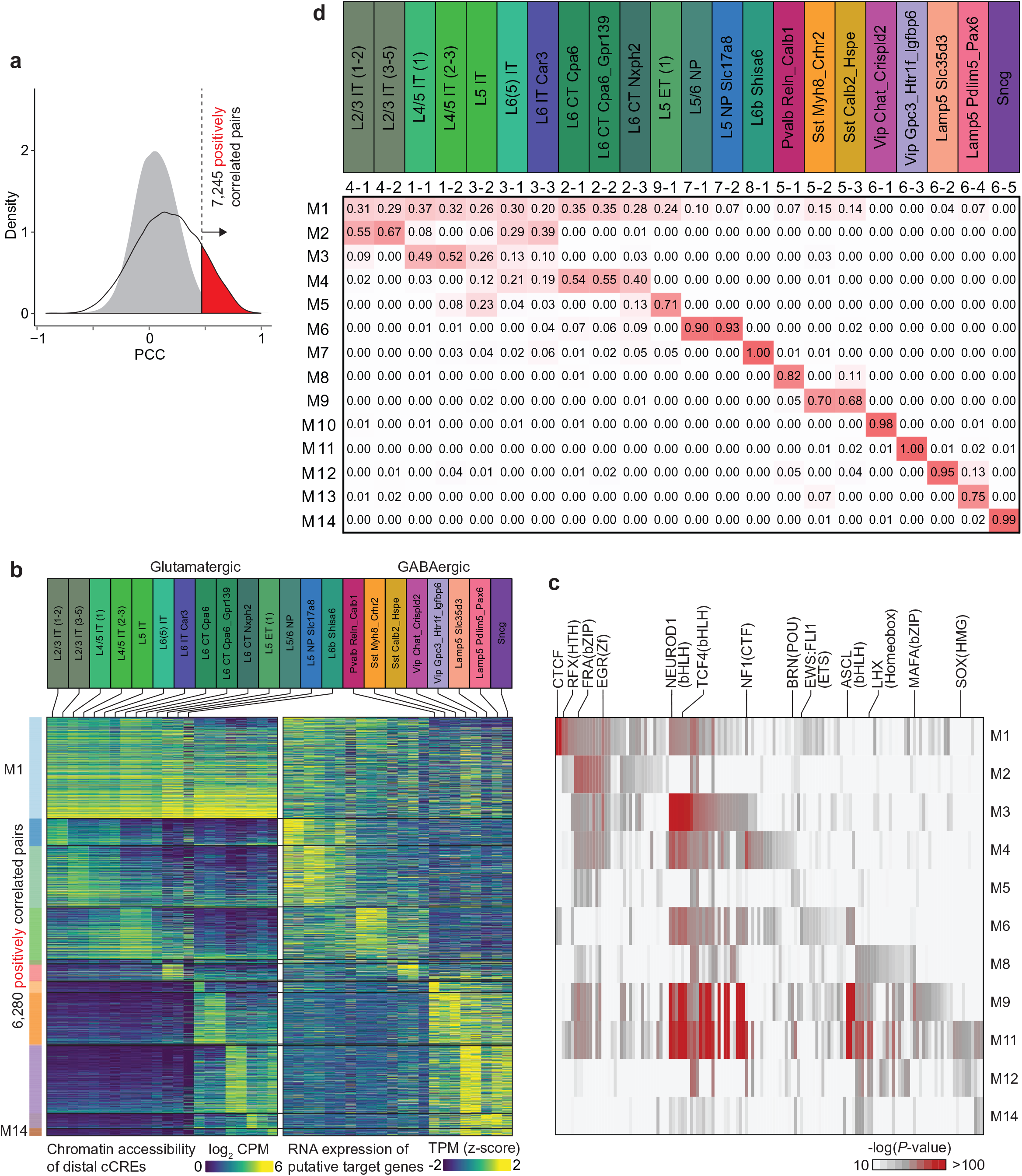
Characterization of putative enhancer-gene pairs. **a,** Detection of putative enhancer-gene pairs. 7,245 pairs of positively correlated cCRE and genes (highlighted in red) were identified using an empirically defined significance threshold of FDR<0.01. Grey filled curve shows the distribution of PCC for randomly shuffled cCRE-gene pairs. **b,** Heatmap of chromatin accessibility of 6,280 putative enhancers across joint cell clusters (left) and expression of 2,490 target genes (right). Note genes are displayed for each putative enhancer separately. CPM: counts per million, TPM: transcripts per million. **c,** Enrichment of known transcription factor motifs in distinct enhancer-gene modules. Displayed are known motifs from HOMER with enrichment -log p-value >5. In module M1, de novo motif analysis of putative enhancers in this module showed enrichment of sequence motif recognized by transcription factors CTCF, MEF2. CTCF is a widely expressed DNA binding protein with a well-established role in transcriptional insulation and chromatin organization, but recently it was also reported that CTCF can promote neurogenesis by binding to promoters and enhancers of related genes. In the L2/3 IT selective module M2, the putative enhancers were enriched for the binding motif for Zinc-finger transcription factor EGR, a known master transcriptional regulator of excitatory neurons ^97^. In the Pvalb selective module M8, the putative enhancers were enriched for sequence motifs recognized by the MADS factor MEF2, which is associated with regulating cortical inhibitory and excitatory synapses and behaviors relevant to neurodevelopmental disorders ^98^. **d,** Heatmap shows the weights of each joint cell cluster in each module, which were derived from the coefficient matrix. The values of each column are scaled (0–1).

**Extended Data Figure 3.**
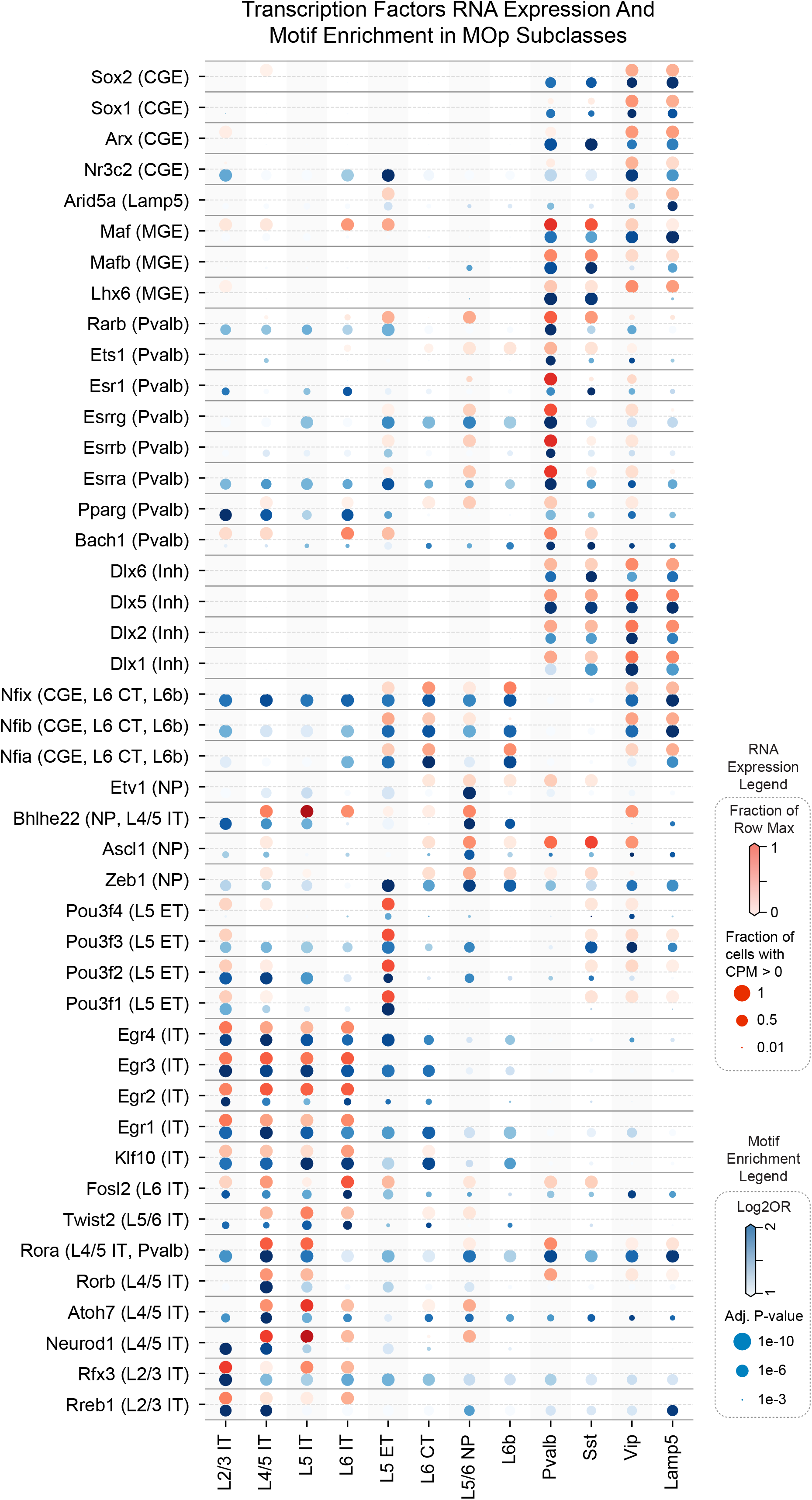
Dot plot illustrating the RNA expression levels (red) and hypo-CG-DMR motif enrichments (blue) of transcription factors (TFs) in mouse MOp subclasses. The size and color of red dots indicate the proportion of expressing cells and the average expression level in each subclass, respectively. The size and color of blue dots indicate adjusted P-value and log2(Odds Ratio) of motif enrichment analysis, respectively.

## Notes

### Competing Interest Statement

The competing interests are detailed in the Competing Interests section in the manuscript file.

